# Comprehensive Phylogenomic Inference of Eucalypts Reveals Taxonomic Limits through Gene Tree Concordance

**DOI:** 10.64898/2026.09.25.753384

**Authors:** Jeremias Ivan, Bruce S. Martin, Kevin Murray, Anne-Cécile Colin, Carsten Kuelheim, Rose Andrew, Dean Nicolle, Zander A. Myburg, James Leebens-Mack, Ashley Jones, Robert Lanfear, Justin O. Borevitz

**Affiliations:** Research School of Biology, Australian National University, ACT 2601, Australia; Department of Plant Biology, University of Georgia, GA 30602, USA; Max Planck Institute for Biology Tübingen, 72076 Tübingen, Germany; Hawkesbury Institute for the Environment, Western Sydney University, NSW 2751, Australia; College of Forest Resource and Environmental Science, Michigan Technological University, MI 49931, USA; School of Environmental and Rural Science, University of New England, NSW 2351, Australia; Currency Creek Arboretum, PO Box 808 Melrose Park, SA 5039, Australia; Department of Genetics, Stellenbosch University, Stellenbosch 7600, South Africa

**Keywords:** *incomplete lineage sorting*, *hybridisation*, *BUSCO*, *concordance factors*, *taxonomic consistency test*

## Abstract

Speciation can be a slow process for long-lived organisms with large populations due to persistent incomplete lineage sorting, while hybridisation can further obscure species boundaries through admixture. In eucalypts, there is tremendous morphological and environmental variation with almost a thousand described species, subspecies, and hybrid taxa falling into multiple subgenera and intermediate sections. Multilocus molecular studies have generated concatenated species trees supporting deeper divergence between the *Corymbia + Blakella* + *Angophora* clade with the rest of eucalypts. However, the proportion of gene trees that recover the concatenated tree topology, including the taxonomic relationships among individual subgenera and sections, remains underappreciated. In this study, we assessed the predictive power of current eucalypt taxonomy - represented as a concatenated tree - in explaining topological variation across gene trees. To do this, we first extracted 1,187 BUSCO loci from short-read data of 701 samples representing roughly 500 described species using CAPTUS. We then inferred individual gene trees and a concatenated species tree using IQ-TREE2. Finally, we calculated the concordance factors for every described subgenus and section on the concatenated tree, and developed a taxonomic consistency test that tracks the taxonomic placement of each sample across gene trees. We show that, even though most taxonomic groups are monophyletic on the concatenated tree with 100% bootstrap support, several groups exhibit very low concordance, indicating that most gene trees do not recover their relationships as represented by the concatenated tree. These results highlight important considerations for current eucalypt taxonomic classification and provide a phylogenomic inference pipeline that can be applied not only to future eucalypt studies, but also to other evolutionarily complex groups of taxa.

## Introduction

Eucalypts (Myrtaceae: *Eucalyptus* L’Hér *sensu lato*, including (sub)genera *Corymbia* K.D.Hill & L.A.S.Johnson, *Blakella* (L.D.Pryor & L.A.S.Johnson ex Brooker) Crisp & L.G.Cook, and *Angophora* Cav.) are a key group of native Australian plants consisting of over 800 described morphological species that have long posed challenges for phylogenetic inference (Steane et al. 2011; Bayly et al. 2013; Jones et al. 2016; Schuster et al. 2018; Thornhill et al. 2019; Crisp et al. 2024; Orel et al. 2026). Recurring bouts of rapid speciation across the group, combined with large effective population sizes (*N_e_*) (Silva-Junior and Grattapaglia 2015), allow ancestral polymorphisms to persist in eucalypt lineages for many thousands of generations (roughly 4*N_e_* generations), resulting in prolonged incomplete lineage sorting (ILS) (Pamilo and Nei 1988; Pease and Hahn 2013; Smith et al. 2026). Additionally, eucalypts have long been known to hybridise in both natural and controlled settings (Pryor and Johnson 1981; Griffin et al. 1988; Potts and Wiltshire 1997), with modern genetic studies suggesting that complete reproductive isolation may take tens of millions of years to fully develop among diverging lineages (Larcombe et al. 2015; Rutherford et al. 2018; Robins et al. 2021; McLay et al. 2023; Orel et al. 2026). Between widespread ILS and gene flow, the histories of different genetic loci (i.e., gene trees) vary tremendously across eucalypt genomes (McLay et al. 2023; Orel et al. 2026), such that any single bifurcating “species tree” of the group is at best a point estimate of a much more complex set of evolutionary histories. Despite this, the molecular basis for the current taxonomic classification of eucalypts was mainly derived from statistical support for particular clades in a single species tree inferred by concatenating multiple loci (e.g., Crisp et al. 2024), which does not explicitly account for variation in gene trees across genomes (Degnan and Rosenberg 2006, 2009).

Here, we emphasise that the differences among gene trees and the degree of gene tree discordance (i.e., proportion of gene trees that do not agree with the concatenated species tree topology) can have practical taxonomic implications. For instance, if the separation between two clades is broadly recovered across gene trees (i.e., the separation has high concordance), one might argue that the two clades are genuinely two distinct lineages since the lineage designations apply to the majority of the genome. If, however, most gene trees fail to recover clear separation between the two clades (i.e., the separation has low concordance), one might argue against formalising the split since it applies to a relatively small fraction of the genome. Importantly, this measure of genomic support can be applied not only for higher taxonomic ranks like genera – a subject of intense research and ongoing debate in eucalypts (Crisp et al. 2024; Cook et al. 2025; Nicolle et al. 2025) – but also infrageneric ranks like subgenera and sections, which remain relatively underappreciated among many eucalypt researchers. In other words, it is useful to measure the degree of gene tree discordance across individual taxonomic ranks / groups, and concordance factors offer a straightforward way to do this (Lanfear and Hahn 2024).

In general, there are three measures of concordance factors (CFs; Lanfear and Hahn 2024): gene concordance factors (gCF), quartet concordance factors (qCF), and site concordance factors (sCF). gCF represents the proportion of gene trees that recover individual branches on a given species tree topology. As a result, gCF offers the most complete view of local topological variation, but is also sensitive to gene-tree estimation error (e.g., due to lack of phylogenetic signal in individual loci) (Lanfear and Hahn 2024). By comparison, qCF (which works on quartets derived from gene trees) and sCF (which calculates concordance at sites that have information about each particular branch on the tree) both sidestep the issue of gene-tree estimation error in different ways by assuming that there are only three possible resolutions of each branch in the species tree (Lanfear and Hahn 2024). In contrast to branch statistical supports such as bootstraps (Felsenstein 1985) that ask the question, “how robust is each branch on the tree to sampling variance,” CFs ask “how consistently is the given branch recovered across individual loci or sites”. From this perspective, CFs are analogous to standard deviation, measuring topological variation across the genome, while statistical support measures are more similar to standard error, measuring the certainty that a particular topology is best-supported among alternatives (Lanfear and Hahn 2024). Accordingly, bootstrap support tends to converge to 100% as the size of the dataset increases, while CF estimates become more accurate and show no such monotonic change (Lanfear and Hahn 2024). For example, Crisp et al. (2024) showed that all clades separating the (sub)genera *Angophora*, *Blakella*, and *Corymbia* have 100% bootstrap support according to their concatenated species tree analysis. However, out of 101 loci, the clade comprising *Corymbia* and *Angophora* only had 17% gCF, while *Blakella* had 21.6% gCF (Crisp et al. 2024). In other words, even though these clades were consistently recovered from resampled datasets (i.e., 100% bootstrap support), only one in roughly five gene trees actually recovered the exact same clades. This highlights the importance of not only looking at a single concatenated species tree and its statistical support, but also at individual gene tree topologies inferred from different genomic regions.

In this study, we aim to assess the predictive power of current eucalypt taxonomic classification (which is represented as a species tree) in describing topological variation in individual gene trees. We go well beyond previous studies by: (i) substantially increasing the number of loci and samples analysed; and (ii) calculating CFs for all nodes spanning individual (sub)genera and sections on a newly inferred species tree. To do this, we first extract 1,187 BUSCO loci (Simão et al. 2015; Manni et al. 2021) from whole genome short-read data of 701 independent eucalypt samples representing approximately 500 described species (Nicolle 2024) using CAPTUS (Ortiz et al. 2026). We then infer individual gene trees for each locus, and concatenate the locus alignments to build the species tree using IQ-TREE2 (Minh et al. 2020b). We finally calculate the gCF, sCF, and qCF of individual branches on the species tree using IQ-TREE2 (Minh et al. 2020a) and ASTRAL-IV (Zhang et al. 2025), and develop a taxonomic consistency test that tracks the taxonomic placement of each sample across gene trees based on their nearest neighbours on each gene tree. Throughout this study, we use the one-genus classification system from Nicolle (2024) for two main reasons: (i) it better standardises subgeneric ranks among eucalypts, as genetic divergence among *Corymbia*, *Blakella*, and *Angophora* is roughly equivalent to that among subgenera in the rest of eucalypts (which we refer to as *Eucalyptus sensu stricto*), and (ii) all relevant taxonomic information under this system is readily accessible as a regularly updated and versioned spreadsheet including synonyms, thereby ensuring the taxonomic interoperability of this work for the foreseeable future. Nevertheless, we fully acknowledge ongoing debates regarding the generic circumscription of eucalypts (the authors of this paper alone hold opposing viewpoints on the topic; see Nicolle et al. (2025) and Cook et al. (2025)). Overall, we present a phylogenomic pipeline to assess gene-tree-based genomic support for individual taxonomic groupings, and perform the most comprehensive phylogenomic analyses of eucalypts to date.

## Materials & Methods

### Overview

Our dataset comprised 990 eucalypt samples from Currency Creek Arboretum (CCA; Fig. 1), as well as one *E. melliodora* A.Cunn. ex Schauer sample from The Australian National University (ANU). For each sample, we collected their leaf tissues from CCA and ANU, respectively, and extracted their DNA in batches of 96-well plates. We generated whole-genome short-read libraries and sequences of 190 samples at ANU, and sent the remaining samples to Joint Genome Institute (JGI) for library and sequencing as part of a community sequencing program grant to Myburg, Borevitz, and Wegrzyn. We then processed each set of short-reads through CAPTUS (Ortiz et al. 2026) to extract 1,187 single-copy, complete BUSCO loci (Simão et al. 2015) shared between the 36 reference genomes from (Ferguson et al. 2024b). We also incorporated the short-read data from Ferguson et al. (2024b) to assess the clustering of samples that come from the same species on the species tree. These steps filtered out three failed CCA samples with zero loci recovered, resulting in 1,187 BUSCO alignments with up to 1,024 samples (987 CCA + two replicates of the *E. melliodora* sample from ANU + 35 samples from Ferguson et al. (2024b)) per alignment.

**Figure 1.**
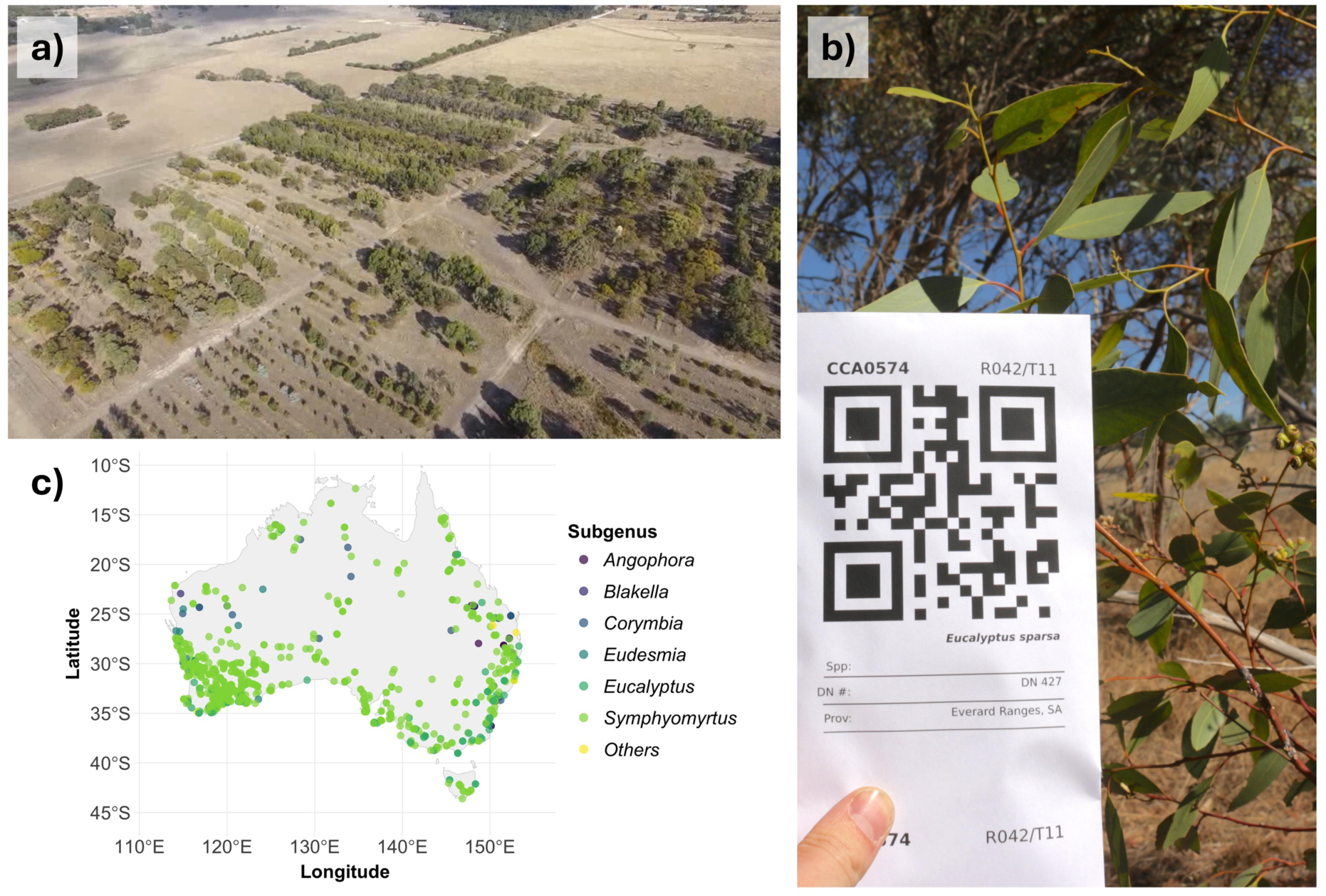
Eucalypt samples from Currency Creek Arboretum (CCA). a) 100 m above ground level image of CCA; b) Example of a tree sample from *Eucalyptus sparsa*; c) Maternal tree seed collection sites for 912 CCA progeny samples used in this study (excluding 78 samples with unavailable coordinates).

For each BUSCO locus, we inferred individual gene trees using IQ-TREE2 (Minh et al. 2020b) and used TreeShrink (Mai and Mirarab 2018) to trim outlier long branches. In order to identify mislabeled outliers and putative contaminated samples, we then inferred a concatenated tree from all samples, as well as developing a taxonomic consistency test that tracks the position of each sample across gene trees (see *Running taxonomic consistency test* section below). These steps filtered out 323 samples, resulting in a final dataset of 701 CCA samples. We re-inferred individual gene trees and the concatenated tree using the subsetted alignments, reran the taxonomic consistency test, and calculated concordance factors (CFs) for every branch on the concatenated tree. Figure 2 shows the overview of this workflow, with details of individual steps provided in the following paragraphs. The code to run this pipeline is available at https://github.com/jeremiasivan/EucsPhylogenomics (DOI: 10.5281/zenodo.22875230).

**Figure 2.**
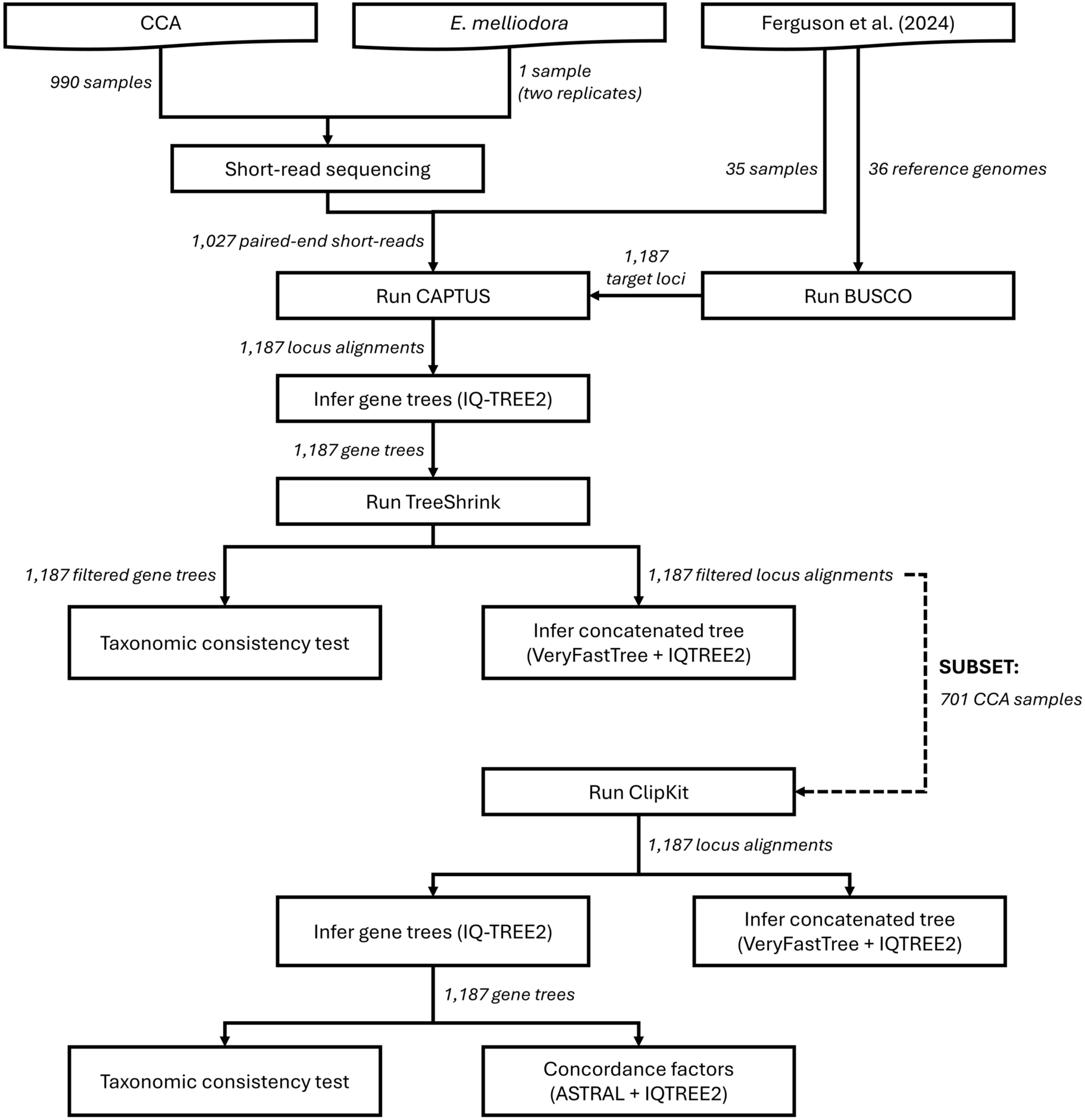
Overview of methods in this study. The workflow includes initial analyses of 1,027 eucalypt samples from three different sources, and the final tree inference of the subsetted 701 CCA samples. Please refer to the *Detailed Methods* section for details of individual steps.

## Detailed Methods

### Sample collection and sequencing

#### Collecting leaf tissues and sequencing of CCA + E. melliodora samples

Leaf tissue was collected from 990 eucalypt samples at Currency Creek Arboretum, Currency Creek, South Australia, Australia, and silica-dried in envelopes for storage. We also collected leaf tissue of an *E. melliodora* individual from The Australian National University, Canberra, Australia (which we labelled as *melliodora_Csiro* as it grew on the walking path up to The Commonwealth Scientific and Industrial Research Organisation (CSIRO)). For DNA extraction, approximately twenty 3 mm leaf discs of each sample were excised using a Qiagen Uni-Core (WB100078, Qiagen, Venlo, Netherlands) and placed into individual wells of 96- well, 1.1 mL tube array (Axygen Scientific, Union City, CA, USA, MTS-11-8-C-R-10 and MTS-8CP-C-1), each pre-loaded with a single 3 mm ball bearing. Tissues were flash-frozen in liquid nitrogen, then disrupted on a TissueLyser II (Qiagen), and genomic DNA was extracted using an Invisorb DNA Plant HTS 96-well plate kit (Stratec Molecular 7037300400).

Whole-genome short-read libraries for the first two 96-well plates were generated using our optimised transposase method (Jones et al. 2023), which adapts the Illumina DNA Prep (M) Tagmentation kit (24 Samples, IPB; Cat# 20060060) and its bead-linked transposase (BLT; Illumina document# 1000000033561 v05). Tagmented DNA underwent 14 cycles of PCR with custom index primers, after which the pooled libraries were size-selected on a PippinHT (Sage Science) to retain 300-500 bp inserts.

An initial 190 samples (189 CCA + *melliodora_Csiro* sample) were sequenced on a NovaSeq 6000 (Illumina) using an S4 flow cell with a 300-cycle kit (150 bp paired-end) at the Biomolecular Resource Facility, Australian National University, Canberra, Australia. DNA from the remaining samples was sent to the Joint Genome Institute, United States Department of Energy, Berkeley, California, USA, where equivalent short-read libraries were constructed and sequenced on a NovaSeq 6000 (Illumina) under proposal ID: 508017. Note that we sequenced the *melliodora_Csiro* sample twice, resulting in two replicates of short-read data denoted as *melliodora_1_Csiro* and *melliodora_2_Csiro*.

#### Pre-processing short-read datasets

For each set of short-reads from the 990 CCA samples and the *melliodora_Csiro* sample, we separated the forward and reverse reads using `reformat.sh` from BBTools v39.10 (Bushnell 2014) following CAPTUS requirements. In addition, we retrieved the short-read data of the 35 eucalypt samples from Ferguson et al. (2024b) that had been adaptor-trimmed using AdapterRemoval v2 (Schubert et al. 2016). As a result, our final short-read dataset consisted of 1,027 paired-end samples.

### Extraction of BUSCO loci and initial tree inference

#### Extracting target BUSCO loci from 36 eucalypt reference genomes

We ran the BUSCO pipeline (Simão et al. 2015; Manni et al. 2021) on each of the 36 eucalypt reference genomes from Ferguson et al. (2024b) and extracted all loci that were identified as single-copy and complete across all reference genomes, totaling 2,310 loci. For each locus, we filtered out poorly aligned sequences and/or columns based on automatic filtering by TAPER (Zhang et al. 2021) and identification of gappy sequences / columns by Biostrings package (Pagès et al. 2024) in R (R Core Team 2023), as well as manual curation by eye. Lastly, we kept all BUSCO loci with at least one representative species from the subgenera *Corymbia*, *Blakella*, or *Angophora* (forming a well-supported monophyletic “CBA” clade) and 17 from the rest of the *Eucalyptus s.s.* reference genomes, resulting in 1,187 retained loci. Details of this step are provided in Supplementary Text.

#### Extracting target BUSCO loci from short-read datasets using CAPTUS

We extracted the 1,187 reference BUSCO loci from the short-read data of the 1,027 samples using CAPTUS v1.6.5 (Ortiz et al. 2026) with default parameters, setting the target loci as miscellaneous DNA markers. CAPTUS started with cleaning the short-reads using `bbduk.sh` from BBTools (Bushnell 2014), followed by *de novo* assembly using MEGAHIT (Li et al. 2015). BUSCO loci were then extracted from each assembly using BLAT (Kent 2002). For each sample, we extracted BUSCO loci using only target sequences from reference genomes belonging to the same major clade (i.e., CBA or *Eucalyptus s.s.*) to reduce the impact of reference bias on phylogenetic inference (Bertels et al. 2014; Valiente-Mullor et al. 2021; Ivan and Lanfear 2026a). The pipeline then aligned individual BUSCO alignments using MAFFT (Katoh et al. 2002; Katoh and Standley 2013) excluding paralogs, masked putative alignment errors using TAPER (Zhang et al. 2021), and removed gappy columns / exceptionally short sequences using ClipKIT (Steenwyk et al. 2020). For each BUSCO locus, we then removed sequences with ≥50% gaps and subsequently columns with ≥50% gaps using the Biostrings package in R. This step filtered out three CCA samples with zero loci recovered, resulting in 1,024 samples.

#### Inferring individual gene trees and initial concatenated tree

We inferred individual gene trees for each BUSCO locus using IQ-TREE v2.4.0 (Minh et al. 2020b) with the best nucleotide models from ModelFinder (Kalyaanamoorthy et al. 2017). We then used TreeShrink v1.3.9 (Mai and Mirarab 2018) to remove taxa on long branches from each gene tree and its corresponding BUSCO alignment, resulting in a dataset with 884 to 1,021 (out of 1,024 possible) samples per alignment. We then re-estimated the best substitution model for each post-TreeShrink BUSCO alignment using ModelFinder. We concatenated the 1,187 BUSCO alignments using AMAS (Borowiec 2016) and converted all question marks (denoting missing data) to gaps. Lastly, we inferred a concatenated tree using VeryFastTree v4.0.5 (Piñeiro et al. 2020) under a GTR model, and then refined the resulting tree using IQ-TREE2 with the following commands:

~~~
iqtree2 -t veryfasttree_tree.tre -s concatenated_alignment.fna
-p locus_partition.nex
~~~

where `-t veryfasttree_tree.tre` initiates the IQ-TREE2 tree search with the VeryFastTree tree, `concatenated_alignment.fna` refers to the concatenated BUSCO alignment, and `locus_partition.nex` denotes per-locus partitioning scheme and their best substitution models from the gene tree inference. The code to run these analyses is available at https://github.com/jeremiasivan/EucsPhylogenomics/blob/main/codes/2_run.Rmd.

### Running taxonomic consistency test

While CFs reflect the proportion of gene trees that recover a given clade, they do not reveal *which* samples from the corresponding clade move across gene trees more frequently than others (e.g., due to contamination). To address this, we developed a *taxonomic consistency test* that tracks the taxonomic placement of individual samples across gene trees. In this test, we first assigned the described taxonomy (i.e., subgenus and section) of each sample following the one-genus classification system (Nicolle 2024). For each sample in each gene tree, we identified the taxonomy of its five nearest neighbours based on pairwise phylogenetic distances - calculated using cophenetic.phylo function in the ape package (Paradis et al. 2004) - and assigned the majority group (i.e., taxonomic group represented by the majority of neighbours) as the sample’s taxon for that particular locus. Finally, we calculated each sample’s *taxonomic consistency score*, which reflects the proportion of gene trees in which the assigned taxonomy matched the sample’s described taxonomy. Here, we considered the five nearest neighbours to account for inaccurate taxonomic placements caused by mislabeled or contaminated samples (if the number of neighbours is too small), or by taxonomic groups with limited representative samples (if the number of neighbours is too large). Details of the taxonomic consistency test are provided in the Supplementary Materials, while the code required to run it is available at https://github.com/jeremiasivan/TaxonomicConsistencyTest/.

### Filtering out problematic samples and technical replicates

In this large, multi-stage study, potential errors could occur from seed collection, planting, field sampling, plate layout, DNA extraction, or barcode sequence library construction. Thus, strict quality control is required prior to the phylogenomic inference. We identified and fixed cases of mislabeling due to readily-observable plate rotations (Fig. S1; Table S1). However, 27 other samples were putatively contaminated and/or individually mislabeled, with inconsistent phylogenomic placements on the concatenated tree and/or low taxonomic consistency scores (Fig. S1). Moreover, there were four samples with invalid CCA identifiers, six described inter-sectional hybrids, and seven samples with <25% of locus recovery (Table S1). As these samples might affect gene tree inference and thus the calculations of both CFs and taxonomic consistency scores, we took a conservative approach and excluded all of them from the final tree.

Furthermore, the initial 1,024 samples included many samples that come from the same maternal family (but with different fathers) as positive controls. As expected, these siblings tended to cluster together on the concatenated tree (Fig. S1). As we were not attempting to distinguish the very low species-level concordance in this study, for each set of family-level replicates (i.e., samples with different CCA identifiers but identical voucher - those from the same maternal family), we kept one sample with the highest taxonomic consistency scores and/or least frequent recovery with distantly-related alternative groups based on the taxonomic consistency test, and excluded the rest from the final tree. This step filtered out 242 family-level replicates, as well as 37 non-CCA samples, so that our final dataset consists of 701 CCA samples.

### Inferring species tree and calculating branch concordance factors

We retrieved the BUSCO sequences of the 701 samples from the post-TreeShrink BUSCO alignments generated above and filtered out poorly-aligned sites from the subsetted alignments using ClipKIT v2.12 with default parameters (Steenwyk et al. 2020). We then inferred individual gene trees using IQ-TREE2 with the same parameters as above and re-calculated the taxonomic consistency scores across all 701 samples based on their one, three, five, and ten closest neighbours on each gene tree. Following the previous analyses, we also concatenated the 1,187 BUSCO alignments using AMAS, inferred the VeryFastTree tree, and then refined the tree using IQ-TREE2 with 1,000 UFBoot replicates (Hoang et al. 2018). We calculated qCFs of the IQ-TREE2 tree using ASTRAL-IV from ASTER (Zhang et al. 2025) with the following commands:

~~~
astral4 -u 2 -i all_busco_trees.tre -c concatenated_tree.tre
-o concatenated_tree_qCF.tre 2> astral.log
~~~

where `-u 2` reflects fully annotated branches (with per-branch local posterior probability and qCF), `all_busco_trees.tre` refers to a file containing the list of BUSCO gene trees in Newick format with one tree per line, `-c concatenated_tree.tre` fixes the topology to the IQ-TREE2 tree, `concatenated_tree_qCF.tre` refers to outputted IQ-TREE2 tree with qCF, and `astral.log` refers to outputted ASTRAL log file. Lastly, we computed the gCFs and sCFs using IQ-TREE2 (Minh et al. 2020b) with the following command:

~~~
iqtree2 -t concatenated_tree_qCF.tre --gcf all_busco_trees.tre
-p dir_busco_alignment --scf 100 --prefix iqtree_CF
~~~

where `dir_busco_alignment` refers to the directory storing individual BUSCO alignments, and `--prefix` flag defines the output filenames. The code to run these analyses is available at https://github.com/jeremiasivan/EucsPhylogenomics/blob/main/codes/2_run.Rmd.

## Results

Our final concatenated alignment comprises 701 CCA samples with a total length of 2,036,453 bp (approximately 0.4% of a eucalypt genome) and 692,711 parsimony-informative sites. This alignment includes 1,187 BUSCO loci, where each locus consists of 623 to 701 samples (average = 691 samples; Fig. S2a), and ranges in length from 218 bp to 14,130 bp (average = 1,715 bp; Fig. S2b). We then inferred a concatenated ML tree of the 701 samples based on the locus-based partition model (Fig. 3). We also calculated the gCF, sCF, and qCF of individual branches on the species tree based on the 1,187 BUSCO alignments and trees. The three measures of CFs are significantly correlated with each other (Fig. S3), as well as with branch lengths (Fig. S4).

**Figure 3.**
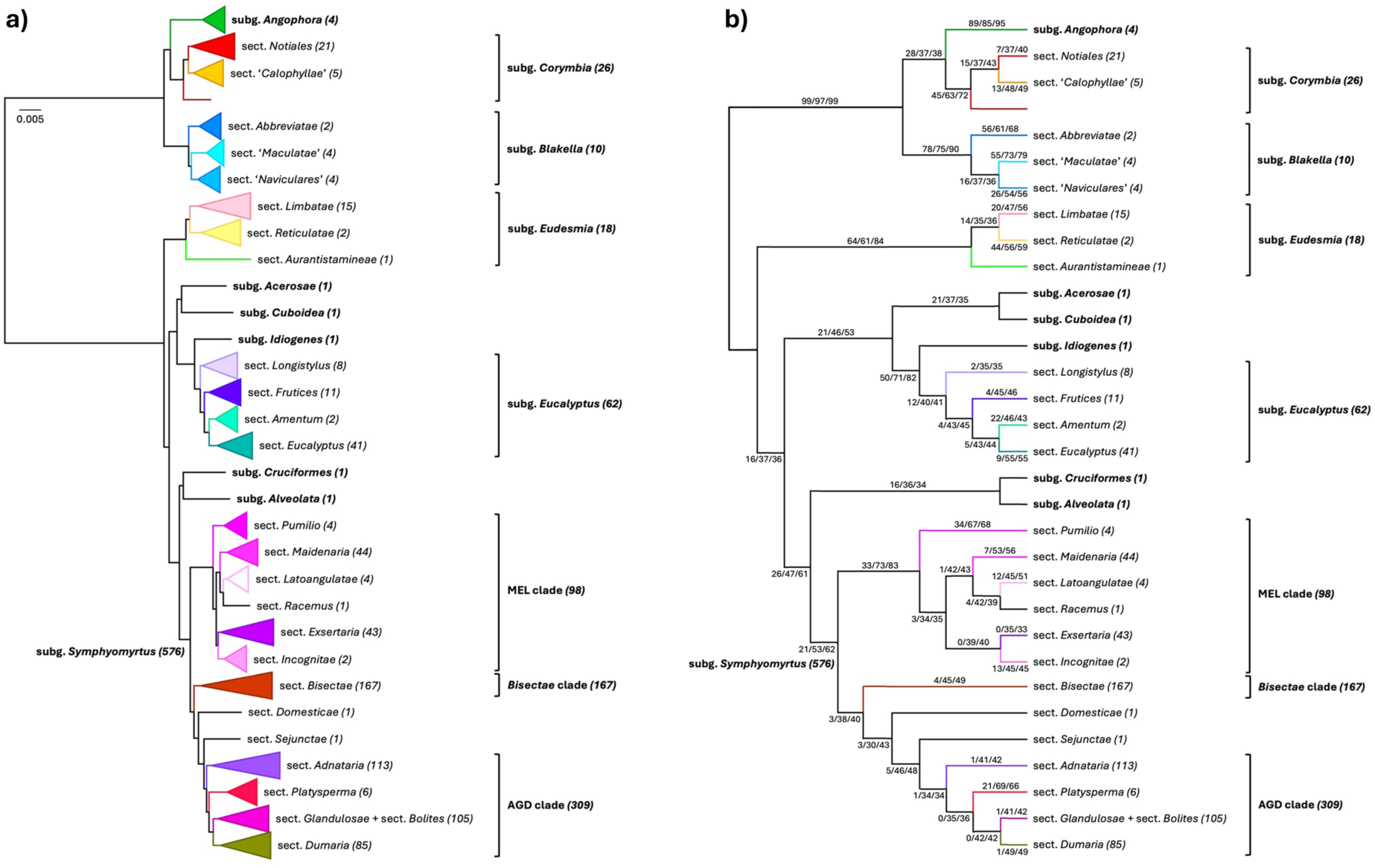
Collapsed eucalypt species tree inferred using IQ-TREE2 shown as (a) phylogenetic tree and (b) cladogram. Branch lengths of the phylogenetic tree are in the unit of substitution per site. Branch labels on the cladogram reflect gCF/sCF/qCF. Branch colours denote different sections. Numbers in parentheses reflect the number of samples in each group.

The eucalypt species tree of 701 samples shows very clear and deep separation between the CBA clade and the rest of eucalypts, with very high concordance across gene trees (99% gCF / 97% sCF / 99% qCF; Fig. 3). While most subgenera and sections are monophyletic on the species tree with 100% UFBoot support, the CFs vary substantially across groups (Fig. 3b, Tables S2-S5). The details of each taxonomic group, including its corresponding CFs and subtree topology, are discussed in the following paragraphs.

### Subgenera Angophora, Blakella, and Corymbia

For the CBA clade, we show that subg. *Corymbia* groups together with subg. *Angophora*, and both are sisters to subg. *Blakella* (Figs. 4, S5a). The branch with the highest CFs leads to subg. *Angophora* (89% gCF / 85% sCF / 95% qCF), while the second highest CFs denote subg. *Blakella* (78% gCF / 75% sCF / 90% qCF). The next two highest CFs are found on branches leading to sect. *Abbreviatae* Brooker (56% gCF; 61% sCF / 68% qCF) and sect. ‘*Maculatae*’ (Blakely) D.Nicolle (55% gCF / 73% sCF / 79% qCF), both are part of subg. *Blakella* (Fig. 4). On the other hand, subg. *Corymbia* has 45% gCF / 63% sCF / 72% qCF, while the clade that comprises *Angophora* + *Corymbia* only has 28% gCF / 37% sCF / 38% qCF, even though both have 100% UFBoot support (Table S2). If we fix *Corymbia* + *Blakella* to be monophyletic and recalculate the gCF using the same set of gene trees, the gCF of the clade goes down to 21%. If we switch the positions of the two subgenera on the tree (i.e., subg. *Blakella* is the sister to subg. *Angophora*), the *Angophora* + *Blakella* clade only has 19% gCF. This means that the arrangement of *Angophora + Corymbia* clade shown in Figure 4 is recovered by more gene trees compared to the other two alternative topologies. Interestingly, one *E. trachyphloia* F.Muell. sample (CCA1440) is placed as a sister taxon to the rest of subg. *Corymbia* and not clustered together with other samples from sect. *Notiales* – including the other *E. trachyphloia* sample (CCA5936) (Fig. 4) – similar to the tree from Crisp et al. (2024).

**Figure 4.**
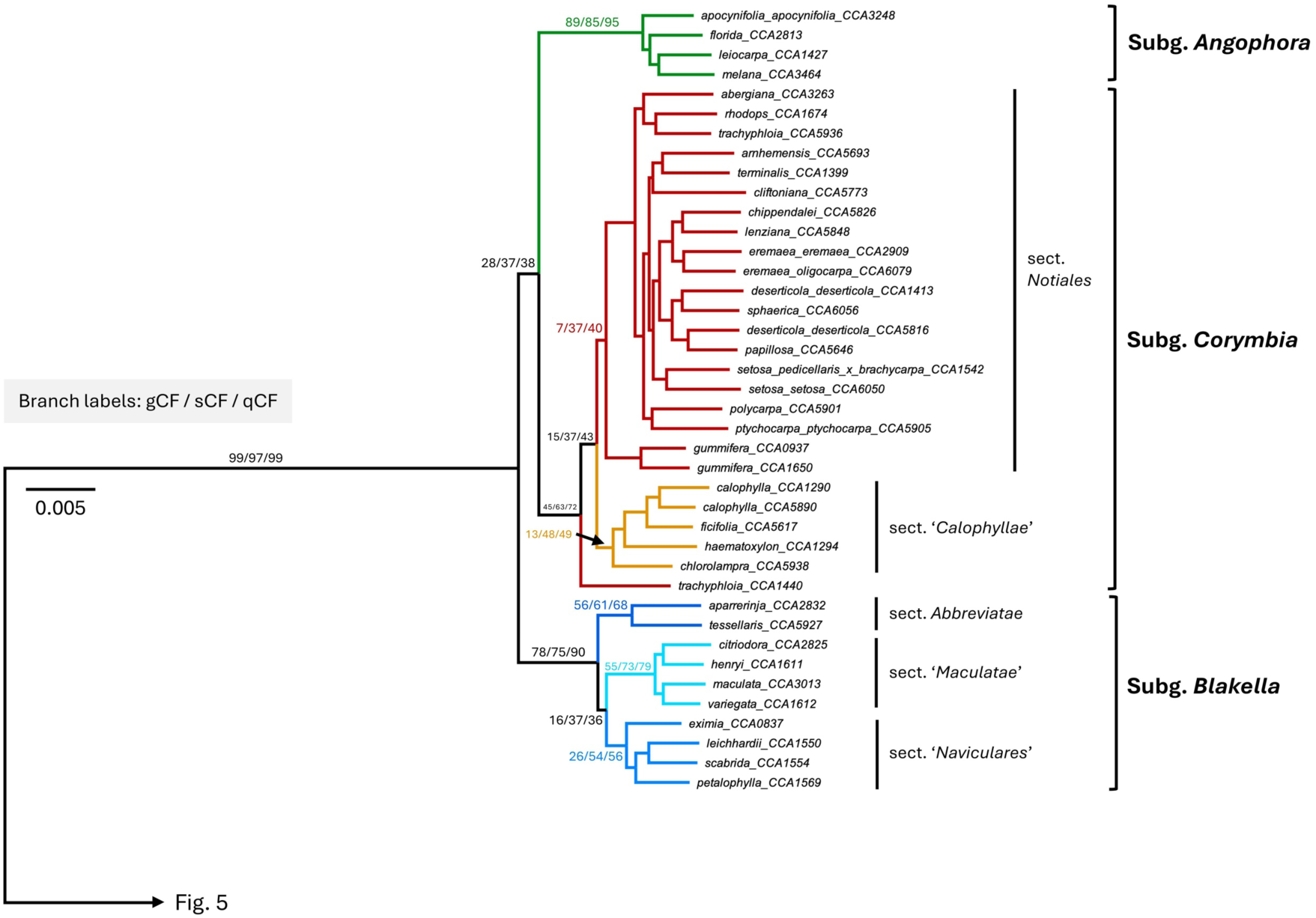
Subtree of eucalypt species tree inferred using IQ-TREE2: subgenera *Angophora, Blakella,* and *Corymbia*. Branch lengths are in the unit of substitution per site. Branch labels reflect gCF/sCF/qCF for major splits. Branch colours denote different sections. Bootstrap values of major splits are provided in Table S2.

### Subgenera Eudesmia and Eucalyptus

For the *Eucalyptus s.s.* clade, subg. *Eudesmia* (F.L.Bauer) L.A.S.Johnson & K.D.Hill diverges first (Fig. 5), with the clade having 64% gCF / 61% sCF / 84% qCF and all internal branches having >10% gCF (Table S3; Fig. S5b). The next split results in two familiar clades: one clade largely corresponding to subg. *Eucalyptus* and another to subg. *Symphyomyrtus* (Schauer) Brooker (Fig. 5). The first clade comprises three monotypic and deeply-diverged subgenera in addition to the large radiating subg. *Eucalyptus*: subg. *Acerosae* Brooker (*E. curtisii* Blakely & C.T.White), subg. *Cuboidea* Brooker (*E. tenuipes* Blakely & C.T.White), and subg. *Idiogenes* L.D. Pryor & L.A.S. Johnson ex Brooker (*E. cloeziana* F.Muell.). The branch that leads to subg. *Eucalyptus* has 12% gCF / 40% sCF / 41% qCF, with sections within this subgenus having gCF ranging from 2% (sect. *Longistylus* Brooker) to 22% (sect. *Amentum* Brooker) (Figs. 5, S5b; Table S4). However, if we consider the clade that groups subg. *Eucalyptus* and subg. *Idiogenes* together, the gCF of the corresponding branch is approximately 4.1x higher than the gCF of subg. *Eucalyptus* alone (50% gCF; Fig. 5; Table S4), suggesting that a wider circumscription of subg. *Eucalyptus* that includes *E. cloeziana* could better reflect patterns of genomic divergence among these lineages.

**Figure 5.**
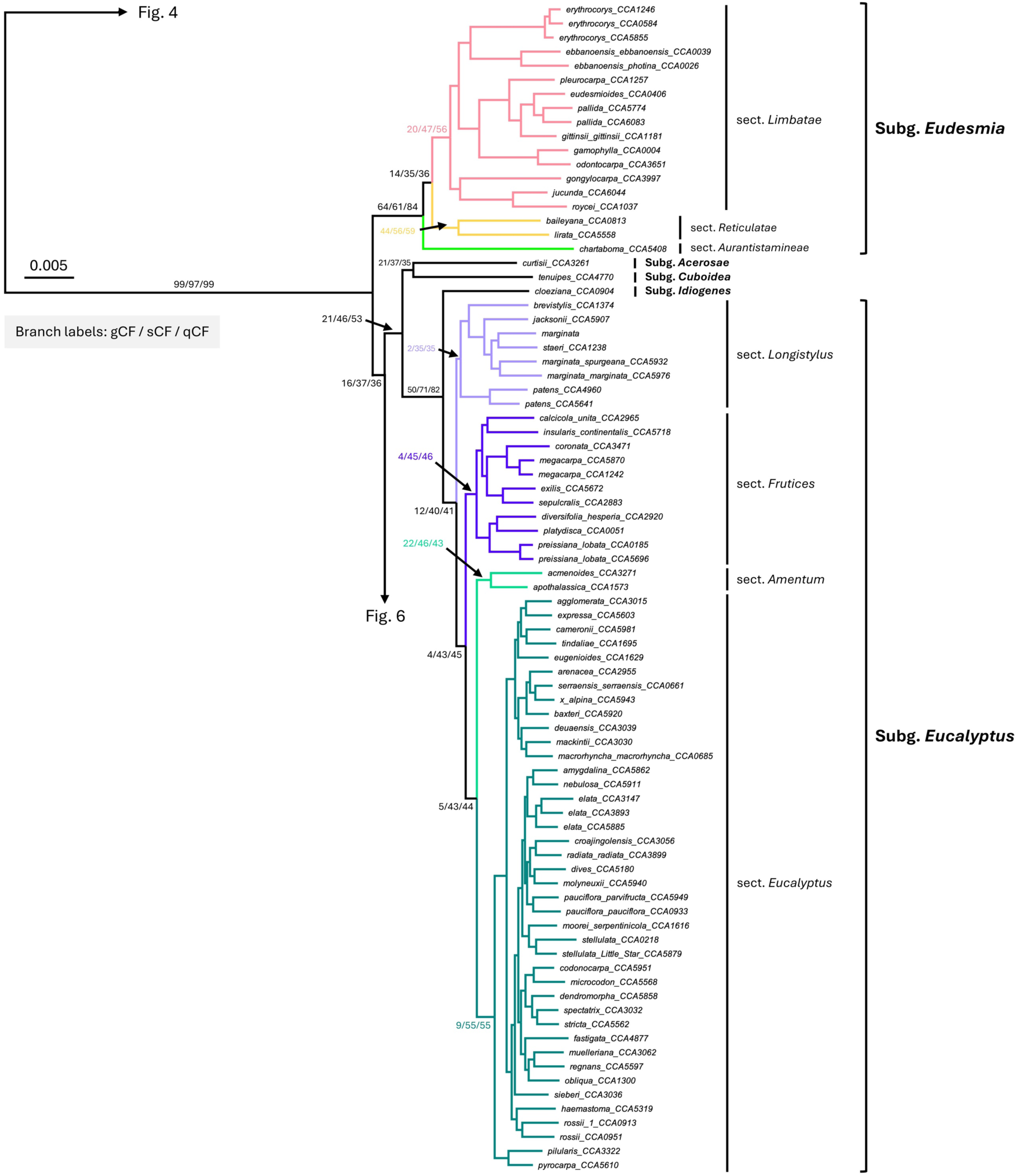
Subtree of eucalypt species tree inferred using IQ-TREE2: subgenera *Eudesmia* and *Eucalyptus*. Branch lengths are in the unit of substitution per site. Branch labels reflect gCF/sCF/qCF for major splits. Branch colours denote different sections. Bootstrap values of major splits are provided in Tables S3-S4.

### Subgenus Symphyomyrtus

The next branching event then splits the monotypic subg. *Alveolata* (Maiden) Brooker (*E. microcorys*) and subg. *Cruciformes* Brooker (*E. guilfoylei*) from the biggest subgenus in eucalypts: subg. *Symphyomyrtus* (Fig. 6). The subg. *Symphyomyrtus* has 21% gCF / 53% sCF / 62% qCF (Fig. 6; Table S5). Within this subgenus, the species tree can broadly be divided into three major clades: the “MEL” clade (consisting of sect. *Maidenaria* L.D.Pryor & L.A.S.Johnson ex Brooker, sect. *Exsertaria* L.D.Pryor & L.A.S.Johnson ex Brooker, sect. *Latoangulatae* Brooker, sect. *Pumilio* Brooker, sect. *Incognitae* (D.Nicolle) Brooker, and sect. *Racemus* Brooker; Fig. 6), the *Bisectae* clade (consisting of sect. *Bisectae* Maiden ex Brooker; Fig. 7), and the “AGD” clade (consisting of sect. *Adnataria* L.D.Pryor & L.A.S.Johnson ex Brooker, sect. *Glandulosae* (Brooker) D.Nicolle, sect. *Dumaria* L.D.Pryor & L.A.S.Johnson ex Brooker, sect. *Bolites* Brooker, and sect. *Platysperma* Brooker; Fig. 8), with a monotypic sect. *Sejunctae* Brooker (*E. cladocalyx* F.Muell.) as the sister to the AGD clade and a smaller sect. *Domesticae* Brooker (including both *E. raveretiana* F.Muell. and *E. howittiana* F.Muell. but only represented by *E. raveretiana* here) as the sister to the sect. *Sejunctae* + AGD clade (Fig. 8). Notably, subg. *Symphyomyrtus* also includes an additional small sect. *Equatoria* L.D.Pryor & L.A.S.Johnson ex Brooker consisting of *E. deglupta* Blume and *E. brachyandra* F.Muell., but we lack samples from these species here.

**Figure 6.**
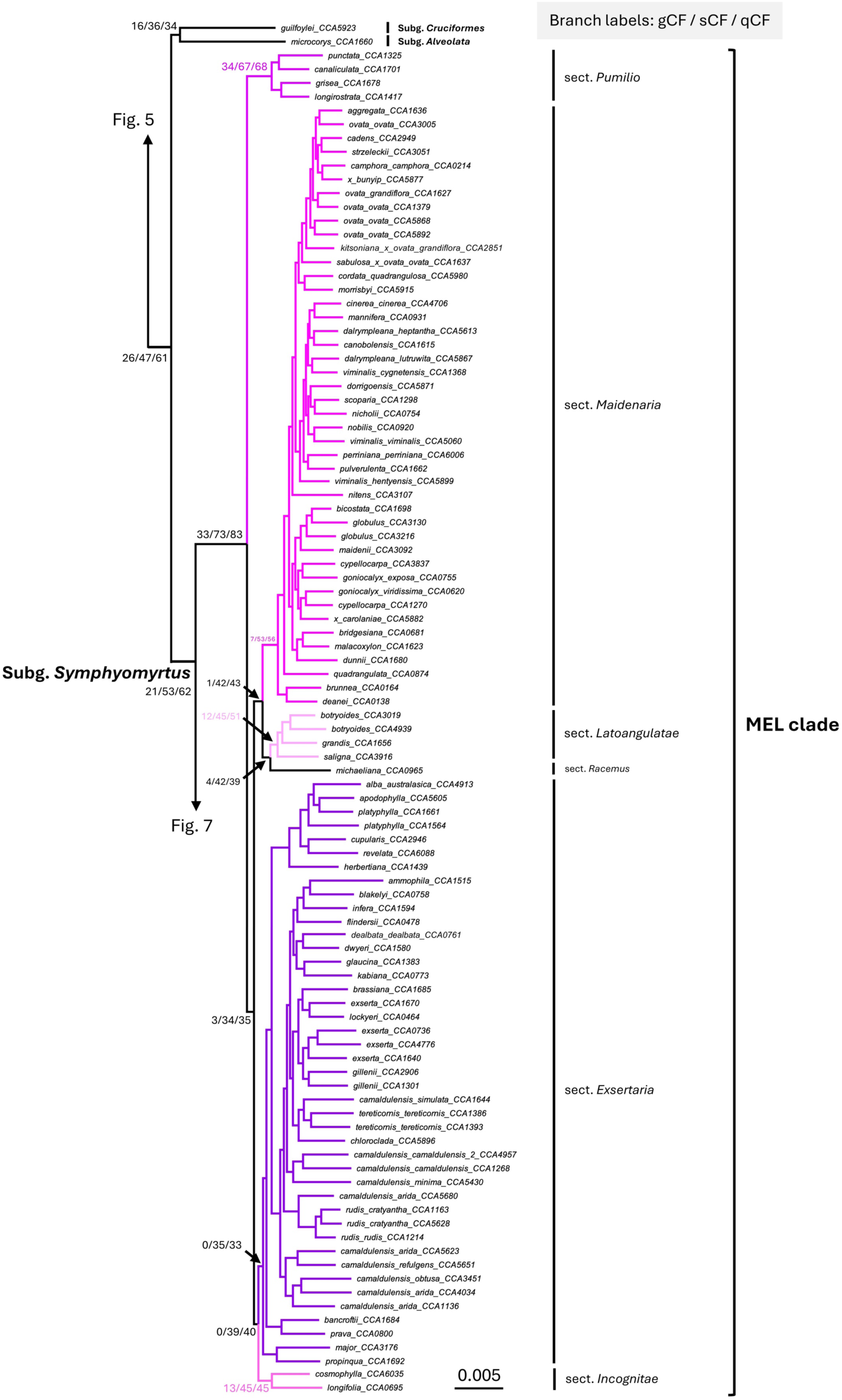
Subtree of eucalypt species tree inferred using IQ-TREE2: subgenus *Symphyomyrtus* (MEL clade). Branch lengths are in the unit of substitution per site. Branch labels reflect gCF/sCF/qCF for major splits. Branch colours denote different sections. Bootstrap values of major splits are provided in Table S5.

**Figure 7.**
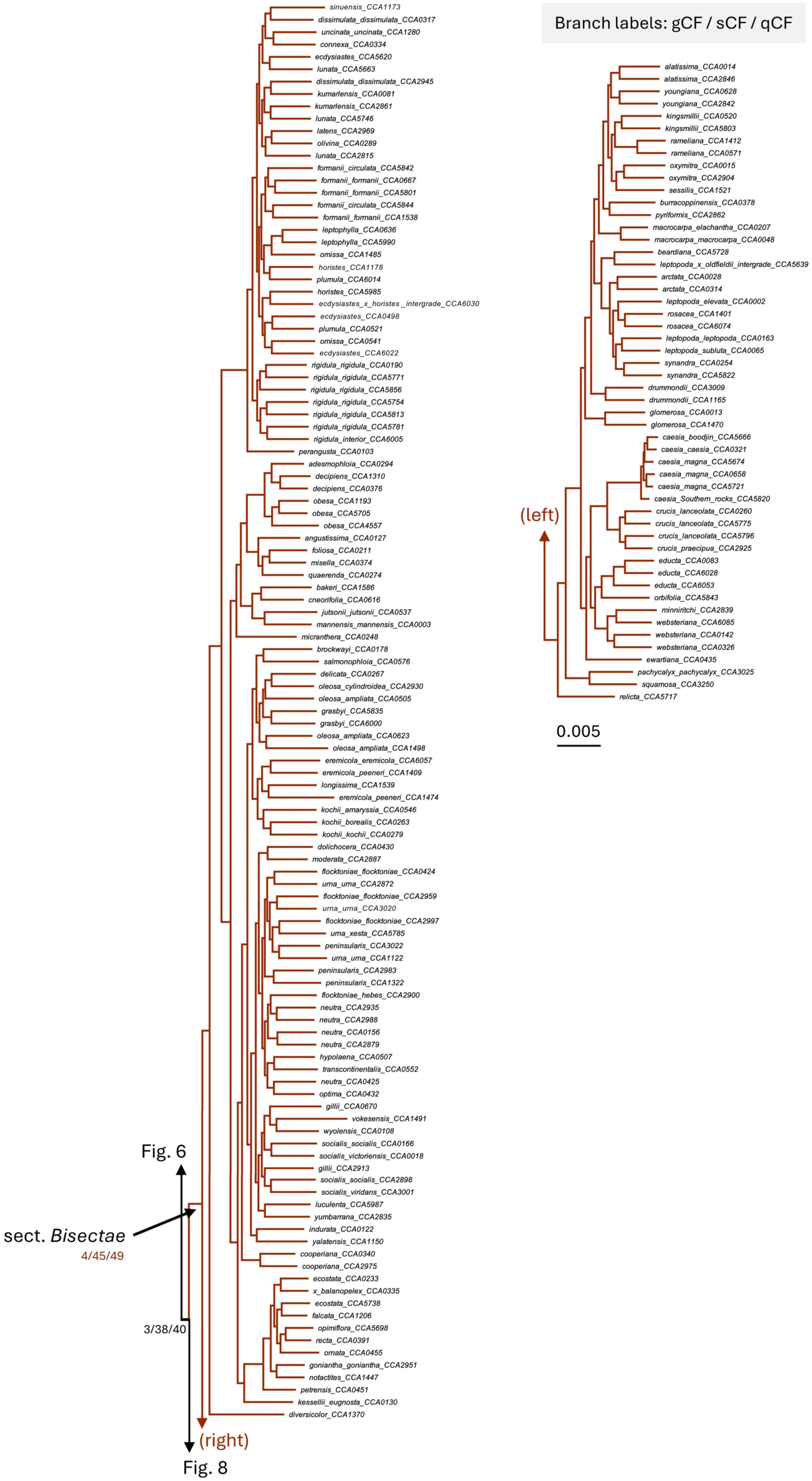
Subtree of eucalypt species tree inferred using IQ-TREE2: subgenus *Symphyomyrtus* (Bisectae clade). Branch lengths are in the unit of substitution per site. Branch labels reflect gCF/sCF/qCF for major splits. Branch colours denote different sections. Bootstrap values of major splits are provided in Table S5.

**Figure 8.**
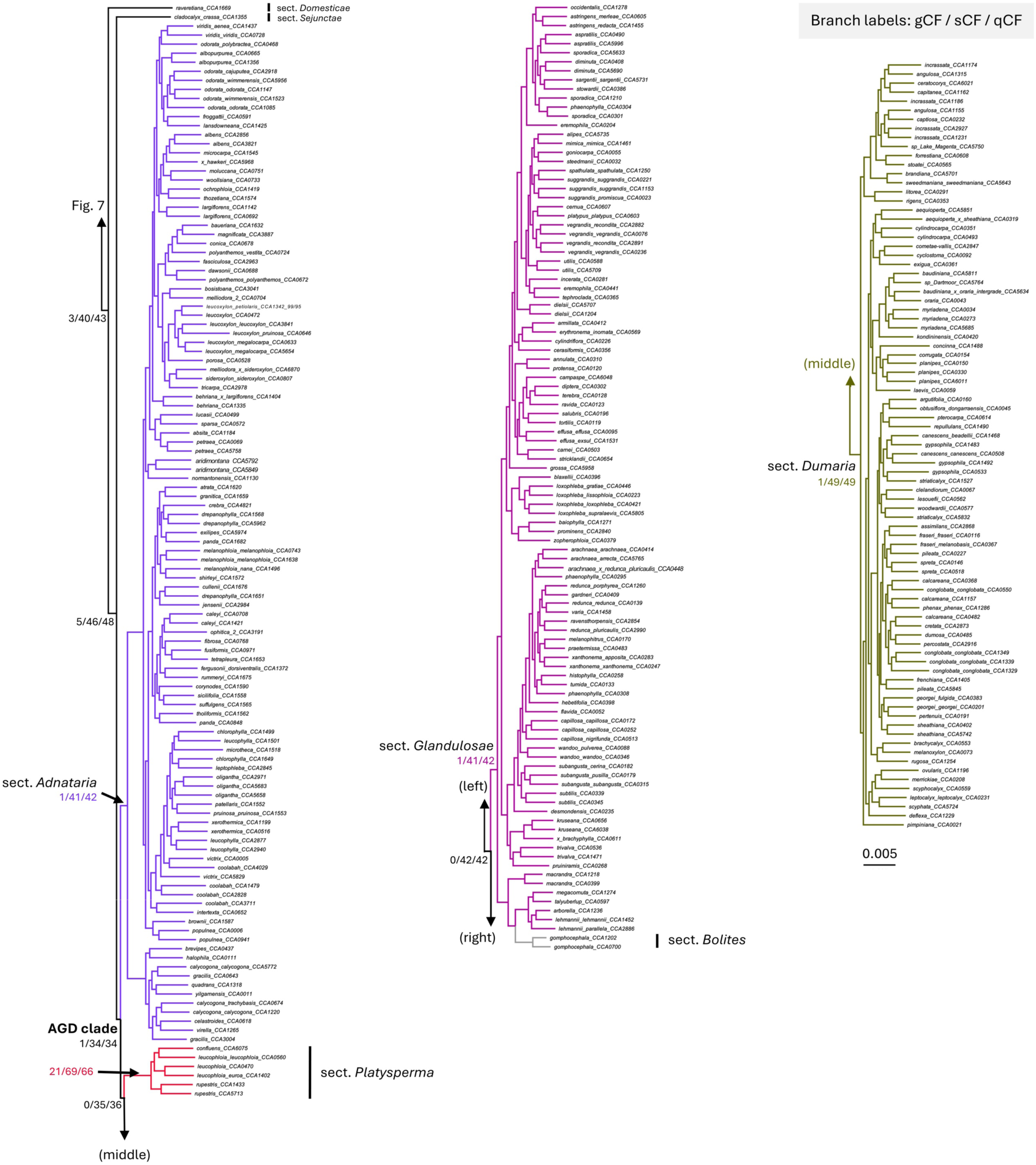
Subtree of eucalypt species tree inferred using IQ-TREE2: subgenus *Symphyomyrtus* (AGD clade). Branch lengths are in the unit of substitution per site. Branch labels reflect gCF/sCF/qCF for major splits. Branch colours denote different sections. Bootstrap values of major splits are provided in Table S5.

The MEL clade has 33% gCF / 73% sCF / 83% qCF (Fig. 6; Table S5). Within the MEL clade, sect. *Exsertaria* has practically 0% gCF with similarly low 35% sCF and 33% qCF (Fig. 6; Table S5), meaning that most gene trees do not recover the monophyly of this section on the species tree. The other separately named sections within this clade also have relatively low gCF (except for the narrow sect. *Pumilio* with 34% gCF / 67% sCF / 68% qCF; Fig. 6; Table S5). The other two clades within subg. *Symphyomyrtus*, the *Bisectae* and the AGD clades, have 4% gCF / 45% sCF / 49% qCF and 1% gCF / 34% sCF / 34% qCF, respectively (Figs. 7-8). Similar to the MEL clade, most named sections within the AGD clade have very low gCF, ranging from 0% to 1% (except for sect. *Platysperma* with 21% gCF / 69% sCF / 66% qCF; Fig. 8; Table S5). Notably, most internal branches within subg. *Symphyomyrtus* have <11% gCF, while those with relatively high gCF are found on within-species clades (Fig. S5c-e). For example, the *E. caesia* Benth. clade (Fig. S5d; right) has 66% gCF, while the two *E. gomphocephala* DC. samples from monotypic sect. *Bolites* have 51% gCF (Fig. S5e; middle). The only exception is the branch that leads to *E. brandiana* Hopper & McQuoid (CCA5701) and *E. sweedmaniana* Hopper & McQuoid (CCA5643), which has 42% gCF (Fig. S5e; right).

### Taxonomic consistency scores

For each of the 701 samples, we calculated their taxonomic consistency scores based on their one, three, five, and ten nearest neighbours across 1,187 BUSCO trees. Compared to CFs that describe dataset-wide concordance for each branch in the tree, taxonomic consistency scores describe the taxonomic placement of each sample across gene trees. Figures S6-S8 show that the taxonomic consistency test produces similar results across different numbers of neighbours for well-represented taxonomic groups (in this case, roughly 20 samples per group). Following the initial analyses (Fig. S1), we discuss the results that consider the five nearest neighbours on each gene tree in the following paragraphs.

As shown in Figure S6, the consistency scores across subgenera are generally high, meaning that taxonomic placement of a sample is highly consistent across loci at the level of the subgenus. For subg. *Angophora* samples, the average taxonomic consistency score is 84% (Fig. S6; top-left). However, note that there are only four subg. *Angophora* samples, which might have prevented it from being selected as the major taxonomic group in some gene trees. On the other hand, subg. *Blakella* has a 95% average consistency score (Fig. S6; top-middle), while subg. *Corymbia* has a 98% average consistency score (Fig. S6; top-right). This is consistent with limited gene flow between present-day samples of these subgenera, which supports the relatively high gCF values of the corresponding nodes on the species tree (Fig. 3). Similar to the CBA clade, the average consistency scores across the three major subgenera in *Eucalyptus s.s.* are high. Specifically, subg. *Eudesmia* has a 91% average consistency score (which is affected by the three samples from undersampled sect. *Reticulatae* Brooker and sect. *Aurantistamineae* Bayly & R.Fowler with <82% consistency scores), while subg. *Eucalyptus* and subg. *Symphyomyrtus* have 98% and 99% average scores, respectively (Fig. S6; bottom).

At section level, sections under subg. *Blakella* and subg. *Corymbia* with ≤5 samples (i.e., sect. ‘*Maculatae*’, sect. ‘*Naviculares*’ (Maiden) D.Nicolle, sect. ‘*Calophyllae*’ (Parra-Os. & Ladiges) D.Nicolle) have consistency scores ranging from 50% to 71% (Fig. S7). Similar to subg. *Angophora*, these scores might represent a lower bound due to the small number of samples per group. For a well-sampled sect. *Notiales*, the average consistency score is 91% across 21 samples, with 18 samples having ≥92% scores and the other three having much lower scores: one from *E. trachyphloia* sample (CCA1440; 64%) and two from *E. gummifera* (Sol. Ex Gaertn.) Hochr. (CCA0937 and CCA1650; both 67%) (Fig. S7; middle-left). However, consistent with the subgenus consistency scores (Fig. S6), the alternative group for each section (i.e., taxonomic group with the second-highest count across samples) always comes from the same subgenus as the expected section (Fig. S7). For well-sampled sections (i.e., sections with ≥20 samples) in the other three major subgenera, there are four sections with <90 average consistency scores: sect. *Exsertaria* (82%), sect. *Adnataria* (88%), sect. *Glandulosae* (85%), and sect. *Dumaria* (83%) (Fig. S8). For sect. *Exsertaria*, the alternative group is sect. *Maidenaria* with 10% average scores across samples (Fig. S8; top-middle). For sect. *Adnataria*, consistency scores are bimodally distributed across samples, with all eleven samples from ser. *Heterostemones* Benth. forming a small clade sister to the rest of sect. *Adnataria* (Fig. 8) with <77% consistency scores (while the rest have >83%; Fig. S8; middle-right). For the other two AGD sections, roughly 8% of gene trees recover sect. *Glandulosae* samples within the sect. *Dumaria* clade (Fig. S8; bottom-left), while approximately 10% gene trees place sect. *Dumaria* samples together within the sect. *Glandulosae* clade (Fig. S8; bottom-middle).

## Discussion

Prior studies investigating the molecular basis for eucalypt taxonomy primarily focused on a single bifurcating species tree and its statistical support, which has been shown to be inadequate in explaining the high gene tree discordance found in this group due to ILS and introgression (McLay et al. 2023; Orel et al. 2026). Moreover, eucalypt phylogenetic studies often focus on resolving higher taxonomic ranks (e.g., Thornhill et al. 2019; Crisp et al. 2024), with less attention to intermediate ranks, such as subgenus and section. Here, we present a phylogenomic perspective on current eucalypt taxonomy by calculating the concordance factors (CFs) of each subgenus and section across 1,187 BUSCO loci comprising approximately 500 described species (Nicolle 2024). While most taxonomic groups are monophyletic on the species tree with 100% bootstrap statistical support given the huge data set (Fig. 3; Tables S2-S5), CFs vary substantially across groups. We discuss major findings of our analyses in the following paragraphs.

### Subgenera *Angophora*, *Corymbia*, and *Blakella* have moderate-to-high concordance

In the CBA clade, we show that subg. *Blakella* has relatively high CFs of 78% gCF / 75% sCF / 90% qCF, compared to subg. *Angophora* (89% gCF / 85% sCF / 95% qCF) and subg. *Corymbia* (45% gCF / 63% sCF / 72% qCF) (Fig. 4; Table S2). Crisp et al. (2024) reported lower concordance for these groups (but with similar patterns of subg. *Angophora* having the highest gCF and sCF, followed by the two other subgenera). The lower gCF of subg. *Corymbia* (compared to the other two subgenera) might reflect one or more samples from this subgenus often clustering together with other subgenera across gene trees, which is also supported by the taxonomic consistency test. For example, there are three samples from subg. *Corymbia* that have relatively low consistency scores at the level of section: one *E. trachyphloia* sample (CCA1440) and two *E. gummifera* samples (CCA0937, CCA1650) (Fig. S7). Similar to the tree from Crisp et al. (2024) and Orel et al. (2026), *E. trachyphloia* is the sister to the rest of subg. *Corymbia*, while *E. gummifera* is the sister to the rest of sect. *Notiales* (Fig. 4). Orel et al. (2026) demonstrated deep divergence between the plastid genomes of their two *E. trachyphloia* samples despite them forming a well-supported monophyletic clade based on the nuclear data. This means that the variation in phylogenetic placements among the different *E. trachyphloia* samples here may reflect introgression, even though the evidence of successful hybridisation between *E. trachyphloia* and other species is still lacking (Dickinson et al. 2012). Interestingly, species of sect. ‘*Maculatae*’ have been shown to hybridise with several other species from subg. *Corymbia*: for example, *E. × nowraensis* Maiden refers to a naturally occurring hybrid between the aforementioned E. *gummifera* and E. *maculata* Hook. (Hill and Johnson 1995). In addition, Dickinson et al. (2012) successfully hybridised *E. torelliana* with the subg. *Corymbia* species (*E. clarksoniana* D.J.Carr & S.G.M.Carr and *E. erythrophloia* Blakely) in controlled settings. These observations appear to support a more recent divergence of subg. *Corymbia* and subg. *Blakella* compared to subg. *Angophora*.

However, while we did find that the *Angophora* + *Corymbia* clade has a relatively low gCF of 28% (with similarly low sCF and qCF of 37% and 38%, respectively; Fig. 4; Table S2), the same set of gene trees also show that the alternative groupings have lower gCFs of 21% (*Corymbia* + *Blakella*) and 19% (*Angophora* + *Blakella*). These results are consistent with a recent study by Orel et al. (2026). Using a larger set of 168 samples from 97 CBA species, the authors showed that the *Angophora* + *Corymbia* clade is recovered by 38% of gene trees, while the alternative groupings are supported by 36% (*Corymbia* + *Blakella*) and 26% gene trees (*Angophora* + *Blakella*), respectively. This might reflect a high level of ILS during divergence of the group approximately 40 Ma (Thornhill et al. 2019; Orel et al. 2026), followed by independent accumulation of genetic divergence between the two groups.

### Subgenus *Eucalyptus* has higher concordance when grouped with subgenus *Idiogenes*

We found that subg. *Eucalyptus* only has 12% gCF / 40% sCF / 41% qCF, which is generally lower than other subgenera (Fig. 5; Tables S2-S5). However, if we included the *E. cloeziana* sample (from monotypic subg. *Idiogenes*), the gCF of the branch goes up by 4.1x, while the sCF and qCF increase by roughly 2-fold (Fig. 5; Table S4). In the one-genus classification system, Brooker (2000) placed *E. cloeziana* in a monotypic subgenus due to its distinct morphological features with respect to subg. *Eucalyptus*, such as its flaky rough bark, strongly discolourous adult leaves, compound inflorescences, and four-rowed ovules. However, molecular studies have shown that *E. cloeziana* is genetically similar to species from subg. *Eucalyptus*, sometimes placing it within the subgenus rather than as a sister lineage (Bayly et al. 2013; Schuster et al. 2018). Moreover, Stokoe et al. (2001) reported natural hybridisation between *E. cloeziana* and *E. acmenoides* Schauer of subg. *Eucalyptus*, supported by both morphology and molecular markers. These findings suggest that the view of *E. cloeziana* as a distinct, isolated subgenus should be reassessed with respect to subg. *Eucalyptus*. Our analyses show that this single change would increase the extent to which the subg. *Eucalyptus* is predictive of gene tree topologies by more than four-fold, from 12% to 50% (Table S4).

### Subgenus *Symphyomyrtus* exhibits very high discordance

Even though subg. *Symphyomyrtus* has 21% gCF, most sections within this subgenus have very low gCF (Figs. 6-8, Table S4), reflecting pervasive ILS due to the rapid radiation of these sections. Therefore, instead of seeing each section as a distinct, evolutionary-diverged lineage, we discuss these sections as three different clades, which we hereby refer to as *supersections*: the MEL clade, the *Bisectae* clade, and the AGD clade. The term “MEL” was derived from Nicolle and Jones (2019), who used the “MEL+5” term in referring to the three biggest sections (i.e., sections with the most number of species) of this clade (sect. *Maidenaria*, sect. *Exsertaria*, and sect. *Latoangulatae*) along with five smaller “sections” (sect. *Racemus*, sect. *Incognitae*, and sect. *Pumilio,* as well as ser. *Liberivalvae* (Brooker) D.Nicolle & R.C.Jones and ser. *Similares* (Brooker) D.Nicolle & R.C.Jones that were previously considered sections but are now treated as series within sect. *Exsertaria* and sect. *Incognitae*, respectively). On our tree, the MEL clade has 33% gCF and relatively high 73% sCF / 83% qCF (Table S5), with clear separation from the other two clades (Figs. 3, 6-8). Inter-sectional hybrids have been well documented within the MEL clade (Griffin et al. 1988; Larcombe et al. 2015), but it has almost zero crossability with the *Bisectae* and AGD clades (Larcombe et al. 2015). Possible exceptions are a rare *E. drummondii* × *E. rudis* hybrid (Rossetto et al. 1997) and a putative *E. camaldulensis* subsp. *camaldulensis* × *E. largiflorens* hybrid (also known as E. × *oxypoma* Blakely; Nicolle 2024). Thus, the MEL clade can be seen as a largely distinct lineage from the other two.

On the other hand, the *Bisectae* and AGD clades have 4% gCF and 1% gCF, respectively (Figs. 7-8, Table S5). Here, we coined the term “AGD” in referring to the three biggest sections in this clade (sect. *Adnataria*, sect. *Glandulosae*, and sect. *Dumaria*) along with two smaller related sections (sect. *Platysperma* and sect. *Bolites*). Notably, we confirm previous findings from Crisp et al. (2024) that the monotypic sect. *Bolites*, consisting of *E. gomphocephala*, is extremely closely related to sect. *Glandulosae*. In fact, the two *E. gomphocephala* samples in our dataset appear to be nested within sect. *Glandulosae* on the species tree (Fig. 8), particularly ser. *Lehmannianae* D.J.Carr and S.G.M.Carr. As such, *E. gomphocephala* should probably be considered as a member of sect. *Glandulosae* and perhaps even ser. *Lehmannianae*, given its current phylogenetic position.

Sect. *Glandulosae* was previously considered as a series of sect. *Bisectae* based on their morphology (Brooker 2000), but they were shown to be paraphyletic based on later phylogenetic analyses (Steane et al. 2002). To maintain the monophyly of the section, *Glandulosae* was then elevated to a section level (Nicolle 2022). As members of these clades are very young and still radiating (Thornhill et al. 2019; Orel et al. 2026), the concordance of the internal branches on the species tree is generally very low (i.e., <11% gCF; Figs. S5d-e). Moreover, a number of hybrids have been documented between the *Bisectae* and AGD clades (Griffin et al. 1988; Delaporte et al. 2001; Nicolle 2022), which suggests that intrinsic reproductive barriers between the two clades may be incomplete. Despite the incomplete reproductive barriers between the two clades and the very low concordance of each group, we discuss the two clades separately here because: (i) members of each clade consistently group together with other members of their expected clade across gene trees based on the taxonomic consistency test (Fig. S8); and (ii) combining the two clades together may not be taxonomically informative as it would represent approximately two-thirds of named species in subg. *Symphyomyrtus* (Nicolle 2024).

### Future work and concluding remarks

Overall, this study shows the limitations of morphology-based taxonomy - which is represented as a single concatenated species tree (with its statistical support) - in describing the topological variation of gene trees across the genome. While a species tree might represent our “best” point estimate of evolutionary relationships between individual subgenera and sections, the low concordance found in several groups reflect much more complex relationships in this group that cannot be captured by a single bifurcating species tree nor a hierarchical taxonomy. Thus, we propose several taxonomic recommendations as discussed above and present a phylogenomic inference pipeline that emphasises the variability of evolutionary history along the genome by calculating concordance factors (CFs) and taxonomic consistency scores across thousands of loci. We believe that our phylogenomic inference pipeline will be useful for other complex groups of taxa with multiple radiations. It is also adaptable for future studies, as it allows users to add their own samples with common short-read genome data onto the previously-published species tree (such as the eucalypt tree shown in Figure 3) following the same steps conducted in this study: locus extraction from short-read data using CAPTUS (Ortiz et al. 2026), species and gene trees inference using IQ-TREE2 (Minh et al. 2020b), calculation of CFs using ASTRAL-IV (Zhang et al. 2025) and IQ-TREE2 (Minh et al. 2020a), and the taxonomic consistency test developed in this study. Please refer to https://github.com/jeremiasivan/EucsPhylogenomics for details on how to run the pipeline using your own datasets.

While we put extra attention to fix putative mislabeling and/or contamination, we acknowledge that we might have not addressed all errors present in the dataset, and that any dataset will have unique samples that do not represent a larger group. Beyond sampling, the inferred locus sequences might still be affected by reference bias during mapping, despite the attempts to perform *de novo* assembly before mapping and use target genes from closely-related taxa (Ivan and Lanfear 2026a). This might result in an increased estimate of genetic divergence among samples from the CBA and *Eucalyptus s.s.* clades (Fig. 3). In addition, we note that our locus sequences are unphased due to difficulties in phasing short sequencing reads, relying on several factors such as read depth and reference selection (e.g., Nauheimer et al. 2021). As the maternal and paternal haplotype representation is randomly resolved as a single locus sequence, a chimeric sequence from a wide hybrid might put the sample deeper on the tree. With the development of long-read sequencing technologies and the generation of high-quality eucalypt reference genomes (Ferguson et al. 2024b, 2024a; Lötter et al. 2025; Zhuang et al. 2026), future phylogenomic studies should include haplotype-resolved reference genomes to better understand finer-scale taxonomic relationships in eucalypts. Lastly, eucalypts have been shown to exhibit high recombination rate and low linkage disequilibrium (Gion et al. 2016; Murray et al. 2019), which means that a single BUSCO locus used in this study might span multiple recombination breakpoints and thus contain multiple distinct evolutionary histories. As gene tree inference assumes a single genealogy per locus, the inferred gene tree topology might be inaccurate due to information loss from concatenation (Kubatko and Degnan 2007; Ivan et al. 2025; Ivan and Lanfear 2026b), thus potentially affecting the gCF calculation. Gene tree estimation error should be less pronounced for sCF (which is estimated directly from sequence alignment) and qCF (which is estimated from quartets). However, both gCF and qCF might be inflated when discordance is high, because there are only three possible resolutions for any quartet, and each quartet must be assigned to one of them regardless whether the data have sufficient signal to do so (Lanfear and Hahn 2024). Therefore, it is important to calculate and compare the three measures of concordance factors for a robust interpretation of branch concordance factors. Despite these limitations, we believe that our study provides the most comprehensive phylogenomic perspective on eucalypt taxonomy to date, which is not only useful for understanding current and future eucalypt evolutionary histories, but also is broadly applicable for other groups of taxa.

## Data Availability

The raw short-read data generated in this study are available on ENA PRJEB108804 and JGI Genome Portal (proposal ID: 508017). The phylogenomic inference pipeline (DOI: <u>10.5281/zenodo.22875230</u>) and taxonomic consistency test (DOI: <u>10.5281/zenodo.22875205</u>) are available on Zenodo, while individual locus alignments and trees generated in this study are available on FigShare (DOI: <u>10.6084/m9.figshare.33919381</u>).

## Conflict of Interest

D.N. is the main author of Nicolle et al. (2025) that proposed the one-genus classification system for eucalypts, while R.A. is the contributing author of Cook et al. (2025) that supported the four-genus classification system.

## Acknowledgements

We thank Chloe Tan, Dave Stanley, Juan Garces, Grey Monroe, Momena Khandaker, Helen Bothwell, James Kondilios, Scott Ferguson, Ming-Dao Chia, Jasmine Janes, and Andrew Almonte for help with collecting and sample processing. Claude Sonnet 4.6 (Anthropic 2026) was used to help generate the `config.yaml` and `run_pipeline.R` scripts in both the phylogenomic inference pipeline and taxonomic consistency test. All AI-generated code has been reviewed and validated by the authors before use.

## Funding

This work was supported by Australian Research Council (grant numbers CE140100008, DP150103591, LP240200679); and the Joint Genome Institute Community Sequencing Program Project ID: 508017.

## Notes

### Summary of Updates

- Updated authors' ORCiD ID - Updated DN's correspondence - Updated informal sect. 'Maculatae', sect. 'Naviculares', sect. 'Calophyllae' - Updated E. gummifera's authorship

