## Supplementary Materials for "Comprehensive Phylogenomic Inference of Eucalypts Reveals Taxonomic Limits through Gene Tree Concordance"

### SUPPLEMENTARY TEXT

#### Extracting Reference BUSCO Loci

We downloaded the genome assemblies of 36 Eucalypts from (Ferguson et al. 2024); BioProject PRJ509734) using NCBI Datasets (https://www.ncbi.nlm.nih.gov/datasets/) based on their accession numbers. Then, we ran BUSCO v5.8.0 (Simão et al. 2015; Manni et al. 2021) on each genome assembly using `eudicots_odb10` because it is the most specific taxonomic group that encompasses Eucalypts, with MetaEuk (Levy Karin et al. 2020) as the gene prediction tool as it offers a balance between speed and accuracy (i.e., it is faster than Augustus (Stanke and Waack 2003) and less sensitive to highly-divergent assemblies compared to miniprot (Li 2023)). For each genome assembly, we retrieved the nucleotide sequences of all BUSCO loci that were identified as single-copy and complete – excluding duplicated and fragmented BUSCO to ensure that the subsequent BUSCO extractions using CAPTUS (Ortiz et al. 2026) were not affected by paralogs – totaling in 2,310 BUSCO loci.

For each of the 2,310 BUSCO loci, we removed sequences that were 5% shorter or longer than the median length, because manual inspection of preliminary alignments suggested that long insertions and/or deletions tended to be associated with the BUSCO pipeline recovering misidentified orthologues. Then, we aligned the sequences of each BUSCO using MAFFT v7.520 (Katoh et al. 2002; Katoh and Standley 2013) with default settings (FFT-NS-2). For each BUSCO alignment, we ran TAPER v1.0.2 (Zhang et al. 2021) to mask putative alignment errors, and removed sequences and sites with ≥50% gaps to filter out badly-aligned samples and/or regions using Biostrings package (Pagès et al. 2024) in R (R Core Team 2023).

Next, we checked each BUSCO alignment by eye using Geneious Prime v2025.0.1 (https://www.geneious.com), manually removing individual taxa from each alignment if they contained one or more highly-divergent regions that might reflect putative paralogy. From this cleaned set of BUSCO alignments, we excluded those with zero species from the *Angophora, Blakella*, or *Corymbia* subgenera ("CBA" clade), and/or <17 species from the other *Eucalyptus* subgenera (i.e., half of the initial number of genome assemblies from Ferguson et al. (2024)). This resulted in a final set of 1,187 reference BUSCO alignments, where each alignment contained at least one representative from CBA clade and at least 17 representatives from the rest of eucalypts. We then unaligned all BUSCO alignments by removing all gaps from each sequence. Finally, we created two target FASTA files of sequences for each BUSCO: one consisting of only species from the CBA clade, and one of only species from the other *Eucalyptus* subgenera. These target files were used to extract BUSCO sequences from short-read datasets using CAPTUS.

**Taxonomic Consistency Test**

We developed a taxonomic consistency test that tracks the taxonomic placement of each sample on every gene tree. The test begins by assigning a taxonomy group to every tip on the trees, including our samples of interest. For each sample, we then identify its *n* closest neighbours on each gene tree (i.e., tips with the smallest pairwise phylogenetic distances to the sample) and retrieve the most common taxonomic group among them, which becomes the sample's assigned taxon for the corresponding gene tree. In case of ties, all groups are retained and assigned equal weight (i.e., each tied group has a weight of 1/*k*, where *k* represents the total number of tied groups). We repeat this process across individual gene trees, and count the proportion of gene trees that support each unique taxonomic group. We then calculate a *taxonomic consistency score* for each sample, which reflects the weighted proportion of gene trees that recover their expected (i.e., initially assigned) taxonomy. For instance, a score of 90% means that the corresponding sample groups with its expected taxon in 90% of gene trees, and with other taxa in the remaining 10%. Using the taxonomic consistency scores, we will be able to not only measure the proportion of gene trees that support the current taxonomy of a sample, but also identify which alternative groupings that are recovered by the remainder of the trees. Note that if the sample's taxonomy is unknown, we instead assign the taxonomic group recovered by the majority of gene trees as the 'expected' taxon of the sample. The code to run these analyses is available at https://github.com/jeremiasivan/TaxonomicConsistencyTest, and also has been incorporated in the phylogenomic inference pipeline proposed in this study.

Even though CFs and taxonomic consistency scores both quantify topological variation across gene trees, they capture different underlying phenomena. Gene tree discordance driven by ILS and hybridisation is expected to occur at a roughly constant rate over time, meaning that the resulting topological variation should scale primarily with branch lengths - a proxy of genetic divergence - rather than with absolute time. As divergence increases, ILS (and eventually hybridisation) should decline, resulting in less frequent discordant topologies. In other words, two highly-diverged, present-day taxa are much more likely to be separated than grouped together on individual gene trees. This should directly affect taxonomic consistency scores, because they are computed based on current placements of the sample. CFs, on the other hand, should be less sensitive to this effect, because they capture cumulative signals of historical events rather than current evolutionary relationships. For example, if clades X and Y diverged via rapid radiation, the branch separating them on the species tree is likely to have low CFs, since ILS was pervasive at the moment of that split. However, if the two clades then evolved independently over a long period of time, species from clade X will have accumulated substitutions that are specific to that lineage (and likewise for clade Y species). As a result, present-day samples from clade X are very likely to cluster together with other clade X species on individual gene trees, resulting in high taxonomic consistency scores. Thus, taxonomic consistency scores could inform present-day gene flow barriers between taxa (which reflect current taxonomic boundaries), while CFs reveal deeper evolutionary history.

### SUPPLEMENTARY TABLES

**Table S1.** Metadata of 1,024 eucalypt samples used in this study (excluding three CCA samples with zero locus recovered). Sample: tip label on the tree; voucher: tree voucher; subgenus: described subgenus from Nicolle (2024); section: described section from Nicolle (2024); n_locus: number of locus after the filtering steps; notes: additional notes of individual samples, including label fix and/or exclusion criterion.

| sample | voucher | subgenus | section | n_locus | notes |
|---|---|---|---|---:|---|
| abergiana_CCA3263 | DN 1303 | *Corymbia* | *Notiales* | 1144 | |
| absita_CCA1184 | DN 1154 | *Symphyomyrtus* | *Adnataria* | 1187 | label fix (previous: CCA1278) |
| acies_CCA6062 | DN 225 | *Eucalyptus* | *Frutices* | 1172 | putative contamination / mislabelling |
| acmenoides_CCA3271 | DN 1299 | *Eucalyptus* | *Amentum* | 1135 | |
| adesmophloia_CCA0294 | DN 216 | *Symphyomyrtus* | *Bisectae* | 1187 | |
| aequioperta_CCA5851 | DN 5458 | *Symphyomyrtus* | *Dumaria* | 1187 | |
| aequioperta_x_sheathiana_CCA0319 | DN 307 | *Symphyomyrtus* | *Dumaria* | 1178 | |
| agglomerata_CCA3015 | DN 1754 | *Eucalyptus* | *Eucalyptus* | 1187 | |
| aggregata_CCA1636 | DN 1224 | *Symphyomyrtus* | *Maidenaria* | 1185 | |
| alatissima_CCA0014 | DN 7 | *Symphyomyrtus* | *Bisectae* | 1187 | |
| alatissima_CCA2846 | DN 1530 | *Symphyomyrtus* | *Bisectae* | 1185 | |
| alba_australasica_CCA4913 | DN 4211 | *Symphyomyrtus* | *Exsertaria* | 1166 | |
| albens | | *Symphyomyrtus* | *Adnataria* | 1187 | invalid CCA identifier |
| albens_CCA2856 | DN 1676 | *Symphyomyrtus* | *Adnataria* | 1186 | |
| albens_CCA2857 | DN 1676 | *Symphyomyrtus* | *Adnataria* | 1159 | technical replicates |
| albens_CCA3821 | DN 2898 | *Symphyomyrtus* | *Adnataria* | 1154 | |

| | | | | | |
|---|---|---|---|---|---|
| albens_Ferguson2024 | | *Symphyomyrtus* | *Adnataria* | 1186 | not CCA |
| albopurpurea_CCA0662 | | *Symphyomyrtus* | *Adnataria* | 996 | technical replicates |
| albopurpurea_CCA0665 | | *Symphyomyrtus* | *Adnataria* | 1185 | |
| albopurpurea_CCA1345 | DN 825 | *Symphyomyrtus* | *Adnataria* | 1187 | putative contamination / mislabelling |
| albopurpurea_CCA1356 | DN 942 | *Symphyomyrtus* | *Adnataria* | 1187 | |
| alipes_CCA5735 | DN 5480 | *Symphyomyrtus* | *Glandulosae* | 1186 | |
| ammophila_CCA1515 | DN 1284 | *Symphyomyrtus* | *Exsertaria* | 367 | |
| amygdalina_CCA5862 | DN 6722 | *Eucalyptus* | *Eucalyptus* | 1187 | |
| amygdalina_CCA5863 | DN 6722 | *Eucalyptus* | *Eucalyptus* | 1186 | technical replicates |
| ANBG9806169_Ferguson2024 | | *Eucalyptus* | *Eucalyptus* | 1185 | not CCA |
| angulosa_CCA1155 | DN 1136 | *Symphyomyrtus* | *Dumaria* | 1187 | label fix (previous: CCA1315) |
| angulosa_CCA1313 | DN 939 | *Symphyomyrtus* | *Dumaria* | 1184 | label fix (previous: CCA1156); technical replicates |
| angulosa_CCA1315 | DN 939 | *Symphyomyrtus* | *Dumaria* | 1187 | label fix (previous: CCA1155) |
| angustissima_CCA0124 | DN 150 | *Symphyomyrtus* | *Bisectae* | 1140 | technical replicates |
| angustissima_CCA0127 | DN 150 | *Symphyomyrtus* | *Bisectae* | 1186 | |
| annulata_CCA0310 | DN 206 | *Symphyomyrtus* | *Glandulosae* | 1186 | |
| aparrerinja_CCA2829 | | *Blakella* | *Abbreviatae* | 1144 | technical replicates |
| aparrerinja_CCA2832 | | *Blakella* | *Abbreviatae* | 1163 | |
| apocynifolia_apocynifolia_CCA3247 | DN 2073 | *Angophora* | | 1150 | technical replicates |
| apocynifolia_apocynifolia_CCA3248 | DN 2073 | *Angophora* | | 1151 | |
| apodophylla_CCA5604 | DN 2456 | *Symphyomyrtus* | *Exsertaria* | 1186 | technical replicates |
| apodophylla_CCA5605 | DN 2456 | *Symphyomyrtus* | *Exsertaria* | 1186 | |
| apothalassica_CCA1573 | DN 1263 | *Eucalyptus* | *Amentum* | 1187 | |
| arachnaea_arachnaea_CCA0414 | DN 296 | *Symphyomyrtus* | *Glandulosae* | 1187 | |
| arachnaea_arrecta_CCA5765 | DN 5556 | *Symphyomyrtus* | *Glandulosae* | 1186 | |

| | | | | | |
|---|---|---|---|---|---|
| arachnaea_arrecta_CCA5767 | DN 5556 | *Symphyomyrtus* | *Glandulosae* | 1187 | technical replicates |
| arachnaea_x_redunca_pluricaulis_CCA0448 | DN 251 | *Symphyomyrtus* | *Glandulosae* | 1184 | |
| arborella_CCA1236 | DN 1127 | *Symphyomyrtus* | *Glandulosae* | 1186 | label fix (previous: CCA1246) |
| arctata_CCA0028 | DN 291 | *Symphyomyrtus* | *Bisectae* | 1183 | |
| arctata_CCA0029 | DN 291 | *Symphyomyrtus* | *Bisectae* | 1186 | technical replicates |
| arctata_CCA0314 | DN 276 | *Symphyomyrtus* | *Bisectae* | 1186 | |
| arenacea_CCA2955 | DN 1588 | *Eucalyptus* | *Eucalyptus* | 1187 | |
| argutifolia_CCA0160 | DN 249 | *Symphyomyrtus* | *Dumaria* | 1186 | |
| aridimontana_CCA5792 | DN 5778 | *Symphyomyrtus* | *Adnataria* | 1187 | |
| aridimontana_CCA5793 | DN 5778 | *Symphyomyrtus* | *Adnataria* | 1187 | technical replicates |
| aridimontana_CCA5849 | DN 5766 | *Symphyomyrtus* | *Adnataria* | 1187 | |
| armillata_CCA0412 | DN 287 | *Symphyomyrtus* | *Glandulosae* | 1187 | |
| arnhemensis_CCA5693 | DN 2484 | *Corymbia* | *Notiales* | 1165 | |
| aspersa_CCA2993 | DN 1890 | *Symphyomyrtus* | *Bisectae* | 1186 | putative contamination / mislabelling |
| aspratilis_CCA0490 | DN 328 | *Symphyomyrtus* | *Glandulosae* | 1187 | |
| aspratilis_CCA5995 | DN 7001 | *Symphyomyrtus* | *Glandulosae* | 1185 | technical replicates |
| aspratilis_CCA5996 | DN 7001 | *Symphyomyrtus* | *Glandulosae* | 1186 | |
| assimilans_CCA2867 | DN 1826 | *Symphyomyrtus* | *Dumaria* | 1187 | technical replicates |
| assimilans_CCA2868 | DN 1826 | *Symphyomyrtus* | *Dumaria* | 1187 | |
| astringens_merleae_CCA0605 | DN 194 | *Symphyomyrtus* | *Glandulosae* | 1187 | |
| astringens_redacta_CCA1455 | DN 214 | *Symphyomyrtus* | *Glandulosae* | 1185 | |
| astringens_redacta_CCA1456 | DN 214 | *Symphyomyrtus* | *Glandulosae* | 1185 | technical replicates |
| atrata_CCA1619 | DN 1308 | *Symphyomyrtus* | *Adnataria* | 1186 | technical replicates |
| atrata_CCA1620 | DN 1308 | *Symphyomyrtus* | *Adnataria* | 1186 | |
| baileyana_CCA0813 | DN 665 | *Eudesmia* | *Reticulatae* | 1171 | |

| | | | | | |
|---|---|---|---|---|---|
| baiophylla_CCA1271 | DN 1181 | *Symphyomyrtus* | *Glandulosae* | 1186 | label fix (previous: CCA1193) |
| bakeri_CCA1586 | DN 1269 | *Symphyomyrtus* | *Bisectae* | 1183 | |
| balladoniensis_balladoniensis_CCA0459 | DN 118 | *Symphyomyrtus* | *Bisectae* | 1186 | putative contamination / mislabelling |
| bancroftii_CCA1684 | DN 1241 | *Symphyomyrtus* | *Exsertaria* | 1187 | |
| baudiniana_CCA5808 | DN 5577 | *Symphyomyrtus* | *Dumaria* | 1186 | technical replicates |
| baudiniana_CCA5811 | DN 5577 | *Symphyomyrtus* | *Dumaria* | 1187 | |
| baudiniana_x_oraria_intergrade_CCA5634 | DN 5575 | *Symphyomyrtus* | *Dumaria* | 1185 | |
| baudiniana_x_oraria_intergrade_CCA5636 | DN 5575 | *Symphyomyrtus* | *Dumaria* | 1185 | technical replicates |
| baueriana_CCA1632 | DN 1252 | *Symphyomyrtus* | *Adnataria* | 1187 | |
| baxteri_CCA5918 | DN 7234 | *Eucalyptus* | *Eucalyptus* | 1174 | technical replicates |
| baxteri_CCA5920 | DN 7234 | *Eucalyptus* | *Eucalyptus* | 1187 | |
| baxteri_CCA5921 | DN 7234 | *Eucalyptus* | *Eucalyptus* | 1186 | technical replicates |
| beardiana_CCA5727 | DN 5708 | *Symphyomyrtus* | *Bisectae* | 1187 | technical replicates |
| beardiana_CCA5728 | DN 5708 | *Symphyomyrtus* | *Bisectae* | 1186 | |
| behriana_CCA1333 | DN 783 | *Symphyomyrtus* | *Adnataria* | 1187 | technical replicates |
| behriana_CCA1335 | DN 783 | *Symphyomyrtus* | *Adnataria* | 1182 | label fix (previous: CCA1144) |
| behriana_x_largiflorens_CCA1404 | DN 2065 | *Symphyomyrtus* | *Adnataria* | 1187 | |
| bicostata_CCA0137 | DN 468 | *Symphyomyrtus* | *Maidenaria* | 1170 | technical replicates |
| bicostata_CCA1698 | DN 468 | *Symphyomyrtus* | *Maidenaria* | 1186 | |
| blakelyi_CCA0758 | DN 722 | *Symphyomyrtus* | *Exsertaria* | 1143 | |
| blaxellii_CCA0395 | DN 272 | *Symphyomyrtus* | *Glandulosae* | 1186 | technical replicates |
| blaxellii_CCA0396 | DN 272 | *Symphyomyrtus* | *Glandulosae* | 1187 | |
| bosistoana_CCA3041 | DN 1733 | *Symphyomyrtus* | *Adnataria* | 1185 | |
| botryoides_CCA0671 | | *Symphyomyrtus* | *Latoangulatae* | 1181 | putative contamination / mislabelling |
| botryoides_CCA3019 | DN 1739 | *Symphyomyrtus* | *Latoangulatae* | 1187 | |

| | | | | | |
|---|---|---|---|---|---|
| botryoides_CCA4939 | DN 6307 | *Symphyomyrtus* | *Latoangulatae* | 1169 | |
| brachycalyx_CCA0553 | DN 553 | *Symphyomyrtus* | *Dumaria* | 1187 | |
| brandiana_CCA5701 | DN 5514 | *Symphyomyrtus* | *Dumaria* | 1187 | |
| brandiana_Ferguson2024 | | *Symphyomyrtus* | *Dumaria* | 1187 | not CCA |
| brassiana_CCA1685 | DN 1316 | *Symphyomyrtus* | *Exsertaria* | 1187 | |
| brevipes_CCA0437 | DN 299 | *Symphyomyrtus* | *Adnataria* | 1185 | |
| brevipes_CCA0438 | DN 299 | *Symphyomyrtus* | *Adnataria* | 1185 | technical replicates |
| brevistylis_CCA1374 | DN 1141 | *Eucalyptus* | *Longistylus* | 1186 | |
| bridgesiana_CCA0681 | DN 760 | *Symphyomyrtus* | *Maidenaria* | 1181 | |
| bridgesiana_CCA0682 | DN 760 | *Symphyomyrtus* | *Maidenaria* | 1186 | technical replicates |
| brockwayi_CCA0175 | DN 136 | *Symphyomyrtus* | *Bisectae* | 1035 | technical replicates |
| brockwayi_CCA0178 | DN 136 | *Symphyomyrtus* | *Bisectae* | 1186 | |
| brownii_CCA1587 | DN 1293 | *Symphyomyrtus* | *Adnataria* | 1185 | |
| brunnea_CCA0164 | DN 669 | *Symphyomyrtus* | *Maidenaria* | 1181 | |
| burracoppinensis_CCA0378 | DN 306 | *Symphyomyrtus* | *Bisectae* | 1187 | |
| cadens_CCA2949 | DN 1772 | *Symphyomyrtus* | *Maidenaria* | 1187 | |
| caesia_boodjin_CCA5666 | DN 5542 | *Symphyomyrtus* | *Bisectae* | 1186 | |
| caesia_boodjin_CCA5668 | DN 5542 | *Symphyomyrtus* | *Bisectae* | 1187 | technical replicates |
| caesia_caesia_CCA0321 | DN 308 | *Symphyomyrtus* | *Bisectae* | 1187 | |
| caesia_caesia_CCA0324 | DN 308 | *Symphyomyrtus* | *Bisectae* | 1187 | technical replicates |
| caesia_magna_CCA0658 | DN 304 | *Symphyomyrtus* | *Bisectae* | 1187 | |
| caesia_magna_CCA5673 | DN 5455 | *Symphyomyrtus* | *Bisectae* | 1187 | technical replicates |
| caesia_magna_CCA5674 | DN 5455 | *Symphyomyrtus* | *Bisectae* | 1187 | |
| caesia_magna_CCA5721 | DN 5610 | *Symphyomyrtus* | *Bisectae* | 1187 | |
| caesia_magna_CCA5722 | DN 5610 | *Symphyomyrtus* | *Bisectae* | 1187 | technical replicates |

| | | | | | |
|---|---|---|---|---|---|
| caesia_Southern_rocks_CCA5820 | DN 5550 | *Symphyomyrtus* | *Bisectae* | 1187 | |
| calcareana_CCA0368 | DN 103 | *Symphyomyrtus* | *Dumaria* | 1187 | |
| calcareana_CCA0482 | DN 60 | *Symphyomyrtus* | *Dumaria* | 1187 | |
| calcareana_CCA1156 | DN 1074 | *Symphyomyrtus* | *Dumaria* | 1187 | label fix (previous: CCA1313); technical replicates |
| calcareana_CCA1157 | DN 1074 | *Symphyomyrtus* | *Dumaria* | 1187 | label fix (previous: CCA1311) |
| calcicola_unita_CCA2965 | DN 1878 | *Eucalyptus* | *Frutices* | 1186 | |
| calcicola_unita_CCA2968 | DN 1878 | *Eucalyptus* | *Frutices* | 1187 | technical replicates |
| caleyi_CCA0708 | DN 746 | *Symphyomyrtus* | *Adnataria* | 1176 | |
| caleyi_CCA1421 | DN 2102 | *Symphyomyrtus* | *Adnataria* | 1187 | |
| caleyi_Ferguson2024 | | *Symphyomyrtus* | *Adnataria* | 1187 | not CCA |
| caliginosa_CCA2994 | DN 1249 | *Eucalyptus* | *Eucalyptus* | 1187 | putative contamination / mislabelling |
| calophylla_CCA1290 | DN 1160 | *Corymbia* | '*Calophyllae*' | 1078 | |
| calophylla_CCA5889 | DN 6707 | *Corymbia* | '*Calophyllae*' | 1172 | technical replicates |
| calophylla_CCA5890 | DN 6707 | *Corymbia* | '*Calophyllae*' | 1167 | |
| calophylla_Ferguson2024 | | *Corymbia* | '*Calophyllae*' | 1167 | not CCA |
| calycogona_calycogona_CCA1220 | DN 1112 | *Symphyomyrtus* | *Adnataria* | 1186 | label fix (previous: CCA1262) |
| calycogona_calycogona_CCA1222 | DN 1112 | *Symphyomyrtus* | *Adnataria* | 1186 | label fix (previous: CCA1260); technical replicates |
| calycogona_calycogona_CCA5772 | DN 4558 | *Symphyomyrtus* | *Adnataria* | 1187 | |
| calycogona_trachybasis_CCA0674 | DN 732 | *Symphyomyrtus* | *Adnataria* | 1181 | |
| camaldulensis_arida_CCA1135 | DN 1206 | *Symphyomyrtus* | *Exsertaria* | 1180 | technical replicates |
| camaldulensis_arida_CCA1136 | DN 1206 | *Symphyomyrtus* | *Exsertaria* | 1187 | label fix (previous: CCA1343) |
| camaldulensis_arida_CCA4034 | DN 4177 | *Symphyomyrtus* | *Exsertaria* | 1047 | |
| camaldulensis_arida_CCA5622 | DN 5571 | *Symphyomyrtus* | *Exsertaria* | 1187 | technical replicates |
| camaldulensis_arida_CCA5623 | DN 5571 | *Symphyomyrtus* | *Exsertaria* | 1187 | |
| camaldulensis_arida_CCA5677 | DN 5077 | *Symphyomyrtus* | *Exsertaria* | 1187 | technical replicates |

| | | | | | |
|---|---|---|---|---|---|
| camaldulensis_arida_CCA5680 | DN 5077 | *Symphyomyrtus* | *Exsertaria* | 1187 | |
| camaldulensis_camaldulensis_1_CCA4957 | DN 4733 | *Symphyomyrtus* | *Exsertaria* | 1169 | technical replicates |
| camaldulensis_camaldulensis_2_CCA4957 | DN 4733 | *Symphyomyrtus* | *Exsertaria* | 1153 | |
| camaldulensis_camaldulensis_CCA1266 | DN 823 | *Symphyomyrtus* | *Exsertaria* | 1154 | technical replicates |
| camaldulensis_camaldulensis_CCA1267 | DN 823 | *Symphyomyrtus* | *Exsertaria* | 1169 | technical replicates |
| camaldulensis_camaldulensis_CCA1268 | DN 823 | *Symphyomyrtus* | *Exsertaria* | 1121 | |
| camaldulensis_camaldulensis_CCA1269 | DN 823 | *Symphyomyrtus* | *Exsertaria* | 1186 | label fix (previous: CCA1197); technical replicates |
| camaldulensis_Ferguson2024 | | *Symphyomyrtus* | *Exsertaria* | 1186 | not CCA |
| camaldulensis_minima_CCA5430 | DN 5233 | *Symphyomyrtus* | *Exsertaria* | 1078 | |
| camaldulensis_obtusa_CCA3451 | DN 2489 | *Symphyomyrtus* | *Exsertaria* | 1155 | |
| camaldulensis_refulgens_CCA5651 | DN 5747 | *Symphyomyrtus* | *Exsertaria* | 1187 | |
| camaldulensis_refulgens_CCA5652 | DN 5747 | *Symphyomyrtus* | *Exsertaria* | 1187 | technical replicates |
| camaldulensis_simulata_CCA1644 | DN 1314 | *Symphyomyrtus* | *Exsertaria* | 1168 | |
| camaldulensis_simulata_CCA1645 | DN 1314 | *Symphyomyrtus* | *Exsertaria* | 1187 | technical replicates |
| cameronii_CCA5981 | DN 709 | *Eucalyptus* | *Eucalyptus* | 1187 | |
| campaspe_CCA6048 | DN 5660 | *Symphyomyrtus* | *Glandulosae* | 1187 | |
| camphora_camphora_CCA0214 | DN 616 | *Symphyomyrtus* | *Maidenaria* | 1187 | |
| canaliculata_CCA1701 | DN 1232 | *Symphyomyrtus* | *Pumilio* | 1187 | |
| candida_CCA6067 | DN 7230 | *Blakella* | *Abbreviatae* | 1153 | putative contamination / mislabelling |
| canescens_beadellii_CCA1468 | DN 1506 | *Symphyomyrtus* | *Dumaria* | 1186 | |
| canescens_canescens_CCA0508 | DN 487 | *Symphyomyrtus* | *Dumaria* | 1186 | |
| canobolensis_CCA1615 | DN 1222 | *Symphyomyrtus* | *Maidenaria* | 1187 | |
| capillosa_capillosa_CCA0172 | DN 300 | *Symphyomyrtus* | *Glandulosae* | 1186 | |
| capillosa_capillosa_CCA0174 | DN 300 | *Symphyomyrtus* | *Glandulosae* | 1186 | technical replicates |
| capillosa_capillosa_CCA0252 | DN 241 | *Symphyomyrtus* | *Glandulosae* | 1187 | |

| | | | | | |
|---|---|---|---|---|---|
| capillosa_nigrifunda_CCA0513 | DN 533 | *Symphyomyrtus* | *Glandulosae* | 1187 | |
| capitanea_CCA1161 | DN 1076 | *Symphyomyrtus* | *Dumaria* | 1187 | label fix (previous: CCA1310); technical replicates |
| capitanea_CCA1162 | DN 1076 | *Symphyomyrtus* | *Dumaria* | 1187 | label fix (previous: CCA1307) |
| captiosa_CCA0232 | DN 203 | *Symphyomyrtus* | *Dumaria* | 1187 | |
| carnei_CCA0503 | DN 547 | *Symphyomyrtus* | *Glandulosae* | 1187 | |
| celastroides_CCA0618 | DN 132 | *Symphyomyrtus* | *Adnataria* | 1185 | |
| cerasiformis_CCA0356 | DN 329 | *Symphyomyrtus* | *Glandulosae* | 1187 | |
| ceratocorys_CCA6018 | DN 6943 | *Symphyomyrtus* | *Dumaria* | 1187 | technical replicates |
| ceratocorys_CCA6021 | DN 6943 | *Symphyomyrtus* | *Dumaria* | 1185 | |
| cernua_CCA0607 | DN 182 | *Symphyomyrtus* | *Glandulosae* | 1186 | |
| chartaboma_CCA5408 | DN 5143 | *Eudesmia* | *Aurantistamineae* | 1126 | |
| chippendalei_CCA5826 | DN 4288 | *Corymbia* | *Notiales* | 1166 | |
| chloroclada_CCA5893 | DN 6464 | *Symphyomyrtus* | *Exsertaria* | 1186 | technical replicates |
| chloroclada_CCA5896 | DN 6464 | *Symphyomyrtus* | *Exsertaria* | 1186 | |
| chlorolampra_CCA5938 | DN 7227 | *Corymbia* | ‘*Calophyllae*’ | 1167 | |
| chlorophylla_CCA1499 | DN 1334 | *Symphyomyrtus* | *Adnataria* | 1184 | |
| chlorophylla_CCA1649 | DN 1315 | *Symphyomyrtus* | *Adnataria* | 1186 | |
| cinerea_cinerea_CCA4706 | DN 3211 | *Symphyomyrtus* | *Maidenaria* | 1186 | |
| citriodora_CCA0866 | DN 702 | *Blakella* | ‘*Maculatae*’ | 1110 | technical replicates |
| citriodora_CCA2823 | DN 702 | *Blakella* | ‘*Maculatae*’ | 1162 | technical replicates |
| citriodora_CCA2825 | DN 702 | *Blakella* | ‘*Maculatae*’ | 1163 | |
| cladocalyx_crassa_CCA1355 | DN 1053 | *Symphyomyrtus* | *Sejunctae* | 1187 | |
| cladocalyx_Ferguson2024 | | *Symphyomyrtus* | *Sejunctae* | 1186 | not CCA |
| clelandiorum_CCA0067 | DN 333 | *Symphyomyrtus* | *Dumaria* | 1187 | |
| cliftoniana_CCA5773 | DN 5851 | *Corymbia* | *Notiales* | 1138 | |

| | | | | | |
|---|---|---|---|---|---|
| cloeziana_CCA0904 | DN 696 | *Idiogenes* | | 1179 | |
| cloeziana_Ferguson2024 | | *Idiogenes* | | 1185 | not CCA |
| cneorifolia_CCA0615 | DN 75 | *Symphyomyrtus* | *Bisectae* | 1160 | technical replicates |
| cneorifolia_CCA0616 | DN 75 | *Symphyomyrtus* | *Bisectae* | 1187 | |
| codonocarpa_CCA5950 | DN 5025 | *Eucalyptus* | *Eucalyptus* | 1186 | technical replicates |
| codonocarpa_CCA5951 | DN 5025 | *Eucalyptus* | *Eucalyptus* | 1186 | |
| cometae-vallis_CCA2847 | DN 1897 | *Symphyomyrtus* | *Dumaria* | 1186 | |
| concinna_CCA1488 | DN 1391 | *Symphyomyrtus* | *Dumaria* | 804 | |
| confluens_CCA6075 | DN 1908 | *Symphyomyrtus* | *Platysperma* | 1186 | |
| conglobata_conglobata_CCA0550 | DN 559 | *Symphyomyrtus* | *Dumaria* | 1176 | |
| conglobata_conglobata_CCA1329 | DN 944 | *Symphyomyrtus* | *Dumaria* | 1159 | label fix (previous: CCA1130) |
| conglobata_conglobata_CCA1332 | DN 944 | *Symphyomyrtus* | *Dumaria* | 1187 | technical replicates |
| conglobata_conglobata_CCA1339 | DN 938 | *Symphyomyrtus* | *Dumaria* | 1181 | label fix (previous: CCA1143) |
| conglobata_conglobata_CCA1349 | DN 933 | *Symphyomyrtus* | *Dumaria* | 1186 | |
| conica_CCA0677 | DN 747 | *Symphyomyrtus* | *Adnataria* | 1172 | technical replicates |
| conica_CCA0678 | DN 747 | *Symphyomyrtus* | *Adnataria* | 1186 | |
| conica_x_microcarpa_CCA0680 | DN 747 | *Symphyomyrtus* | *Adnataria* | 1186 | technical replicates |
| connexa_CCA0331 | DN 162 | *Symphyomyrtus* | *Bisectae* | 1185 | technical replicates |
| connexa_CCA0334 | DN 162 | *Symphyomyrtus* | *Bisectae* | 1186 | |
| coolabah_CCA1478 | DN 1355 | *Symphyomyrtus* | *Adnataria* | 1069 | technical replicates |
| coolabah_CCA1479 | DN 1355 | *Symphyomyrtus* | *Adnataria* | 1077 | |
| coolabah_CCA1480 | DN 1355 | *Symphyomyrtus* | *Adnataria* | 891 | technical replicates |
| coolabah_CCA2828 | | *Symphyomyrtus* | *Adnataria* | 1187 | |
| coolabah_CCA3711 | DN 2957 | *Symphyomyrtus* | *Adnataria* | 1070 | |
| coolabah_CCA4029 | DN 4186 | *Symphyomyrtus* | *Adnataria* | 1080 | |

| | | | | | |
|---|---|---|---|---|---|
| coolabah_Ferguson2024 | | *Symphyomyrtus* | *Adnataria* | 1179 | not CCA |
| cooperiana_CCA0340 | DN 168 | *Symphyomyrtus* | *Bisectae* | 1187 | |
| cooperiana_CCA2974 | DN 1813 | *Symphyomyrtus* | *Bisectae* | 1187 | technical replicates |
| cooperiana_CCA2975 | DN 1813 | *Symphyomyrtus* | *Bisectae* | 1187 | |
| cordata_quadrangulosa_CCA5977 | DN 1979 | *Symphyomyrtus* | *Maidenaria* | 1187 | technical replicates |
| cordata_quadrangulosa_CCA5980 | DN 1979 | *Symphyomyrtus* | *Maidenaria* | 1187 | |
| cornuta_CCA0460 | DN 173 | *Symphyomyrtus* | *Glandulosae* | 1186 | putative contamination / mislabelling |
| coronata_CCA3471 | DN 2247 | *Eucalyptus* | *Frutices* | 1064 | |
| corrugata_CCA0154 | DN 305 | *Symphyomyrtus* | *Dumaria* | 1187 | |
| corynodes_CCA1590 | DN 1264 | *Symphyomyrtus* | *Adnataria* | 1185 | |
| cosmophylla_CCA6033 | DN 7236 | *Symphyomyrtus* | *Incognitae* | 1171 | technical replicates |
| cosmophylla_CCA6034 | DN 7236 | *Symphyomyrtus* | *Incognitae* | 1187 | technical replicates |
| cosmophylla_CCA6035 | DN 7236 | *Symphyomyrtus* | *Incognitae* | 1187 | |
| crebra_CCA4821 | DN 6453 | *Symphyomyrtus* | *Adnataria* | 1139 | |
| cretata_CCA2873 | DN 1689 | *Symphyomyrtus* | *Dumaria* | 1187 | |
| croajingolensis_CCA3056 | DN 1744 | *Eucalyptus* | *Eucalyptus* | 1133 | |
| crucis_lanceolata_CCA0260 | DN 303 | *Symphyomyrtus* | *Bisectae* | 1187 | |
| crucis_lanceolata_CCA5775 | DN 5456 | *Symphyomyrtus* | *Bisectae* | 1185 | |
| crucis_lanceolata_CCA5776 | DN 5456 | *Symphyomyrtus* | *Bisectae* | 1186 | technical replicates |
| crucis_lanceolata_CCA5796 | DN 5603 | *Symphyomyrtus* | *Bisectae* | 1187 | |
| crucis_praecipua_CCA2925 | DN 1561 | *Symphyomyrtus* | *Bisectae* | 1187 | |
| cullenii_CCA1676 | DN 1322 | *Symphyomyrtus* | *Adnataria* | 1187 | |
| cupularis_CCA2946 | DN 1938 | *Symphyomyrtus* | *Exsertaria* | 1187 | |
| curtisii_CCA3261 | DN 2108 | *Acerosae* | | 1162 | |
| curtisii_Ferguson2024 | | *Acerosae* | | 1185 | not CCA |

| | | | | | |
|---|---|---|---|---|---|
| cyclostoma_CCA0092 | DN 125 | *Symphyomyrtus* | *Dumaria* | 1187 | |
| cylindriflora_CCA0226 | DN 144 | *Symphyomyrtus* | *Glandulosae* | 1187 | |
| cylindrocarpa_CCA0351 | DN 121 | *Symphyomyrtus* | *Dumaria* | 1187 | |
| cylindrocarpa_CCA0493 | DN 526 | *Symphyomyrtus* | *Dumaria* | 1186 | |
| cypellocarpa_CCA1270 | DN 356 | *Symphyomyrtus* | *Maidenaria* | 1187 | label fix (previous: CCA1196) |
| cypellocarpa_CCA3837 | DN 2946 | *Symphyomyrtus* | *Maidenaria* | 1168 | |
| dalrympleana_heptantha_CCA5613 | DN 5022 | *Symphyomyrtus* | *Maidenaria* | 1187 | |
| dalrympleana_heptantha_CCA5614 | DN 5022 | *Symphyomyrtus* | *Maidenaria* | 1187 | technical replicates |
| dalrympleana_lutruwita_CCA5866 | DN 6736 | *Symphyomyrtus* | *Maidenaria* | 1187 | technical replicates |
| dalrympleana_lutruwita_CCA5867 | DN 6736 | *Symphyomyrtus* | *Maidenaria* | 1187 | |
| dawsonii_CCA0687 | DN 743 | *Symphyomyrtus* | *Adnataria* | 1176 | technical replicates |
| dawsonii_CCA0688 | DN 743 | *Symphyomyrtus* | *Adnataria* | 1185 | |
| dawsonii_Ferguson2024 | | *Symphyomyrtus* | *Adnataria* | 1186 | not CCA |
| dealbata_dealbata_CCA0761 | DN 723 | *Symphyomyrtus* | *Exsertaria* | 1171 | |
| deanei_CCA0138 | DN 633 | *Symphyomyrtus* | *Maidenaria* | 1187 | |
| decipiens_CCA0375 | DN 227 | *Symphyomyrtus* | *Bisectae* | 1056 | technical replicates |
| decipiens_CCA0376 | DN 227 | *Symphyomyrtus* | *Bisectae* | 1187 | |
| decipiens_CCA1310 | DN 1153 | *Symphyomyrtus* | *Bisectae* | 1186 | label fix (previous: CCA1161) |
| decipiens_CCA1311 | DN 1153 | *Symphyomyrtus* | *Bisectae* | 1180 | label fix (previous: CCA1157); technical replicates |
| decipiens_Ferguson2024 | | *Symphyomyrtus* | *Bisectae* | 1186 | not CCA |
| decorticans_CCA1598 | DN 1260 | *Symphyomyrtus* | *Adnataria* | 1174 | putative contamination / mislabelling |
| deflexa_CCA1229 | DN 1110 | *Symphyomyrtus* | *Dumaria* | 1187 | label fix (previous: CCA1254) |
| delicata_CCA0266 | DN 149 | *Symphyomyrtus* | *Bisectae* | 960 | technical replicates |
| delicata_CCA0267 | DN 149 | *Symphyomyrtus* | *Bisectae* | 1187 | |

| | | | | | |
|---|---|---|---|---|---|
| dendromorpha_CCA5858 | DN 5869 | *Eucalyptus* | *Eucalyptus* | 1187 | |
| deserticola_deserticola_CCA1413 | DN 542 | *Corymbia* | *Notiales* | 1157 | |
| deserticola_deserticola_CCA5816 | DN 5750 | *Corymbia* | *Notiales* | 1164 | |
| deserticola_deserticola_CCA5819 | DN 5750 | *Corymbia* | *Notiales* | 1164 | technical replicates |
| desmondensis_CCA0235 | DN 192 | *Symphyomyrtus* | *Glandulosae* | 1187 | |
| desmondensis_x_occidentalis_CCA0234 | DN 192 | *Symphyomyrtus* | *Glandulosae* | 1187 | technical replicates |
| deuaensis_CCA3039 | DN 1769 | *Eucalyptus* | *Eucalyptus* | 1187 | |
| dielsii_CCA1204 | DN 1120 | *Symphyomyrtus* | *Glandulosae* | 1187 | label fix (previous: CCA1202) |
| dielsii_CCA5707 | DN 5472 | *Symphyomyrtus* | *Glandulosae* | 1187 | |
| diminuta_CCA0408 | DN 271 | *Symphyomyrtus* | *Glandulosae* | 1187 | |
| diminuta_CCA0418 | DN 271 | *Symphyomyrtus* | *Glandulosae* | 1187 | technical replicates |
| diminuta_CCA5689 | DN 5543 | *Symphyomyrtus* | *Glandulosae* | 1187 | technical replicates |
| diminuta_CCA5690 | DN 5543 | *Symphyomyrtus* | *Glandulosae* | 1187 | |
| diminuta_CCA5691 | DN 5543 | *Symphyomyrtus* | *Glandulosae* | 1187 | technical replicates |
| diptera_CCA0302 | DN 146 | *Symphyomyrtus* | *Glandulosae* | 1187 | |
| dissimulata_dissimulata_CCA0317 | DN 205 | *Symphyomyrtus* | *Bisectae* | 1186 | |
| dissimulata_dissimulata_CCA2945 | DN 1629 | *Symphyomyrtus* | *Bisectae* | 1187 | |
| diversicolor_CCA1370 | DN 1142 | *Symphyomyrtus* | *Bisectae* | 1171 | |
| diversifolia_hesperia_CCA2920 | DN 1811 | *Eucalyptus* | *Frutices* | 1186 | |
| dives_CCA5180 | DN 6099 | *Eucalyptus* | *Eucalyptus* | 1175 | |
| dolichocera_CCA0430 | DN 273 | *Symphyomyrtus* | *Bisectae* | 1185 | |
| dorrigoensis_CCA5871 | DN 6798 | *Symphyomyrtus* | *Maidenaria* | 1187 | |
| dorrigoensis_x_saligna_CCA5874 | DN 6798 | *Symphyomyrtus* | | 1184 | inter-sectional hybrid |
| drepanophylla_CCA1568 | DN 1294 | *Symphyomyrtus* | *Adnataria* | 1187 | |
| drepanophylla_CCA1651 | DN 1317 | *Symphyomyrtus* | *Adnataria* | 1186 | |

| | | | | | |
|---|---|---|---|---|---|
| drepanophylla_CCA5962 | DN 2531 | *Symphyomyrtus* | *Adnataria* | 1187 | |
| drepanophylla_CCA5964 | DN 2531 | *Symphyomyrtus* | *Adnataria* | 1186 | technical replicates |
| drummondii_CCA1165 | DN 1156 | *Symphyomyrtus* | *Bisectae* | 1186 | label fix (previous: CCA1298) |
| drummondii_CCA3009 | DN 1892 | *Symphyomyrtus* | *Bisectae* | 1185 | |
| dumosa_CCA0485 | DN 93 | *Symphyomyrtus* | *Dumaria* | 1187 | |
| dundasii_CCA0061 | DN 129 | *Symphyomyrtus* | *Glandulosae* | 1187 | putative contamination / mislabelling |
| dunnii_CCA1680 | DN 1257 | *Symphyomyrtus* | *Maidenaria* | 1130 | |
| dwyeri_CCA1580 | DN 1369 | *Symphyomyrtus* | *Exsertaria* | 1187 | |
| ebbanoensis_ebbanoensis_CCA0037 | DN 293 | *Eudesmia* | *Limbatae* | 1119 | technical replicates |
| ebbanoensis_ebbanoensis_CCA0038 | DN 293 | *Eudesmia* | *Limbatae* | 1120 | technical replicates |
| ebbanoensis_ebbanoensis_CCA0039 | DN 293 | *Eudesmia* | *Limbatae* | 1179 | |
| ebbanoensis_photina_CCA0026 | DN 270 | *Eudesmia* | *Limbatae* | 1179 | |
| ecdysiastes_CCA0498 | DN 527 | *Symphyomyrtus* | *Bisectae* | 1184 | |
| ecdysiastes_CCA5620 | DN 5513 | *Symphyomyrtus* | *Bisectae* | 1185 | |
| ecdysiastes_CCA5621 | DN 5513 | *Symphyomyrtus* | *Bisectae* | 1187 | technical replicates |
| ecdysiastes_CCA6022 | DN 6945 | *Symphyomyrtus* | *Bisectae* | 1187 | |
| ecdysiastes_x_horistes _intergrade_CCA6030 | DN 6951 | *Symphyomyrtus* | *Bisectae* | 1186 | |
| ecdysiastes_x_horistes _intergrade_CCA6031 | DN 6951 | *Symphyomyrtus* | *Bisectae* | 1185 | technical replicates |
| ecostata_CCA0233 | DN 198 | *Symphyomyrtus* | *Bisectae* | 1187 | |
| ecostata_CCA5738 | DN 5496 | *Symphyomyrtus* | *Bisectae* | 1187 | |
| educta_CCA0083 | DN 335 | *Symphyomyrtus* | *Bisectae* | 1187 | |
| educta_CCA6028 | DN 6928 | *Symphyomyrtus* | *Bisectae* | 1187 | |
| educta_CCA6053 | DN 6921 | *Symphyomyrtus* | *Bisectae* | 1187 | |
| effusa_effusa_CCA0095 | DN 127 | *Symphyomyrtus* | *Glandulosae* | 1187 | |
| effusa_exsul_CCA1531 | DN 1554 | *Symphyomyrtus* | *Glandulosae* | 1186 | |

| | | | | | |
|---|---|---|---|---|---|
| elata_CCA3147 | DN 1742 | *Eucalyptus* | *Eucalyptus* | 1141 | |
| elata_CCA3893 | DN 2901 | *Eucalyptus* | *Eucalyptus* | 1180 | |
| elata_CCA5884 | DN 6783 | *Eucalyptus* | *Eucalyptus* | 1187 | technical replicates |
| elata_CCA5885 | DN 6783 | *Eucalyptus* | *Eucalyptus* | 1187 | |
| eremaea_eremaea_CCA2909 | DN 1708 | *Corymbia* | *Notiales* | 1156 | |
| eremaea_oligocarpa_CCA1133 | DN 1208 | *Corymbia* | *Notiales* | 1157 | label fix (previous: CCA1325); technical replicates |
| eremaea_oligocarpa_CCA6079 | DN 1208 | *Corymbia* | *Notiales* | 1167 | |
| eremicola_eremicola_CCA6057 | DN 1500 | *Symphyomyrtus* | *Bisectae* | 1187 | |
| eremicola_peeneri_CCA1409 | DN 1518 | *Symphyomyrtus* | *Bisectae* | 1186 | |
| eremicola_peeneri_CCA1474 | DN 1542 | *Symphyomyrtus* | *Bisectae* | 902 | |
| eremicola_peeneri_CCA1477 | DN 1542 | *Symphyomyrtus* | *Bisectae* | 1 | low coverage |
| eremophila_CCA0204 | DN 238 | *Symphyomyrtus* | *Glandulosae* | 1186 | |
| eremophila_CCA0441 | DN 310 | *Symphyomyrtus* | *Glandulosae* | 1186 | |
| erythrocorys_CCA0584 | DN 265 | *Eudesmia* | *Limbatae* | 1184 | |
| erythrocorys_CCA1246 | DN 1127 | *Eudesmia* | *Limbatae* | 1184 | label fix (previous: CCA1236) |
| erythrocorys_CCA5853 | DN 5540 | *Eudesmia* | *Limbatae* | 1183 | technical replicates |
| erythrocorys_CCA5855 | DN 5540 | *Eudesmia* | *Limbatae* | 1184 | |
| erythrocorys_Ferguson2024 | | *Eudesmia* | *Limbatae* | 1182 | not CCA |
| erythronema_inornata_CCA0569 | DN 245 | *Symphyomyrtus* | *Glandulosae* | 1187 | |
| eudesmioides_CCA0405 | DN 279 | *Eudesmia* | *Limbatae* | 1164 | technical replicates |
| eudesmioides_CCA0406 | DN 279 | *Eudesmia* | *Limbatae* | 1182 | |
| eugenioides_CCA1629 | DN 1235 | *Eucalyptus* | *Eucalyptus* | 1187 | |
| ewartiana_CCA0435 | DN 280 | *Symphyomyrtus* | *Bisectae* | 1185 | |
| ewartiana_x_arctata_CCA0434 | DN 280 | *Symphyomyrtus* | *Bisectae* | 1186 | technical replicates |
| exigua_CCA0361 | DN 322 | *Symphyomyrtus* | *Dumaria* | 1187 | |

| | | | | | |
|---|---|---|---|---|---|
| exilipes_CCA5974 | DN 1289 | *Symphyomyrtus* | *Adnataria* | 1187 | |
| exilipes_CCA5975 | DN 1289 | *Symphyomyrtus* | *Adnataria* | 1183 | technical replicates |
| exilis_CCA5669 | DN 5541 | *Eucalyptus* | *Frutices* | 1186 | technical replicates |
| exilis_CCA5672 | DN 5541 | *Eucalyptus* | *Frutices* | 1187 | |
| eximia_CCA0837 | DN 632 | *Blakella* | ‘*Naviculares*’ | 1160 | |
| expressa_CCA5603 | DN 5266 | *Eucalyptus* | *Eucalyptus* | 1187 | |
| exserta_CCA0736 | DN 682 | *Symphyomyrtus* | *Exsertaria* | 1166 | |
| exserta_CCA1640 | DN 1261 | *Symphyomyrtus* | *Exsertaria* | 1187 | |
| exserta_CCA1670 | DN 1309 | *Symphyomyrtus* | *Exsertaria* | 1187 | |
| exserta_CCA4776 | DN 6168 | *Symphyomyrtus* | *Exsertaria* | 1105 | |
| extrica_CCA1111 | DN 1092 | *Eudesmia* | *Limbatae* | 787 | putative contamination / mislabelling |
| falcata_CCA1206 | DN 1113 | *Symphyomyrtus* | *Bisectae* | 1187 | label fix (previous: CCA1199) |
| fasciculosa_CCA2963 | DN 1716 | *Symphyomyrtus* | *Adnataria* | 1187 | |
| fastigata_CCA4877 | DN 596 | *Eucalyptus* | *Eucalyptus* | 1139 | |
| fergusonii_dorsiventralis_CCA1372 | DN 634 | *Symphyomyrtus* | *Adnataria* | 1187 | |
| fibrosa_CCA0768 | DN 666 | *Symphyomyrtus* | *Adnataria* | 1167 | |
| fibrosa_Ferguson2024 | | *Symphyomyrtus* | *Adnataria* | 1186 | not CCA |
| ficifolia_CCA5615 | DN 5523 | *Corymbia* | ‘*Calophyllae*’ | 1148 | technical replicates |
| ficifolia_CCA5616 | DN 5523 | *Corymbia* | ‘*Calophyllae*’ | 1139 | technical replicates |
| ficifolia_CCA5617 | DN 5523 | *Corymbia* | ‘*Calophyllae*’ | 1163 | |
| flavida_CCA0052 | DN 343 | *Symphyomyrtus* | *Glandulosae* | 1186 | |
| flindersii_CCA0477 | | *Symphyomyrtus* | *Exsertaria* | 1186 | technical replicates |
| flindersii_CCA0478 | | *Symphyomyrtus* | *Exsertaria* | 1185 | |
| flocktoniae_flocktoniae_CCA0424 | DN 190 | *Symphyomyrtus* | *Bisectae* | 1186 | |
| flocktoniae_flocktoniae_CCA2959 | DN 1870 | *Symphyomyrtus* | *Bisectae* | 1187 | |

| | | | | | |
|---|---|---|---|---|---|
| flocktoniae_flocktoniae_CCA2997 | DN 1847 | *Symphyomyrtus* | *Bisectae* | 1187 | |
| flocktoniae_flocktoniae_x_occidentalis_CCA2924 | DN 1881 | *Symphyomyrtus* | | 1186 | inter-sectional hybrid |
| flocktoniae_hebes_CCA2900 | DN 1828 | *Symphyomyrtus* | *Bisectae* | 1187 | |
| florida_CCA2812 | | *Angophora* | | 1148 | technical replicates |
| florida_CCA2813 | | *Angophora* | | 1154 | |
| florida_Ferguson2024 | | *Angophora* | | 1147 | not CCA |
| foliosa_CCA0211 | DN 161 | *Symphyomyrtus* | *Bisectae* | 1187 | |
| formanii_circulata_CCA5840 | DN 5672 | *Symphyomyrtus* | *Bisectae* | 1186 | technical replicates |
| formanii_circulata_CCA5842 | DN 5672 | *Symphyomyrtus* | *Bisectae* | 1187 | |
| formanii_circulata_CCA5844 | DN 5684 | *Symphyomyrtus* | *Bisectae* | 1187 | |
| formanii_formanii_CCA0667 | | *Symphyomyrtus* | *Bisectae* | 1182 | |
| formanii_formanii_CCA1538 | DN 1567 | *Symphyomyrtus* | *Bisectae* | 1186 | |
| formanii_formanii_CCA5801 | DN 5693 | *Symphyomyrtus* | *Bisectae* | 1187 | |
| forrestiana_CCA0608 | DN 160 | *Symphyomyrtus* | *Dumaria* | 1157 | |
| fraseri_fraseri_CCA0115 | DN 116 | *Symphyomyrtus* | *Dumaria* | 1187 | technical replicates |
| fraseri_fraseri_CCA0116 | DN 116 | *Symphyomyrtus* | *Dumaria* | 1187 | |
| fraseri_melanobasis_CCA0367 | DN 128 | *Symphyomyrtus* | *Dumaria* | 1187 | |
| frenchiana_CCA1405 | | *Symphyomyrtus* | *Dumaria* | 1186 | |
| froggattii_CCA0591 | DN 453 | *Symphyomyrtus* | *Adnataria* | 1187 | |
| fruticosa_CCA5787 | DN 5567 | *Symphyomyrtus* | *Bisectae* | 28 | low coverage |
| fruticosa_CCA5788 | DN 5567 | *Symphyomyrtus* | *Bisectae* | 188 | low coverage |
| fusiformis_CCA0971 | DN 645 | *Symphyomyrtus* | *Adnataria* | 1186 | |
| gamophylla_CCA0004 | | *Eudesmia* | *Limbatae* | 1178 | |
| gardneri_CCA0409 | DN 295 | *Symphyomyrtus* | *Glandulosae* | 1186 | |
| georgei_fulgida_CCA0383 | DN 323 | *Symphyomyrtus* | *Dumaria* | 1187 | |

| | | | | | |
|---|---|---|---|---|---|
| georgei_georgei_CCA0201 | DN 325 | *Symphyomyrtus* | *Dumaria* | 1186 | |
| gillenii_CCA1301 | DN 1209 | *Symphyomyrtus* | *Exsertaria* | 1186 | label fix (previous: CCA1163) |
| gillenii_CCA1302 | DN 1209 | *Symphyomyrtus* | *Exsertaria* | 1177 | technical replicates |
| gillenii_CCA2906 | DN 1703 | *Symphyomyrtus* | *Exsertaria* | 1187 | |
| gillii_CCA0670 | | *Symphyomyrtus* | *Bisectae* | 1180 | |
| gillii_CCA2913 | DN 1715 | *Symphyomyrtus* | *Bisectae* | 1185 | |
| gittinsii_gittinsii_CCA1181 | DN 1170 | *Eudesmia* | *Limbatae* | 1183 | label fix (previous: CCA1280) |
| gittinsii_gittinsii_CCA1182 | DN 1170 | *Eudesmia* | *Limbatae* | 1183 | label fix (previous: CCA1281); technical replicates |
| gittinsii_illucida_CCA0240 | DN 252 | *Eudesmia* | *Limbatae* | 1184 | putative contamination / mislabelling |
| glaucina_CCA1380 | CSIRO 11239 | *Symphyomyrtus* | *Exsertaria* | 1185 | technical replicates |
| glaucina_CCA1383 | CSIRO 11239 | *Symphyomyrtus* | *Exsertaria* | 1187 | |
| globulus_CCA3130 | DN 1726 | *Symphyomyrtus* | *Maidenaria* | 1090 | |
| globulus_CCA3216 | DN 1986 | *Symphyomyrtus* | *Maidenaria* | 1171 | |
| globulus_CCA3217 | DN 1986 | *Symphyomyrtus* | *Maidenaria* | 972 | technical replicates |
| globulus_Ferguson2024 | | *Symphyomyrtus* | *Maidenaria* | 1186 | not CCA |
| glomerosa_CCA0013 | | *Symphyomyrtus* | *Bisectae* | 1187 | |
| glomerosa_CCA1470 | DN 1508 | *Symphyomyrtus* | *Bisectae* | 1186 | |
| gomphocephala_CCA0700 | | *Symphyomyrtus* | *Bolites* | 1166 | |
| gomphocephala_CCA1202 | DN 1148 | *Symphyomyrtus* | *Bolites* | 1187 | label fix (previous: CCA1204) |
| gomphocephala_x_zopherophloia_CCA5969 | DN 7226 | *Symphyomyrtus* | | 1186 | inter-sectional hybrid |
| gomphocephala_x_zopherophloia_CCA5972 | DN 7226 | *Symphyomyrtus* | | 1186 | inter-sectional hybrid |
| gongylocarpa_CCA3997 | DN 4303 | *Eudesmia* | *Limbatae* | 1123 | |
| goniantha_goniantha_CCA2951 | DN 1626 | *Symphyomyrtus* | *Bisectae* | 1186 | |
| goniocalyx_exposa_CCA0755 | DN 565 | *Symphyomyrtus* | *Maidenaria* | 1152 | |
| goniocalyx_viridissima_CCA0620 | DN 449 | *Symphyomyrtus* | *Maidenaria* | 1187 | |

| | | | | | |
|---|---|---|---|---|---|
| goniocarpa_CCA0055 | DN 186 | *Symphyomyrtus* | *Glandulosae* | 1186 | |
| gracilis_CCA0643 | DN 444 | *Symphyomyrtus* | *Adnataria* | 1186 | |
| gracilis_CCA3004 | DN 1861 | *Symphyomyrtus* | *Adnataria* | 1187 | |
| grandis_CCA1654 | DN 1237 | *Symphyomyrtus* | *Latoangulatae* | 1170 | technical replicates |
| grandis_CCA1655 | DN 1237 | *Symphyomyrtus* | *Latoangulatae* | 1167 | technical replicates |
| grandis_CCA1656 | DN 1237 | *Symphyomyrtus* | *Latoangulatae* | 1187 | |
| grandis_Ferguson2024 | | *Symphyomyrtus* | *Latoangulatae* | 1186 | not CCA |
| granitica_CCA1657 | DN 1305 | *Symphyomyrtus* | *Adnataria* | 1031 | technical replicates |
| granitica_CCA1659 | DN 1305 | *Symphyomyrtus* | *Adnataria* | 1186 | |
| grasbyi_CCA5835 | DN 5689 | *Symphyomyrtus* | *Bisectae* | 1184 | |
| grasbyi_CCA5836 | DN 5689 | *Symphyomyrtus* | *Bisectae* | 1186 | technical replicates |
| grasbyi_CCA6000 | DN 6997 | *Symphyomyrtus* | *Bisectae* | 1185 | |
| grasbyi_CCA6001 | DN 6997 | *Symphyomyrtus* | *Bisectae* | 1186 | technical replicates |
| griffithsii_CCA0631 | DN 515 | *Symphyomyrtus* | *Dumaria* | 1186 | putative contamination / mislabelling |
| grisea_CCA1678 | DN 1271 | *Symphyomyrtus* | *Pumilio* | 1187 | |
| grossa_CCA5958 | DN 7224 | *Symphyomyrtus* | *Glandulosae* | 1187 | |
| grossa_CCA5959 | DN 7224 | *Symphyomyrtus* | *Glandulosae* | 1186 | technical replicates |
| guilfoylei_CCA5922 | DN 4632 | *Cruciformes* | | 1163 | technical replicates |
| guilfoylei_CCA5923 | DN 4632 | *Cruciformes* | | 1183 | |
| guilfoylei_CCA5925 | DN 4632 | *Cruciformes* | | 1185 | technical replicates |
| guilfoylei_Ferguson2024 | | *Cruciformes* | | 1185 | not CCA |
| gummifera_CCA0937 | | *Corymbia* | *Notiales* | 1110 | |
| gummifera_CCA1650 | DN 1228 | *Corymbia* | *Notiales* | 988 | |
| gypsophila_CCA0533 | DN 525 | *Symphyomyrtus* | *Dumaria* | 947 | |
| gypsophila_CCA1483 | DN 1405 | *Symphyomyrtus* | *Dumaria* | 1187 | |

| | | | | | |
|---|---|---|---|---|---|
| gypsophila_CCA1492 | DN 1520 | *Symphyomyrtus* | *Dumaria* | 1045 | |
| haemastoma_CCA5319 | DN 6795 | *Eucalyptus* | *Eucalyptus* | 1147 | |
| haematoxylon_CCA1294 | DN 1147 | *Corymbia* | ‘*Calophyllae*’ | 1163 | label fix (previous: CCA1173) |
| haematoxylon_CCA1296 | DN 1147 | *Corymbia* | ‘*Calophyllae*’ | 1165 | label fix (previous: CCA1170); technical replicates |
| halophila_CCA0111 | DN 155 | *Symphyomyrtus* | *Adnataria* | 1187 | |
| hebetifolia_CCA0398 | DN 240 | *Symphyomyrtus* | *Glandulosae* | 1185 | |
| henryi_CCA1609 | DN 1259 | *Blakella* | ‘*Maculatae*’ | 1156 | technical replicates |
| henryi_CCA1610 | DN 1259 | *Blakella* | ‘*Maculatae*’ | 1160 | technical replicates |
| henryi_CCA1611 | DN 1259 | *Blakella* | ‘*Maculatae*’ | 1163 | |
| herbertiana_CCA1439 | DN 1933 | *Symphyomyrtus* | *Exsertaria* | 1178 | |
| histophylla_CCA0258 | DN 124 | *Symphyomyrtus* | *Glandulosae* | 1186 | |
| histophylla_CCA1445 | DN 124 | *Symphyomyrtus* | *Glandulosae* | 1187 | technical replicates |
| horistes_CCA1178 | DN 1169 | *Symphyomyrtus* | *Bisectae* | 1186 | label fix (previous: CCA1286) |
| horistes_CCA5984 | DN 6985 | *Symphyomyrtus* | *Bisectae* | 1185 | technical replicates |
| horistes_CCA5985 | DN 6985 | *Symphyomyrtus* | *Bisectae* | 1185 | |
| hypolaena_CCA0507 | DN 513 | *Symphyomyrtus* | *Bisectae* | 1187 | |
| incerata_CCA0281 | DN 327 | *Symphyomyrtus* | *Glandulosae* | 1186 | |
| incrassata_CCA1174 | DN 930 | *Symphyomyrtus* | *Dumaria* | 1179 | label fix (previous: CCA1291) |
| incrassata_CCA1186 | DN 1108 | *Symphyomyrtus* | *Dumaria* | 1187 | label fix (previous: CCA1275) |
| incrassata_CCA1231 | DN 1129 | *Symphyomyrtus* | *Dumaria* | 1187 | label fix (previous: CCA1250) |
| incrassata_CCA2927 | DN 1638 | *Symphyomyrtus* | *Dumaria* | 1186 | |
| indurata_CCA0121 | DN 165 | *Symphyomyrtus* | *Bisectae* | 1185 | technical replicates |
| indurata_CCA0122 | DN 165 | *Symphyomyrtus* | *Bisectae* | 1186 | |
| infera_CCA1594 | DN 1254 | *Symphyomyrtus* | *Exsertaria* | 1177 | |
| infera_CCA1595 | DN 1254 | *Symphyomyrtus* | *Exsertaria* | 1184 | putative contamination / mislabelling |

| | | | | | |
|---|---|---|---|---|---|
| insularis_continentalis_CCA5718 | DN 1637 | *Eucalyptus* | *Frutices* | 1186 | |
| intertexta_CCA0650 | DN 423 | *Symphyomyrtus* | *Adnataria* | 999 | technical replicates |
| intertexta_CCA0652 | DN 423 | *Symphyomyrtus* | *Adnataria* | 1187 | |
| intertexta_CCA0653 | DN 423 | *Symphyomyrtus* | *Adnataria* | 1186 | technical replicates |
| jacksonii_CCA5906 | DN 7223 | *Eucalyptus* | *Longistylus* | 1186 | technical replicates |
| jacksonii_CCA5907 | DN 7223 | *Eucalyptus* | *Longistylus* | 1187 | |
| jensenii_CCA2984 | DN 1916 | *Symphyomyrtus* | *Adnataria* | 1187 | |
| jucunda_CCA6041 | DN 7228 | *Eudesmia* | *Limbatae* | 1162 | technical replicates |
| jucunda_CCA6044 | DN 7228 | *Eudesmia* | *Limbatae* | 1184 | |
| jutsonii_jutsonii_CCA0537 | DN 548 | *Symphyomyrtus* | *Bisectae* | 1056 | |
| kabiana_CCA0773 | DN 720 | *Symphyomyrtus* | *Exsertaria* | 1159 | |
| kessellii_eugnosta_CCA0129 | DN 164 | *Symphyomyrtus* | *Bisectae* | 1187 | technical replicates |
| kessellii_eugnosta_CCA0130 | DN 164 | *Symphyomyrtus* | *Bisectae* | 1186 | |
| kingsmillii_CCA0520 | DN 535 | *Symphyomyrtus* | *Bisectae* | 1187 | |
| kingsmillii_CCA5803 | DN 5779 | *Symphyomyrtus* | *Bisectae* | 1185 | |
| kitsoniana_x_ovata_grandiflora_CCA2851 | DN 1720 | *Symphyomyrtus* | *Maidenaria* | 1187 | |
| kochii_amaryssia_CCA0546 | DN 536 | *Symphyomyrtus* | *Bisectae* | 1184 | |
| kochii_borealis_CCA0263 | DN 277 | *Symphyomyrtus* | *Bisectae* | 1186 | |
| kochii_kochii_CCA0278 | DN 288 | *Symphyomyrtus* | *Bisectae* | 1113 | technical replicates |
| kochii_kochii_CCA0279 | DN 288 | *Symphyomyrtus* | *Bisectae* | 1186 | |
| kondininensis_CCA0420 | DN 242 | *Symphyomyrtus* | *Dumaria* | 1186 | |
| kruseana_CCA0656 | DN 337 | *Symphyomyrtus* | *Glandulosae* | 1186 | |
| kruseana_CCA6038 | DN 6993 | *Symphyomyrtus* | *Glandulosae* | 1187 | |
| kumarlensis_CCA0081 | DN 141 | *Symphyomyrtus* | *Bisectae* | 1186 | |
| kumarlensis_CCA2861 | DN 1844 | *Symphyomyrtus* | *Bisectae* | 1186 | |

| | | | | | |
|---|---|---|---|---|---|
| laevis_CCA0059 | DN 120 | *Symphyomyrtus* | *Dumaria* | 1187 | |
| lansdowneana_CCA0819 | DN 57 | *Symphyomyrtus* | *Adnataria* | 1162 | technical replicates |
| lansdowneana_CCA1425 | DN 57 | *Symphyomyrtus* | *Adnataria* | 1186 | |
| lansdowneana_Ferguson2024 | | *Symphyomyrtus* | *Adnataria* | 1187 | not CCA |
| largiflorens_CCA0691 | | *Symphyomyrtus* | *Adnataria* | 1176 | technical replicates |
| largiflorens_CCA0692 | | *Symphyomyrtus* | *Adnataria* | 1186 | |
| largiflorens_CCA1139 | DN 980 | *Symphyomyrtus* | *Adnataria* | 1187 | label fix (previous: CCA1342); technical replicates |
| largiflorens_CCA1142 | DN 980 | *Symphyomyrtus* | *Adnataria* | 1186 | label fix (previous: CCA1339) |
| largiflorens_CCA1143 | DN 980 | *Symphyomyrtus* | *Adnataria* | 1187 | label fix (previous: CCA1336); technical replicates |
| latens_CCA2969 | DN 1613 | *Symphyomyrtus* | *Bisectae* | 1186 | |
| lehmannii_lehmannii_CCA1452 | DN 222 | *Symphyomyrtus* | *Glandulosae* | 1187 | |
| lehmannii_parallela_CCA2886 | DN 1639 | *Symphyomyrtus* | *Glandulosae* | 1187 | |
| leichhardii_CCA1548 | DN 1277 | *Blakella* | ‘*Naviculares*’ | 1156 | technical replicates |
| leichhardii_CCA1550 | DN 1277 | *Blakella* | ‘*Naviculares*’ | 1157 | |
| leiocarpa_CCA1427 | DN 2103 | *Angophora* | | 1153 | |
| leiocarpa_CCA1430 | DN 2103 | *Angophora* | | 1151 | technical replicates |
| lenziana_CCA5848 | DN 5733 | *Corymbia* | *Notiales* | 1167 | |
| leptocalyx_leptocalyx_CCA0231 | DN 153 | *Symphyomyrtus* | *Dumaria* | 1187 | |
| leptoloma_CCA1688 | DN 1302 | *Blakella* | ‘*Naviculares*’ | 1162 | putative contamination / mislabelling |
| leptophleba_CCA2845 | DN 1313 | *Symphyomyrtus* | *Adnataria* | 1185 | |
| leptophylla_CCA0636 | DN 447 | *Symphyomyrtus* | *Bisectae* | 1185 | |
| leptophylla_CCA5990 | DN 6807 | *Symphyomyrtus* | *Bisectae* | 1186 | |
| leptophylla_CCA5991 | DN 6807 | *Symphyomyrtus* | *Bisectae* | 1187 | technical replicates |
| leptopoda_elevata_CCA0002 | DN 8 | *Symphyomyrtus* | *Bisectae* | 1183 | |
| leptopoda_leptopoda_CCA0163 | DN 298 | *Symphyomyrtus* | *Bisectae* | 1186 | |

| | | | | | |
|---|---|---|---|---|---|
| leptopoda_subluta_CCA0065 | DN 336 | *Symphyomyrtus* | *Bisectae* | 1187 | |
| leptopoda_x_oldfieldii_intergrade_CCA5639 | DN 5583 | *Symphyomyrtus* | *Bisectae* | 1187 | |
| lesouefii_CCA0562 | DN 549 | *Symphyomyrtus* | *Dumaria* | 1187 | |
| leucophloia_CCA0469 | | *Symphyomyrtus* | *Platysperma* | 1184 | technical replicates |
| leucophloia_CCA0470 | | *Symphyomyrtus* | *Platysperma* | 1186 | |
| leucophloia_euroa_CCA1402 | DN 1332 | *Symphyomyrtus* | *Platysperma* | 1187 | |
| leucophloia_Ferguson2024 | | *Symphyomyrtus* | *Platysperma* | 1185 | not CCA |
| leucophloia_leucophloia_CCA0560 | DN 539 | *Symphyomyrtus* | *Platysperma* | 1186 | |
| leucophylla_CCA1500 | DN 1331 | *Symphyomyrtus* | *Adnataria* | 934 | technical replicates |
| leucophylla_CCA1501 | DN 1331 | *Symphyomyrtus* | *Adnataria* | 1163 | |
| leucophylla_CCA2877 | DN 1941 | *Symphyomyrtus* | *Adnataria* | 1186 | |
| leucophylla_CCA2940 | DN 1937 | *Symphyomyrtus* | *Adnataria* | 1185 | |
| leucoxylon_CCA0472 | | *Symphyomyrtus* | *Adnataria* | 1187 | |
| leucoxylon_leucoxylon_CCA3841 | DN 2585 | *Symphyomyrtus* | *Adnataria* | 1176 | |
| leucoxylon_megalocarpa_CCA0633 | DN 473 | *Symphyomyrtus* | *Adnataria* | 1186 | |
| leucoxylon_megalocarpa_CCA5654 | DN 4310 | *Symphyomyrtus* | *Adnataria* | 1186 | |
| leucoxylon_megalocarpa_CCA5656 | DN 4310 | *Symphyomyrtus* | *Adnataria* | 1185 | technical replicates |
| leucoxylon_petiolaris_CCA1342 | DN 945 | *Symphyomyrtus* | *Adnataria* | 1186 | label fix (previous: CCA1142) |
| leucoxylon_pruinosa_CCA0646 | DN 351 | *Symphyomyrtus* | *Adnataria* | 444 | |
| lirata_CCA5558 | DN 1917 | *Eudesmia* | *Reticulatae* | 1129 | |
| litorea_CCA0291 | DN 199 | *Symphyomyrtus* | *Dumaria* | 1187 | |
| lockyeri_CCA0461 | | *Symphyomyrtus* | *Exsertaria* | 1187 | technical replicates |
| lockyeri_CCA0464 | | *Symphyomyrtus* | *Exsertaria* | 1187 | |
| longifolia_CCA0695 | Brooker 4700 | *Symphyomyrtus* | *Incognitae* | 1146 | |
| longirostrata_CCA1417 | DN 2112 | *Symphyomyrtus* | *Pumilio* | 1187 | |

| | | | | | |
|---|---|---|---|---|---|
| longissima_CCA1539 | DN 1569 | *Symphyomyrtus* | *Bisectae* | 1184 | |
| loxophleba_gratiae_CCA0446 | DN 313 | *Symphyomyrtus* | *Glandulosae* | 1187 | |
| loxophleba_lissophloia_CCA0223 | DN 139 | *Symphyomyrtus* | *Glandulosae* | 1185 | |
| loxophleba_loxophleba_CCA0421 | DN 239 | *Symphyomyrtus* | *Glandulosae* | 1187 | |
| loxophleba_supralaevis_CCA5804 | DN 5593 | *Symphyomyrtus* | *Glandulosae* | 1187 | technical replicates |
| loxophleba_supralaevis_CCA5805 | DN 5593 | *Symphyomyrtus* | *Glandulosae* | 1186 | |
| lucasii_CCA0499 | DN 545 | *Symphyomyrtus* | *Adnataria* | 1186 | |
| lucasii_CCA0502 | DN 545 | *Symphyomyrtus* | *Adnataria* | 1186 | technical replicates |
| luculenta_CCA5987 | DN 6956 | *Symphyomyrtus* | *Bisectae* | 1185 | |
| luculenta_CCA5988 | DN 6956 | *Symphyomyrtus* | *Bisectae* | 1187 | technical replicates |
| lunata_CCA2815 | | *Symphyomyrtus* | *Bisectae* | 1187 | |
| lunata_CCA2816 | | *Symphyomyrtus* | *Bisectae* | 1186 | technical replicates |
| lunata_CCA5663 | DN 5518 | *Symphyomyrtus* | *Bisectae* | 1182 | |
| lunata_CCA5664 | DN 5518 | *Symphyomyrtus* | *Bisectae* | 1187 | technical replicates |
| lunata_CCA5746 | DN 5503 | *Symphyomyrtus* | *Bisectae* | 1187 | |
| lunata_CCA5747 | DN 5503 | *Symphyomyrtus* | *Bisectae* | 1186 | technical replicates |
| mackintii_CCA3030 | DN 1737 | *Eucalyptus* | *Eucalyptus* | 1187 | |
| macrandra_CCA0399 | DN 312 | *Symphyomyrtus* | *Glandulosae* | 1187 | |
| macrandra_CCA1218 | DN 1132 | *Symphyomyrtus* | *Glandulosae* | 1187 | label fix (previous: CCA1265) |
| macrocarpa_elachantha_CCA0207 | DN 250 | *Symphyomyrtus* | *Bisectae* | 1186 | |
| macrocarpa_macrocarpa_CCA0048 | DN 261 | *Symphyomyrtus* | *Bisectae* | 1187 | |
| macrorhyncha_macrorhyncha_CCA0684 | DN 737 | *Eucalyptus* | *Eucalyptus* | 938 | putative contamination / mislabelling |
| macrorhyncha_macrorhyncha_CCA0685 | DN 737 | *Eucalyptus* | *Eucalyptus* | 1187 | |
| macta_CCA1691 | DN 1301 | *Symphyomyrtus* | *Latoangulatae* | 131 | low coverage |
| maculata_CCA3011 | DN 1752 | *Blakella* | ‘*Maculatae*’ | 1134 | technical replicates |

| | | | | | |
|---|---|---|---|---|---|
| maculata_CCA3012 | DN 1752 | *Blakella* | ‘*Maculatae*’ | 1151 | technical replicates |
| maculata_CCA3013 | DN 1752 | *Blakella* | ‘*Maculatae*’ | 1164 | |
| maculata_Ferguson2024 | | *Blakella* | ‘*Maculatae*’ | 1161 | not CCA |
| magnificata_CCA3887 | DN 2945 | *Symphyomyrtus* | *Adnataria* | 1153 | |
| maidenii_CCA3092 | DN 1764 | *Symphyomyrtus* | *Maidenaria* | 1179 | |
| major_CCA3176 | DN 2107 | *Symphyomyrtus* | *Exsertaria* | 1030 | |
| malacoxylon_CCA1623 | DN 1368 | *Symphyomyrtus* | *Maidenaria* | 1187 | |
| mannensis_mannensis_CCA0003 | | *Symphyomyrtus* | *Bisectae* | 1176 | |
| mannensis_mannensis_CCA6066 | DN 6941 | *Symphyomyrtus* | *Bisectae* | 1179 | putative contamination / mislabelling |
| mannifera_CCA0931 | DN 581 | *Symphyomyrtus* | *Maidenaria* | 1178 | |
| marginata | | *Eucalyptus* | *Longistylus* | 1185 | |
| marginata_Ferguson2024 | | *Eucalyptus* | *Longistylus* | 1187 | not CCA |
| marginata_marginata_CCA5976 | DN 4655 | *Eucalyptus* | *Longistylus* | 1175 | |
| marginata_spurgeana_CCA5931 | DN 7220 | *Eucalyptus* | *Longistylus* | 1186 | technical replicates |
| marginata_spurgeana_CCA5932 | DN 7220 | *Eucalyptus* | *Longistylus* | 1186 | |
| megacarpa_CCA1242 | DN 1137 | *Eucalyptus* | *Frutices* | 1186 | label fix (previous: CCA1210) |
| megacarpa_CCA5870 | DN 6705 | *Eucalyptus* | *Frutices* | 1186 | |
| megacornuta_CCA1274 | DN 1121 | *Symphyomyrtus* | *Glandulosae* | 1187 | label fix (previous: CCA1189) |
| megacornuta_CCA1275 | DN 1121 | *Symphyomyrtus* | *Glandulosae* | 1186 | label fix (previous: CCA1186); technical replicates |
| melana_CCA3464 | DN 2552 | *Angophora* | | 1150 | |
| melanophitrus_CCA0170 | DN 211 | *Symphyomyrtus* | *Glandulosae* | 1185 | |
| melanophloia | | *Symphyomyrtus* | *Adnataria* | 1185 | invalid CCA identifier |
| melanophloia_melanophloia_CCA0743 | DN 690 | *Symphyomyrtus* | *Adnataria* | 1177 | |
| melanophloia_melanophloia_CCA1638 | DN 1285 | *Symphyomyrtus* | *Adnataria* | 1186 | |
| melanophloia_nana_CCA1496 | DN 1330 | *Symphyomyrtus* | *Adnataria* | 1186 | |

| | | | | | |
|---|---|---|---|---|---|
| melanoxylon_CCA0073 | DN 143 | *Symphyomyrtus* | *Dumaria* | 1185 | |
| melliodora_1_CCA0704 | DN 777 | *Symphyomyrtus* | *Adnataria* | 1172 | technical replicates |
| melliodora_1_Csiro | | *Symphyomyrtus* | *Adnataria* | 1182 | not CCA |
| melliodora_2_CCA0704 | DN 777 | *Symphyomyrtus* | *Adnataria* | 1169 | |
| melliodora_2_Csiro | | *Symphyomyrtus* | *Adnataria* | 1183 | not CCA |
| melliodora_CCA0705 | DN 777 | *Symphyomyrtus* | *Adnataria* | 1176 | technical replicates |
| melliodora_CCA0706 | DN 777 | *Symphyomyrtus* | *Adnataria* | 1170 | technical replicates |
| melliodora_Ferguson2024 | | *Symphyomyrtus* | *Adnataria* | 1187 | not CCA |
| melliodora_x_sideroxylon_CCA1504 | DN 1219 | *Symphyomyrtus* | *Adnataria* | 1180 | technical replicates |
| melliodora_x_sideroxylon_CCA1505 | DN 1219 | *Symphyomyrtus* | *Adnataria* | 1175 | technical replicates |
| melliodora_x_sideroxylon_CCA1506 | DN 1219 | *Symphyomyrtus* | *Adnataria* | 817 | technical replicates |
| melliodora_x_sideroxylon_CCA1507 | DN 1219 | *Symphyomyrtus* | *Adnataria* | 1170 | technical replicates |
| melliodora_x_sideroxylon_CCA6869 | DN 1219 | *Symphyomyrtus* | *Adnataria* | 1171 | technical replicates |
| melliodora_x_sideroxylon_CCA6870 | DN 1219 | *Symphyomyrtus* | *Adnataria* | 1172 | |
| melliodora_x_sideroxylon_CCA6871 | DN 1219 | *Symphyomyrtus* | *Adnataria* | 1138 | technical replicates |
| merrickiae_CCA0208 | DN 156 | *Symphyomyrtus* | *Dumaria* | 1187 | |
| michaeliana_CCA0964 | DN 650 | *Symphyomyrtus* | *Racemus* | 927 | technical replicates |
| michaeliana_CCA0965 | DN 650 | *Symphyomyrtus* | *Racemus* | 1051 | |
| micranthera_CCA0248 | DN 172 | *Symphyomyrtus* | *Bisectae* | 1168 | |
| microcarpa_CCA1511 | DN 1364 | *Symphyomyrtus* | *Adnataria* | 213 | low coverage |
| microcarpa_CCA1543 | DN 1220 | *Symphyomyrtus* | *Adnataria* | 1154 | technical replicates |
| microcarpa_CCA1545 | DN 1220 | *Symphyomyrtus* | *Adnataria* | 1187 | |
| microcodon_CCA5568 | DN 5010 | *Eucalyptus* | *Eucalyptus* | 1151 | |
| microcorys_CCA1660 | DN 1238 | *Alveolata* | | 1061 | |
| microcorys_Ferguson2024 | | *Alveolata* | | 1186 | not CCA |

| | | | | | |
|---|---|---|---|---|---|
| microtheca_CCA1518 | DN 1338 | *Symphyomyrtus* | *Adnataria* | 1000 | |
| mimica_mimica_CCA1461 | DN 188 | *Symphyomyrtus* | *Glandulosae* | 1187 | |
| minniritchi_CCA2839 | DN 1354 | *Symphyomyrtus* | *Bisectae* | 1186 | |
| misella_CCA0374 | DN 147 | *Symphyomyrtus* | *Bisectae* | 1187 | |
| moderata_CCA2887 | DN 1661 | *Symphyomyrtus* | *Bisectae* | 1187 | |
| moderata_CCA2890 | DN 1661 | *Symphyomyrtus* | *Bisectae* | 1184 | technical replicates |
| moluccana_CCA0751 | DN 712 | *Symphyomyrtus* | *Adnataria* | 1158 | |
| molyneuxii_CCA5940 | DN 6867 | *Eucalyptus* | *Eucalyptus* | 1186 | |
| moorei_serpentinicola_CCA1616 | DN 1233 | *Eucalyptus* | *Eucalyptus* | 1187 | |
| morrisbyi_CCA5915 | DN 1983 | *Symphyomyrtus* | *Maidenaria* | 1187 | |
| morrisbyi_CCA5917 | DN 1983 | *Symphyomyrtus* | *Maidenaria* | 1186 | technical replicates |
| muelleriana_CCA3062 | DN 1753 | *Eucalyptus* | *Eucalyptus* | 1174 | |
| myriadena_CCA0034 | DN 316 | *Symphyomyrtus* | *Dumaria* | 1187 | |
| myriadena_CCA0035 | DN 316 | *Symphyomyrtus* | *Dumaria* | 1186 | technical replicates |
| myriadena_CCA0272 | DN 321 | *Symphyomyrtus* | *Dumaria* | 1187 | technical replicates |
| myriadena_CCA0273 | DN 321 | *Symphyomyrtus* | *Dumaria* | 1186 | |
| myriadena_CCA5685 | DN 5596 | *Symphyomyrtus* | *Dumaria* | 1186 | |
| myriadena_CCA5687 | DN 5596 | *Symphyomyrtus* | *Dumaria* | 1187 | technical replicates |
| nebulosa_CCA5911 | DN 6742 | *Eucalyptus* | *Eucalyptus* | 1187 | |
| nebulosa_CCA5912 | DN 6742 | *Eucalyptus* | *Eucalyptus* | 1187 | technical replicates |
| neutra_CCA0156 | DN 218 | *Symphyomyrtus* | *Bisectae* | 1186 | |
| neutra_CCA0425 | DN 315 | *Symphyomyrtus* | *Bisectae* | 1186 | |
| neutra_CCA2879 | DN 1874 | *Symphyomyrtus* | *Bisectae* | 1186 | |
| neutra_CCA2935 | DN 1855 | *Symphyomyrtus* | *Bisectae* | 1186 | |
| neutra_CCA2987 | DN 1853 | *Symphyomyrtus* | *Bisectae* | 1186 | technical replicates |

| | | | | | |
|---|---|---|---|---|---|
| neutra_CCA2988 | DN 1853 | *Symphyomyrtus* | *Bisectae* | 1187 | |
| nicholii_CCA0754 | DN 651 | *Symphyomyrtus* | *Maidenaria* | 1184 | |
| nicholii_CCA0754_2 | DN 651 | *Symphyomyrtus* | *Maidenaria* | 1184 | technical replicates |
| nitens_CCA3107 | DN 1766 | *Symphyomyrtus* | *Maidenaria* | 973 | |
| nobilis_CCA0920 | DN 753 | *Symphyomyrtus* | *Maidenaria* | 1176 | |
| normantonensis_CCA1130 | DN 1207 | *Symphyomyrtus* | *Adnataria* | 1185 | label fix (previous: CCA1329) |
| notabilis_CCA0831 | DN 630 | *Symphyomyrtus* | *Latoangulatae* | 1174 | putative contamination / mislabelling |
| notactites_CCA1447 | DN 177 | *Symphyomyrtus* | *Bisectae* | 1187 | |
| obesa_CCA1192 | DN 1109 | *Symphyomyrtus* | *Bisectae* | 978 | technical replicates |
| obesa_CCA1193 | DN 1109 | *Symphyomyrtus* | *Bisectae* | 1187 | label fix (previous: CCA1271) |
| obesa_CCA4557 | DN 3724 | *Symphyomyrtus* | *Bisectae* | 1026 | |
| obesa_CCA5705 | DN 5478 | *Symphyomyrtus* | *Bisectae* | 1185 | |
| obliqua_CCA1300 | DN 804 | *Eucalyptus* | *Eucalyptus* | 1182 | |
| obtusiflora_dongarraensis_CCA0045 | DN 266 | *Symphyomyrtus* | *Dumaria* | 1187 | |
| occidentalis_CCA1278 | DN 170 | *Symphyomyrtus* | *Glandulosae* | 1186 | label fix (previous: CCA1184) |
| ochrophloia_CCA1419 | DN 2119 | *Symphyomyrtus* | *Adnataria* | 1186 | |
| odontocarpa_CCA3651 | DN 1199 | *Eudesmia* | *Limbatae* | 1168 | |
| odorata_cajuputea_CCA2917 | DN 1691 | *Symphyomyrtus* | *Adnataria* | 1170 | technical replicates |
| odorata_cajuputea_CCA2918 | DN 1691 | *Symphyomyrtus* | *Adnataria* | 1187 | |
| odorata_odorata_CCA1085 | DN 936 | *Symphyomyrtus* | *Adnataria* | 1133 | |
| odorata_odorata_CCA1147 | DN 822 | *Symphyomyrtus* | *Adnataria* | 1187 | label fix (previous: CCA1323) |
| odorata_polybractea_CCA0465 | | *Symphyomyrtus* | *Adnataria* | 1166 | putative contamination / mislabelling |
| odorata_polybractea_CCA0466 | | *Symphyomyrtus* | *Adnataria* | 1008 | technical replicates |
| odorata_polybractea_CCA0467 | | *Symphyomyrtus* | *Adnataria* | 1185 | technical replicates |
| odorata_polybractea_CCA0468 | | *Symphyomyrtus* | *Adnataria* | 1186 | |

| | | | | | |
|---|---|---|---|---|---|
| odorata_wimmerensis_CCA1523 | DN 1590 | *Symphyomyrtus* | *Adnataria* | 1187 | |
| odorata_wimmerensis_CCA5956 | DN 6866 | *Symphyomyrtus* | *Adnataria* | 1187 | |
| odorata_wimmerensis_CCA5957 | DN 6866 | *Symphyomyrtus* | *Adnataria* | 1186 | technical replicates |
| odorata_x_phenax_CCA1520 | DN 1360 | *Symphyomyrtus* | | 1186 | inter-sectional hybrid |
| oleosa_ampliata_CCA0505 | DN 500 | *Symphyomyrtus* | *Bisectae* | 1185 | |
| oleosa_ampliata_CCA0623 | DN 555 | *Symphyomyrtus* | *Bisectae* | 1185 | |
| oleosa_ampliata_CCA1498 | DN 1408 | *Symphyomyrtus* | *Bisectae* | 1021 | |
| oleosa_cylindroidea_CCA2930 | DN 1642 | *Symphyomyrtus* | *Bisectae* | 1187 | |
| oligantha_CCA2971 | DN 1919 | *Symphyomyrtus* | *Adnataria* | 1187 | |
| oligantha_CCA5658 | DN 5810 | *Symphyomyrtus* | *Adnataria* | 1185 | |
| oligantha_CCA5660 | DN 5810 | *Symphyomyrtus* | *Adnataria* | 1185 | technical replicates |
| oligantha_CCA5681 | DN 5804 | *Symphyomyrtus* | *Adnataria* | 1184 | technical replicates |
| oligantha_CCA5683 | DN 5804 | *Symphyomyrtus* | *Adnataria* | 1186 | |
| olivina_CCA0289 | DN 319 | *Symphyomyrtus* | *Bisectae* | 1184 | |
| olivina_CCA0290 | DN 319 | *Symphyomyrtus* | *Bisectae* | 1185 | technical replicates |
| omissa_CCA0541 | DN 505 | *Symphyomyrtus* | *Bisectae* | 1186 | |
| omissa_CCA1485 | DN 1392 | *Symphyomyrtus* | *Bisectae* | 1182 | |
| ophitica_1_CCA3191 | DN 2099 | *Symphyomyrtus* | *Adnataria* | 976 | technical replicates |
| ophitica_2_CCA3191 | DN 2099 | *Symphyomyrtus* | *Adnataria* | 986 | |
| opimiflora_CCA5698 | DN 5547 | *Symphyomyrtus* | *Bisectae* | 1186 | |
| optima_CCA0431 | DN 115 | *Symphyomyrtus* | *Bisectae* | 1177 | technical replicates |
| optima_CCA0432 | DN 115 | *Symphyomyrtus* | *Bisectae* | 1186 | |
| oraria_CCA0041 | DN 269 | *Symphyomyrtus* | *Dumaria* | 1187 | technical replicates |
| oraria_CCA0043 | DN 269 | *Symphyomyrtus* | *Dumaria* | 1187 | |
| orbifolia_CCA5843 | DN 5606 | *Symphyomyrtus* | *Bisectae* | 1186 | |

| | | | | | |
|---|---|---|---|---|---|
| ornata_CCA0455 | DN 243 | *Symphyomyrtus* | *Bisectae* | 1114 | |
| ornata_CCA0456 | DN 243 | *Symphyomyrtus* | *Bisectae* | 1186 | technical replicates |
| ovata_grandiflora_CCA1627 | DN 1594 | *Symphyomyrtus* | *Maidenaria* | 1187 | |
| ovata_ovata_CCA1377 | DN 805 | *Symphyomyrtus* | *Maidenaria* | 1164 | technical replicates |
| ovata_ovata_CCA1379 | DN 805 | *Symphyomyrtus* | *Maidenaria* | 1187 | |
| ovata_ovata_CCA3005 | DN 1741 | *Symphyomyrtus* | *Maidenaria* | 1179 | |
| ovata_ovata_CCA3006 | DN 1741 | *Symphyomyrtus* | *Maidenaria* | 1187 | technical replicates |
| ovata_ovata_CCA5868 | DN 6741 | *Symphyomyrtus* | *Maidenaria* | 1186 | |
| ovata_ovata_CCA5892 | DN 6725 | *Symphyomyrtus* | *Maidenaria* | 1187 | |
| ovularis_CCA1196 | DN 1107 | *Symphyomyrtus* | *Dumaria* | 1181 | label fix (previous: CCA1270) |
| oxymitra_CCA0015 | | *Symphyomyrtus* | *Bisectae* | 1186 | |
| oxymitra_CCA2904 | DN 1704 | *Symphyomyrtus* | *Bisectae* | 1187 | |
| pachycalyx_pachycalyx_CCA3025 | DN 1307 | *Symphyomyrtus* | *Bisectae* | 1184 | |
| pallida_CCA5774 | DN 5740 | *Eudesmia* | *Limbatae* | 1182 | |
| pallida_CCA6083 | DN 7229 | *Eudesmia* | *Limbatae* | 1181 | |
| panda_CCA0848 | DN 683 | *Symphyomyrtus* | *Adnataria* | 1164 | |
| panda_CCA1682 | DN 1275 | *Symphyomyrtus* | *Adnataria* | 1184 | |
| paniculata_Ferguson2024 | | *Symphyomyrtus* | *Adnataria* | 1186 | not CCA |
| papillosa_CCA5646 | DN 5806 | *Corymbia* | *Notiales* | 1162 | |
| patellaris_CCA1552 | DN 1339 | *Symphyomyrtus* | *Adnataria* | 1187 | |
| patens_CCA4960 | DN 4621 | *Eucalyptus* | *Longistylus* | 1171 | |
| patens_CCA5641 | DN 5530 | *Eucalyptus* | *Longistylus* | 1187 | |
| pauciflora_parvifructa_CCA5946 | DN 7047 | *Eucalyptus* | *Eucalyptus* | 1038 | technical replicates |
| pauciflora_parvifructa_CCA5948 | DN 7047 | *Eucalyptus* | *Eucalyptus* | 1186 | technical replicates |
| pauciflora_parvifructa_CCA5949 | DN 7047 | *Eucalyptus* | *Eucalyptus* | 1186 | |

| | | | | | |
|---|---|---|---|---|---|
| pauciflora_pauciflora_CCA0933 | DN 476 | *Eucalyptus* | *Eucalyptus* | 1183 | |
| peninsularis_CCA1322 | DN 948 | *Symphyomyrtus* | *Bisectae* | 1183 | label fix (previous: CCA1148) |
| peninsularis_CCA2983 | DN 1955 | *Symphyomyrtus* | *Bisectae* | 1185 | |
| peninsularis_CCA3022 | DN 1953 | *Symphyomyrtus* | *Bisectae* | 1186 | |
| perangusta_CCA0103 | DN 148 | *Symphyomyrtus* | *Bisectae* | 1186 | |
| percostata_CCA2914 | DN 1603 | *Symphyomyrtus* | *Dumaria* | 1187 | technical replicates |
| percostata_CCA2916 | DN 1603 | *Symphyomyrtus* | *Dumaria* | 1186 | |
| perriniana_perriniana_CCA6006 | DN 1975 | *Symphyomyrtus* | *Maidenaria* | 1187 | |
| perriniana_perriniana_CCA6008 | DN 1975 | *Symphyomyrtus* | *Maidenaria* | 1186 | technical replicates |
| pertenuis_CCA0191 | DN 326 | *Symphyomyrtus* | *Dumaria* | 1186 | |
| petalophylla_CCA1569 | DN 1262 | *Blakella* | '*Naviculares*' | 1162 | |
| petraea_CCA0069 | DN 339 | *Symphyomyrtus* | *Adnataria* | 1187 | |
| petraea_CCA5757 | DN 5463 | *Symphyomyrtus* | *Adnataria* | 1187 | technical replicates |
| petraea_CCA5758 | DN 5463 | *Symphyomyrtus* | *Adnataria* | 1187 | |
| petrensis_CCA0451 | DN 248 | *Symphyomyrtus* | *Bisectae* | 1187 | |
| phaenophylla_CCA0295 | DN 324 | *Symphyomyrtus* | *Glandulosae* | 1185 | |
| phaenophylla_CCA0304 | DN 196 | *Symphyomyrtus* | *Glandulosae* | 1187 | |
| phaenophylla_CCA0308 | DN 236 | *Symphyomyrtus* | *Glandulosae* | 1185 | |
| phenax_phenax_CCA1286 | DN 929 | *Symphyomyrtus* | *Dumaria* | 1185 | label fix (previous: CCA1178) |
| phenax_phenax_CCA1288 | DN 929 | *Symphyomyrtus* | *Dumaria* | 1185 | label fix (previous: CCA1177); technical replicates |
| pileata_CCA0227 | DN 191 | *Symphyomyrtus* | *Dumaria* | 1187 | |
| pileata_CCA1197 | DN 947 | *Symphyomyrtus* | *Dumaria* | 1185 | label fix (previous: CCA1269);<br>putative contamination / mislabelling |
| pileata_CCA5845 | DN 5477 | *Symphyomyrtus* | *Dumaria* | 1186 | |
| pileata_CCA5847 | DN 5477 | *Symphyomyrtus* | *Dumaria* | 1187 | technical replicates |

| | | | | | |
|---|---|---|---|---|---|
| pilularis_CCA3322 | DN 644 | *Eucalyptus* | *Eucalyptus* | 1163 | |
| pimpiniana_CCA0021 | DN 110 | *Symphyomyrtus* | *Dumaria* | 1186 | |
| planipes_CCA0149 | DN 330 | *Symphyomyrtus* | *Dumaria* | 1187 | technical replicates |
| planipes_CCA0150 | DN 330 | *Symphyomyrtus* | *Dumaria* | 1187 | |
| planipes_CCA0330 | DN 131 | *Symphyomyrtus* | *Dumaria* | 1187 | |
| planipes_CCA6011 | DN 6996 | *Symphyomyrtus* | *Dumaria* | 1187 | |
| planipes_CCA6013 | DN 6996 | *Symphyomyrtus* | *Dumaria* | 1187 | technical replicates |
| platydisca_CCA0051 | DN 133 | *Eucalyptus* | *Frutices* | 1187 | |
| platyphylla_CCA1564 | DN 1350 | *Symphyomyrtus* | *Exsertaria* | 1167 | |
| platyphylla_CCA1661 | DN 1296 | *Symphyomyrtus* | *Exsertaria* | 1186 | |
| platypus_platypus_CCA0602 | DN 208 | *Symphyomyrtus* | *Glandulosae* | 1186 | technical replicates |
| platypus_platypus_CCA0603 | DN 208 | *Symphyomyrtus* | *Glandulosae* | 1186 | |
| pleurocarpa_CCA1256 | DN 154 | *Eudesmia* | *Limbatae* | 1161 | putative contamination / mislabelling |
| pleurocarpa_CCA1257 | DN 154 | *Eudesmia* | *Limbatae* | 1183 | label fix (previous: CCA1224) |
| plumula_CCA0521 | DN 522 | *Symphyomyrtus* | *Bisectae* | 1186 | |
| plumula_CCA0524 | DN 522 | *Symphyomyrtus* | *Bisectae* | 1143 | technical replicates |
| plumula_CCA6014 | DN 6931 | *Symphyomyrtus* | *Bisectae* | 1187 | |
| plumula_CCA6017 | DN 6931 | *Symphyomyrtus* | *Bisectae* | 1187 | technical replicates |
| polyanthemos_Ferguson2024 | | *Symphyomyrtus* | *Adnataria* | 1184 | not CCA |
| polyanthemos_polyanthemos_CCA0672 | DN 742 | *Symphyomyrtus* | *Adnataria* | 1172 | |
| polyanthemos_vestita_CCA0724 | DN 457 | *Symphyomyrtus* | *Adnataria* | 1169 | |
| polycarpa_CCA5901 | DN 6961 | *Corymbia* | *Notiales* | 1167 | |
| polycarpa_CCA5902 | DN 6961 | *Corymbia* | *Notiales* | 1167 | technical replicates |
| populnea_CCA0006 | | *Symphyomyrtus* | *Adnataria* | 1185 | |
| populnea_CCA0941 | DN 679 | *Symphyomyrtus* | *Adnataria* | 1167 | |

| | | | | | |
|---|---|---|---|---|---|
| porosa_CCA0525 | DN 482 | *Symphyomyrtus* | *Adnataria* | 769 | technical replicates |
| porosa_CCA0528 | DN 482 | *Symphyomyrtus* | *Adnataria* | 1185 | |
| praetermissa_CCA0483 | DN 213 | *Symphyomyrtus* | *Glandulosae* | 1185 | |
| prava_CCA0800 | DN 671 | *Symphyomyrtus* | *Exsertaria* | 1172 | |
| preissiana_lobata_CCA0185 | DN 179 | *Eucalyptus* | *Frutices* | 1187 | |
| preissiana_lobata_CCA5695 | DN 5492 | *Eucalyptus* | *Frutices* | 1187 | technical replicates |
| preissiana_lobata_CCA5696 | DN 5492 | *Eucalyptus* | *Frutices* | 1187 | |
| prominens_CCA2840 | DN 1184 | *Symphyomyrtus* | *Glandulosae* | 1186 | |
| propinqua_CCA1692 | DN 1239 | *Symphyomyrtus* | *Exsertaria* | 1184 | |
| protensa_CCA0120 | DN 126 | *Symphyomyrtus* | *Glandulosae* | 1187 | |
| pruiniramis_CCA0268 | DN 284 | *Symphyomyrtus* | *Glandulosae* | 1187 | |
| pruinosa_pruinosa_CCA1553 | DN 1329 | *Symphyomyrtus* | *Adnataria* | 1186 | |
| pterocarpa_CCA0614 | DN 137 | *Symphyomyrtus* | *Dumaria* | 447 | |
| ptychocarpa_ptychocarpa_CCA5905 | DN 1912 | *Corymbia* | *Notiales* | 1166 | |
| pulverulenta_CCA1662 | DN 1226 | *Symphyomyrtus* | *Maidenaria* | 1187 | |
| pumila_Ferguson2024 | | *Symphyomyrtus* | *Pumilio* | 1185 | not CCA |
| punctata_CCA1325 | DN 612 | *Symphyomyrtus* | *Pumilio* | 1175 | label fix (previous: CCA1133) |
| pyriformis_CCA2862 | DN 1660 | *Symphyomyrtus* | *Bisectae* | 1187 | |
| pyrocarpa_CCA5610 | DN 2093 | *Eucalyptus* | *Eucalyptus* | 1187 | |
| quadrangulata_CCA0874 | DN 618 | *Symphyomyrtus* | *Maidenaria* | 1155 | |
| quadrans_CCA1318 | DN 1099 | *Symphyomyrtus* | *Adnataria* | 1186 | label fix (previous: CCA1153) |
| quaerenda_CCA0274 | DN 237 | *Symphyomyrtus* | *Bisectae* | 1187 | |
| quaerenda_CCA0277 | DN 237 | *Symphyomyrtus* | *Bisectae* | 1187 | technical replicates |
| radiata_radiata_CCA3899 | DN 602 | *Eucalyptus* | *Eucalyptus* | 1181 | |
| rameliana_CCA0571 | DN 543 | *Symphyomyrtus* | *Bisectae* | 1186 | label fix (previous: CCA0577) |

| | | | | | |
|---|---|---|---|---|---|
| rameliana_CCA1412 | DN 543 | *Symphyomyrtus* | *Bisectae* | 1186 | |
| ravensthorpensis_CCA2854 | DN 1636 | *Symphyomyrtus* | *Glandulosae* | 1186 | |
| raveretiana_CCA1669 | DN 1297 | *Symphyomyrtus* | *Domesticae* | 1185 | |
| ravida_CCA0123 | DN 134 | *Symphyomyrtus* | *Glandulosae* | 1186 | |
| recta_CCA0390 | DN 294 | *Symphyomyrtus* | *Bisectae* | 1174 | technical replicates |
| recta_CCA0391 | DN 294 | *Symphyomyrtus* | *Bisectae* | 1187 | |
| recta_CCA1450 | DN 294 | *Symphyomyrtus* | *Bisectae* | 1187 | technical replicates |
| redunca_pluricaulis_CCA2990 | DN 1662 | *Symphyomyrtus* | *Glandulosae* | 1187 | |
| redunca_porphyrea_CCA1260 | DN 210 | *Symphyomyrtus* | *Glandulosae* | 1186 | label fix (previous: CCA1220) |
| redunca_redunca_CCA0139 | DN 221 | *Symphyomyrtus* | *Glandulosae* | 1187 | |
| regnans_CCA5597 | DN 4316 | *Eucalyptus* | *Eucalyptus* | 1167 | |
| regnans_Ferguson2024 | | *Eucalyptus* | *Eucalyptus* | 1187 | not CCA |
| relicta_CCA5717 | DN 5535 | *Symphyomyrtus* | *Bisectae* | 1187 | |
| repullulans_CCA1490 | DN 1550 | *Symphyomyrtus* | *Dumaria* | 998 | |
| revelata_CCA6088 | DN 1910 | *Symphyomyrtus* | *Exsertaria* | 1187 | |
| rhodops_CCA1674 | DN 1306 | *Corymbia* | *Notiales* | 1166 | |
| rigens_CCA0353 | DN 157 | *Symphyomyrtus* | *Dumaria* | 1187 | |
| rigidula_rigidula_CCA0189 | DN 292 | *Symphyomyrtus* | *Bisectae* | 1186 | technical replicates |
| rigidula_rigidula_CCA0190 | DN 292 | *Symphyomyrtus* | *Bisectae* | 1186 | |
| rigidula_rigidula_CCA5753 | DN 5671 | *Symphyomyrtus* | *Bisectae* | 1187 | technical replicates |
| rigidula_rigidula_CCA5754 | DN 5671 | *Symphyomyrtus* | *Bisectae* | 1184 | |
| rigidula_rigidula_CCA5770 | DN 5071 | *Symphyomyrtus* | *Bisectae* | 1185 | technical replicates |
| rigidula_rigidula_CCA5771 | DN 5071 | *Symphyomyrtus* | *Bisectae* | 1187 | |
| rigidula_rigidula_CCA5781 | DN 5554 | *Symphyomyrtus* | *Bisectae* | 1185 | |
| rigidula_rigidula_CCA5782 | DN 5554 | *Symphyomyrtus* | *Bisectae* | 80 | low coverage |

| | | | | | |
|---|---|---|---|---|---|
| rigidula_rigidula_CCA5813 | DN 5460 | *Symphyomyrtus* | *Bisectae* | 1187 | |
| rigidula_rigidula_CCA5815 | DN 5460 | *Symphyomyrtus* | *Bisectae* | 1187 | technical replicates |
| rigidula_interior_CCA6003 | DN 6955 | *Symphyomyrtus* | *Bisectae* | 1185 | technical replicates |
| rigidula_interior_CCA6005 | DN 6955 | *Symphyomyrtus* | *Bisectae* | 1185 | |
| rigidula_rigidula_CCA5856 | DN 5578 | *Symphyomyrtus* | *Bisectae* | 1187 | |
| rigidula_rigidula_CCA5857 | DN 5578 | *Symphyomyrtus* | *Bisectae* | 1184 | technical replicates |
| robusta_x_tereticornis_CCA1641 | | *Symphyomyrtus* | | 1187 | inter-sectional hybrid |
| rosacea_CCA1401 | DN 518 | *Symphyomyrtus* | *Bisectae* | 1186 | |
| rosacea_CCA6074 | DN 6939 | *Symphyomyrtus* | *Bisectae* | 1186 | |
| rossii_1_CCA0913 | DN 781 | *Eucalyptus* | *Eucalyptus* | 1155 | |
| rossii_2_CCA0913 | DN 781 | *Eucalyptus* | *Eucalyptus* | 942 | technical replicates |
| rossii_CCA0951 | DN 725 | *Eucalyptus* | *Eucalyptus* | 1183 | |
| roycei_CCA1037 | DN 1175 | *Eudesmia* | *Limbatae* | 826 | |
| rubiginosa_CCA1431 | DN 2114 | *Eucalyptus* | *Amentum* | 1186 | putative contamination / mislabelling |
| rudis_cratyantha_CCA1163 | DN 1146 | *Symphyomyrtus* | *Exsertaria* | 1187 | label fix (previous: CCA1301) |
| rudis_cratyantha_CCA5628 | DN 5538 | *Symphyomyrtus* | *Exsertaria* | 1187 | |
| rudis_cratyantha_CCA5629 | DN 5538 | *Symphyomyrtus* | *Exsertaria* | 1187 | technical replicates |
| rudis_rudis_CCA1212 | DN 1131 | *Symphyomyrtus* | *Exsertaria* | 1180 | technical replicates |
| rudis_rudis_CCA1214 | DN 1131 | *Symphyomyrtus* | *Exsertaria* | 1187 | label fix (previous: CCA1240) |
| rugosa_CCA1254 | DN 806 | *Symphyomyrtus* | *Dumaria* | 1186 | label fix (previous: CCA1229) |
| rummeryi_CCA1675 | DN 1245 | *Symphyomyrtus* | *Adnataria* | 1187 | |
| rupestris_CCA1433 | DN 1920 | *Symphyomyrtus* | *Platysperma* | 1186 | |
| rupestris_CCA5713 | DN 5813 | *Symphyomyrtus* | *Platysperma* | 1187 | |
| sabulosa_x_ovata_ovata_CCA1637 | DN 359 | *Symphyomyrtus* | *Maidenaria* | 1186 | |
| saligna_CCA3916 | DN 2924 | *Symphyomyrtus* | *Latoangulatae* | 1171 | |

| | | | | | |
|---|---|---|---|---|---|
| salmonophloia_CCA0575 | DN 341 | *Symphyomyrtus* | *Bisectae* | 1159 | technical replicates |
| salmonophloia_CCA0576 | DN 341 | *Symphyomyrtus* | *Bisectae* | 1180 | label fix (previous: CCA0593) |
| salubris_CCA0196 | DN 342 | *Symphyomyrtus* | *Glandulosae* | 1187 | |
| sargentii_sargentii_CCA5731 | DN 5551 | *Symphyomyrtus* | *Glandulosae* | 1187 | |
| sargentii_sargentii_CCA5732 | DN 5551 | *Symphyomyrtus* | *Glandulosae* | 1187 | technical replicates |
| sargentii_sargentii_CCA5733 | DN 5551 | *Symphyomyrtus* | *Glandulosae* | 1186 | technical replicates |
| scabrida_CCA1554 | DN 1278 | *Blakella* | '*Naviculares*' | 1158 | |
| scoparia_CCA1297 | DN 672 | *Symphyomyrtus* | *Maidenaria* | 1173 | technical replicates |
| scoparia_CCA1298 | DN 672 | *Symphyomyrtus* | *Maidenaria* | 1186 | label fix (previous: CCA1165) |
| scyphata_CCA5724 | DN 5508 | *Symphyomyrtus* | *Dumaria* | 1187 | |
| scyphata_CCA5725 | DN 5508 | *Symphyomyrtus* | *Dumaria* | 1186 | technical replicates |
| scyphocalyx_CCA0556 | DN 558 | *Symphyomyrtus* | *Dumaria* | 1126 | technical replicates |
| scyphocalyx_CCA0559 | DN 558 | *Symphyomyrtus* | *Dumaria* | 1185 | |
| sepulcralis_CCA2883 | DN 1635 | *Eucalyptus* | *Frutices* | 1186 | |
| serraensis_serraensis_CCA0661 | DN 455 | *Eucalyptus* | *Eucalyptus* | 1187 | |
| sessilis_CCA1521 | DN 1352 | *Symphyomyrtus* | *Bisectae* | 1183 | |
| setosa_pedicellaris_x_brachycarpa_CCA1542 | DN 1286 | *Corymbia* | *Notiales* | 1162 | |
| setosa_setosa_CCA6050 | DN 2441 | *Corymbia* | *Notiales* | 1162 | |
| sheathiana_CCA0401 | DN 302 | *Symphyomyrtus* | *Dumaria* | 1186 | technical replicates |
| sheathiana_CCA0402 | DN 302 | *Symphyomyrtus* | *Dumaria* | 1187 | |
| sheathiana_CCA5742 | DN 5453 | *Symphyomyrtus* | *Dumaria* | 1186 | |
| sheathiana_CCA5743 | DN 5453 | *Symphyomyrtus* | *Dumaria* | 1187 | technical replicates |
| shirleyi_CCA1572 | DN 1292 | *Symphyomyrtus* | *Adnataria* | 1184 | |
| shirleyi_Ferguson2024 | | *Symphyomyrtus* | *Adnataria* | 1187 | not CCA |
| sicilifolia_CCA1558 | DN 1276 | *Symphyomyrtus* | *Adnataria* | 1187 | |

| | | | | | |
|---|---|---|---|---|---|
| sideroxylon | | *Symphyomyrtus* | *Adnataria* | 1187 | invalid CCA identifier |
| sideroxylon_sideroxylon_CCA0807 | DN 674 | *Symphyomyrtus* | *Adnataria* | 1183 | |
| sideroxylon_sideroxylon_CCA0808 | DN 674 | *Symphyomyrtus* | *Adnataria* | 1178 | technical replicates |
| sideroxylon_Ferguson2024 | | *Symphyomyrtus* | *Adnataria* | 1186 | not CCA |
| sideroxylon_x_melliodora_Ferguson2024 | | *Symphyomyrtus* | *Adnataria* | 1140 | not CCA |
| sieberi_CCA3035 | DN 1749 | *Eucalyptus* | *Eucalyptus* | 1185 | technical replicates |
| sieberi_CCA3036 | DN 1749 | *Eucalyptus* | *Eucalyptus* | 1187 | |
| sinuensis_CCA1173 | DN 1081 | *Symphyomyrtus* | *Bisectae* | 1179 | label fix (previous: CCA1294) |
| socialis_socialis_CCA0166 | | *Symphyomyrtus* | *Bisectae* | 1186 | |
| socialis_socialis_CCA0167 | | *Symphyomyrtus* | *Bisectae* | 1187 | technical replicates |
| socialis_socialis_CCA2898 | DN 1804 | *Symphyomyrtus* | *Bisectae* | 1187 | |
| socialis_victoriensis_CCA0016 | DN 113 | *Symphyomyrtus* | *Bisectae* | 1177 | technical replicates |
| socialis_victoriensis_CCA0017 | DN 113 | *Symphyomyrtus* | *Bisectae* | 1187 | technical replicates |
| socialis_victoriensis_CCA0018 | DN 113 | *Symphyomyrtus* | *Bisectae* | 1186 | |
| socialis_viridans_CCA3001 | DN 1946 | *Symphyomyrtus* | *Bisectae* | 1187 | |
| sp_CCA6677 | | | | 1187 | invalid CCA identifier |
| sp_Dartmoor_CCA5762 | DN 5585 | *Symphyomyrtus* | *Dumaria* | 1186 | technical replicates |
| sp_Dartmoor_CCA5764 | DN 5585 | *Symphyomyrtus* | *Dumaria* | 1186 | |
| sp_Lake_Magenta_CCA5749 | DN 5509 | *Symphyomyrtus* | *Dumaria* | 1186 | technical replicates |
| sp_Lake_Magenta_CCA5750 | DN 5509 | *Symphyomyrtus* | *Dumaria* | 1187 | |
| sparsa_CCA0572 | DN 427 | *Symphyomyrtus* | *Adnataria* | 1179 | |
| sparsifolia_CCA0790 | DN 629 | *Eucalyptus* | *Eucalyptus* | 1170 | putative contamination / mislabelling |
| spathulata_spathulata_CCA1248 | DN 234 | *Symphyomyrtus* | *Glandulosae* | 1186 | label fix (previous: CCA1233); technical replicates |
| spathulata_spathulata_CCA1250 | DN 234 | *Symphyomyrtus* | *Glandulosae* | 1187 | label fix (previous: CCA1231) |
| spectatrix_CCA3032 | DN 1751 | *Eucalyptus* | *Eucalyptus* | 1187 | |

| | | | | | |
|---|---|---|---|---|---|
| sphaerica_CCA6056 | DN 2437 | *Corymbia* | *Notiales* | 1166 | |
| sporadica_CCA0301 | DN 204 | *Symphyomyrtus* | *Glandulosae* | 1187 | |
| sporadica_CCA1210 | DN 1122 | *Symphyomyrtus* | *Glandulosae* | 1187 | label fix (previous: CCA1242) |
| sporadica_CCA5630 | DN 5468 | *Symphyomyrtus* | *Glandulosae* | 1187 | technical replicates |
| sporadica_CCA5631 | DN 5468 | *Symphyomyrtus* | *Glandulosae* | 1186 | technical replicates |
| sporadica_CCA5633 | DN 5468 | *Symphyomyrtus* | *Glandulosae* | 1187 | |
| spreta_CCA0146 | DN 122 | *Symphyomyrtus* | *Dumaria* | 1186 | |
| spreta_CCA0518 | DN 551 | *Symphyomyrtus* | *Dumaria* | 1187 | |
| squamosa_CCA3250 | DN 2072 | *Symphyomyrtus* | *Bisectae* | 1141 | |
| staeri_CCA1238 | DN 1133 | *Eucalyptus* | *Longistylus* | 1187 | label fix (previous: CCA1215) |
| steedmanii_CCA0032 | DN 317 | *Symphyomyrtus* | *Glandulosae* | 1187 | |
| stellulata_CCA0218 | DN 582 | *Eucalyptus* | *Eucalyptus* | 1143 | |
| stellulata_Little_Star_CCA5879 | | *Eucalyptus* | *Eucalyptus* | 1186 | |
| stoatei_CCA0565 | DN 181 | *Symphyomyrtus* | *Dumaria* | 1187 | |
| stowardii_CCA0386 | DN 281 | *Symphyomyrtus* | *Glandulosae* | 1186 | |
| stowardii_CCA0389 | DN 281 | *Symphyomyrtus* | *Glandulosae* | 1187 | technical replicates |
| striaticalyx_CCA1527 | DN 1548 | *Symphyomyrtus* | *Dumaria* | 1187 | |
| striaticalyx_CCA5832 | DN 5592 | *Symphyomyrtus* | *Dumaria* | 1186 | |
| stricklandii_CCA0654 | DN 344 | *Symphyomyrtus* | *Glandulosae* | 1187 | |
| stricta_CCA5562 | DN 5252 | *Eucalyptus* | *Eucalyptus* | 1187 | |
| strzeleckii_CCA3051 | DN 1725 | *Symphyomyrtus* | *Maidenaria* | 1182 | |
| subangusta_cerina_CCA0182 | DN 301 | *Symphyomyrtus* | *Glandulosae* | 1186 | |
| subangusta_pusilla_CCA0179 | DN 278 | *Symphyomyrtus* | *Glandulosae* | 1187 | |
| subangusta_subangusta_CCA0315 | DN 290 | *Symphyomyrtus* | *Glandulosae* | 1186 | |
| subtilis_CCA0339 | DN 140 | *Symphyomyrtus* | *Glandulosae* | 1186 | |

| | | | | | |
|---|---|---|---|---|---|
| subtilis_CCA0342 | DN 187 | *Symphyomyrtus* | *Glandulosae* | 1187 | technical replicates |
| subtilis_CCA0345 | DN 187 | *Symphyomyrtus* | *Glandulosae* | 1187 | |
| suffulgens_CCA1565 | DN 1267 | *Symphyomyrtus* | *Adnataria* | 1184 | |
| suggrandis_promiscua_CCA0023 | DN 311 | *Symphyomyrtus* | *Glandulosae* | 1185 | |
| suggrandis_suggrandis_CCA0221 | DN 197 | *Symphyomyrtus* | *Glandulosae* | 1186 | |
| suggrandis_suggrandis_CCA1153 | DN 1128 | *Symphyomyrtus* | *Glandulosae* | 1186 | label fix (previous: CCA1318) |
| sweedmaniana_sweedmaniana_CCA5643 | DN 5487 | *Symphyomyrtus* | *Dumaria* | 1187 | |
| synandra_CCA0253 | DN 297 | *Symphyomyrtus* | *Bisectae* | 1187 | technical replicates |
| synandra_CCA0254 | DN 297 | *Symphyomyrtus* | *Bisectae* | 1187 | |
| synandra_CCA5822 | DN 5588 | *Symphyomyrtus* | *Bisectae* | 1186 | |
| synandra_CCA5823 | DN 5588 | *Symphyomyrtus* | *Bisectae* | 1186 | technical replicates |
| talyuberlup_CCA0597 | DN 228 | *Symphyomyrtus* | *Glandulosae* | 1185 | label fix (previous: CCA0598) |
| tenuipes_CCA4770 | DN 6475 | *Cuboidea* | | 1097 | |
| tenuipes_Ferguson2024 | | *Cuboidea* | | 1182 | not CCA |
| tephroclada_CCA0365 | DN 314 | *Symphyomyrtus* | *Glandulosae* | 1186 | |
| terebra_CCA0128 | DN 117 | *Symphyomyrtus* | *Glandulosae* | 1186 | |
| tereticornis_rotunda_CCA1581 | DN 1272 | *Symphyomyrtus* | *Exsertaria* | 1186 | putative contamination / mislabelling |
| tereticornis_rotunda_CCA1583 | DN 1272 | *Symphyomyrtus* | *Exsertaria* | 1187 | putative contamination / mislabelling |
| tereticornis_tereticornis_CCA1386 | CSIRO 13398 | *Symphyomyrtus* | *Exsertaria* | 1185 | |
| tereticornis_tereticornis_CCA1389 | CSIRO 13398 | *Symphyomyrtus* | *Exsertaria* | 1186 | technical replicates |
| tereticornis_tereticornis_CCA1393 | CSIRO 13661 | *Symphyomyrtus* | *Exsertaria* | 1187 | |
| terminalis_CCA1399 | DN 2120 | *Corymbia* | *Notiales* | 1165 | |
| tessellaris_CCA5926 | | *Blakella* | *Abbreviatae* | 1107 | technical replicates |
| tessellaris_CCA5927 | | *Blakella* | *Abbreviatae* | 1159 | |
| tessellaris_CCA5928 | | *Blakella* | *Abbreviatae* | 1158 | technical replicates |

| | | | | | |
|---|---|---|---|---|---|
| tessellaris_CCA5929 | | *Blakella* | *Abbreviatae* | 1153 | technical replicates |
| tetrapleura_CCA1653 | DN 1242 | *Symphyomyrtus* | *Adnataria* | 999 | |
| tholiformis_CCA1562 | DN 1279 | *Symphyomyrtus* | *Adnataria* | 1185 | |
| tholiformis_CCA1563 | DN 1279 | *Symphyomyrtus* | *Adnataria* | 1187 | technical replicates |
| thozetiana_CCA1574 | DN 1281 | *Symphyomyrtus* | *Adnataria* | 1186 | |
| tindaliae_CCA1695 | DN 1300 | *Eucalyptus* | *Eucalyptus* | 1187 | |
| torquata_CCA0593 | DN 135 | *Symphyomyrtus* | *Dumaria* | 20 | label fix (previous: CCA0597); low coverage |
| tortilis_CCA0119 | DN 130 | *Symphyomyrtus* | *Glandulosae* | 1186 | |
| trachyphloia_CCA0744 | DN 714 | *Corymbia* | *Notiales* | 1141 | technical replicates |
| trachyphloia_CCA1440 | DN 2084 | *Corymbia* | *Notiales* | 1165 | |
| trachyphloia_CCA5935 | DN 714 | *Corymbia* | *Notiales* | 1169 | technical replicates |
| trachyphloia_CCA5936 | DN 714 | *Corymbia* | *Notiales* | 1166 | |
| transcontinentalis_CCA0552 | DN 550 | *Symphyomyrtus* | *Bisectae* | 1185 | |
| tricarpa_CCA2978 | DN 1727 | *Symphyomyrtus* | *Adnataria* | 1187 | |
| trivalva_CCA0536 | DN 529 | *Symphyomyrtus* | *Glandulosae* | 1187 | |
| trivalva_CCA1471 | DN 1534 | *Symphyomyrtus* | *Glandulosae* | 1187 | |
| tumida_CCA0133 | DN 159 | *Symphyomyrtus* | *Glandulosae* | 1187 | |
| uncinata_uncinata_CCA1280 | DN 195 | *Symphyomyrtus* | *Bisectae* | 1185 | label fix (previous: CCA1181) |
| urna_urna_CCA1122 | DN 1087 | *Symphyomyrtus* | *Bisectae* | 1146 | |
| urna_urna_CCA2872 | DN 1817 | *Symphyomyrtus* | *Bisectae* | 1187 | |
| urna_urna_CCA3020 | DN 1838 | *Symphyomyrtus* | *Bisectae* | 1187 | |
| urna_xesta_CCA5785 | DN 5466 | *Symphyomyrtus* | *Bisectae* | 1180 | |
| urna_xesta_CCA5786 | DN 5466 | *Symphyomyrtus* | *Bisectae* | 1185 | technical replicates |
| utilis_CCA0588 | DN 215 | *Symphyomyrtus* | *Glandulosae* | 1185 | |
| utilis_CCA5709 | DN 5493 | *Symphyomyrtus* | *Glandulosae* | 1187 | |

| | | | | | |
|---|---|---|---|---|---|
| utilis_CCA5710 | DN 5493 | *Symphyomyrtus* | *Glandulosae* | 1187 | technical replicates |
| varia_CCA1458 | DN 163 | *Symphyomyrtus* | *Glandulosae* | 1186 | |
| varia_CCA1459 | DN 163 | *Symphyomyrtus* | *Glandulosae* | 1184 | technical replicates |
| variegata_CCA1612 | DN 1246 | *Blakella* | '*Maculatae*' | 1164 | |
| vegrandis_recondita_CCA2882 | DN 1621 | *Symphyomyrtus* | *Glandulosae* | 1186 | |
| vegrandis_recondita_CCA2891 | DN 1883 | *Symphyomyrtus* | *Glandulosae* | 1186 | |
| vegrandis_vegrandis_CCA0076 | DN 207 | *Symphyomyrtus* | *Glandulosae* | 1186 | |
| vegrandis_vegrandis_CCA0236 | DN 201 | *Symphyomyrtus* | *Glandulosae* | 1186 | |
| victrix_CCA0005 | | *Symphyomyrtus* | *Adnataria* | 1185 | |
| victrix_CCA5829 | DN 5568 | *Symphyomyrtus* | *Adnataria* | 1184 | |
| victrix_CCA5831 | DN 5568 | *Symphyomyrtus* | *Adnataria* | 1185 | technical replicates |
| victrix_Ferguson2024 | | *Symphyomyrtus* | *Adnataria* | 1184 | not CCA |
| viminalis_cygnetensis_CCA1366 | DN 941 | *Symphyomyrtus* | *Maidenaria* | 1182 | technical replicates |
| viminalis_cygnetensis_CCA1368 | DN 941 | *Symphyomyrtus* | *Maidenaria* | 1185 | |
| viminalis_Ferguson2024 | | *Symphyomyrtus* | *Maidenaria* | 1187 | not CCA |
| viminalis_hentyensis_CCA5899 | DN 6961 | *Symphyomyrtus* | *Maidenaria* | 1186 | |
| viminalis_hentyensis_CCA5900 | DN 6961 | *Symphyomyrtus* | *Maidenaria* | 1186 | technical replicates |
| viminalis_viminalis_CCA5060 | DN 6158 | *Symphyomyrtus* | *Maidenaria* | 1173 | |
| virella_CCA1265 | DN 286 | *Symphyomyrtus* | *Adnataria* | 1185 | label fix (previous: CCA1218) |
| virginea_CCA5648 | DN 5536 | *Symphyomyrtus* | *Bisectae* | 1183 | putative contamination / mislabelling |
| virginea_Ferguson2024 | | *Symphyomyrtus* | *Bisectae* | 1186 | not CCA |
| viridis_aenea_CCA1437 | DN 2083 | *Symphyomyrtus* | *Adnataria* | 1187 | |
| viridis_viridis_CCA0728 | DN 729 | *Symphyomyrtus* | *Adnataria* | 1164 | |
| vokesensis_CCA1491 | DN 1502 | *Symphyomyrtus* | *Bisectae* | 713 | |
| wandoo_pulverea_CCA0085 | DN 285 | *Symphyomyrtus* | *Glandulosae* | 1186 | technical replicates |

| | | | | | |
|---|---|---|---|---|---|
| wandoo_pulverea_CCA0088 | DN 285 | *Symphyomyrtus* | *Glandulosae* | 1187 | |
| wandoo_wandoo_CCA0346 | DN 230 | *Symphyomyrtus* | *Glandulosae* | 1185 | |
| websteriana_CCA0142 | DN 138 | *Symphyomyrtus* | *Bisectae* | 1185 | |
| websteriana_CCA0143 | DN 138 | *Symphyomyrtus* | *Bisectae* | 1187 | technical replicates |
| websteriana_CCA0326 | DN 331 | *Symphyomyrtus* | *Bisectae* | 1187 | |
| websteriana_CCA6085 | DN 6947 | *Symphyomyrtus* | *Bisectae* | 1186 | |
| websteriana_CCA6087 | DN 6947 | *Symphyomyrtus* | *Bisectae* | 1186 | technical replicates |
| woodwardii_CCA0577 | DN 340 | *Symphyomyrtus* | *Dumaria* | 1187 | label fix (previous: CCA0576) |
| woodwardii_CCA0580 | DN 340 | *Symphyomyrtus* | *Dumaria* | 1187 | technical replicates |
| woollsiana_CCA0733 | DN 681 | *Symphyomyrtus* | *Adnataria* | 1174 | |
| wyolensis_CCA0108 | DN 112 | *Symphyomyrtus* | *Bisectae* | 1185 | |
| x_alpina_CCA5943 | DN 7046 | *Eucalyptus* | *Eucalyptus* | 1187 | |
| x_alpina_CCA5945 | DN 7046 | *Eucalyptus* | *Eucalyptus* | 1187 | technical replicates |
| x_balanopelex_CCA0335 | DN 169 | *Symphyomyrtus* | *Bisectae* | 1186 | |
| x_brachyphylla_CCA0611 | DN 338 | *Symphyomyrtus* | *Glandulosae* | 1186 | |
| x_brachyphylla_CCA0612 | DN 338 | *Symphyomyrtus* | *Glandulosae* | 749 | technical replicates |
| x_bunyip_CCA5876 | DN 6782 | *Symphyomyrtus* | *Maidenaria* | 1187 | technical replicates |
| x_bunyip_CCA5877 | DN 6782 | *Symphyomyrtus* | *Maidenaria* | 1187 | |
| x_carolaniae_CCA5881 | DN 6787 | *Symphyomyrtus* | *Maidenaria* | 1187 | technical replicates |
| x_carolaniae_CCA5882 | DN 6787 | *Symphyomyrtus* | *Maidenaria* | 1187 | |
| x_hawkeri_CCA5966 | DN 6916 | *Symphyomyrtus* | *Adnataria* | 1187 | technical replicates |
| x_hawkeri_CCA5968 | DN 6916 | *Symphyomyrtus* | *Adnataria* | 1187 | |
| x_impensa_CCA1395 | DN 264 | *Symphyomyrtus* | *Bisectae* | 1182 | putative contamination / mislabelling |
| xanthonema_apposita_CCA0283 | DN 232 | *Symphyomyrtus* | *Glandulosae* | 1186 | |
| xanthonema_xanthonema_CCA0247 | DN 202 | *Symphyomyrtus* | *Glandulosae* | 1186 | |

| | | | | | |
|---|---|---|---|---|---|
| xerothermica_CCA0516 | DN 541 | *Symphyomyrtus* | *Adnataria* | 1187 | |
| xerothermica_CCA1199 | DN 1187 | *Symphyomyrtus* | *Adnataria* | 1184 | label fix (previous: CCA1206) |
| yalatensis_CCA1150 | DN 950 | *Symphyomyrtus* | *Bisectae* | 1186 | label fix (previous: CCA1319) |
| yilgarnensis_CCA0010 | | *Symphyomyrtus* | *Adnataria* | 1185 | technical replicates |
| yilgarnensis_CCA0011 | | *Symphyomyrtus* | *Adnataria* | 1187 | |
| yilgarnensis_CCA0012 | | *Symphyomyrtus* | *Adnataria* | 1161 | technical replicates |
| youngiana_CCA0628 | DN 523 | *Symphyomyrtus* | *Bisectae* | 1187 | |
| youngiana_CCA2842 | DN 1535 | *Symphyomyrtus* | *Bisectae* | 1187 | |
| yumbarrana_CCA2835 | DN 1808 | *Symphyomyrtus* | *Bisectae* | 1186 | |
| zopherophloia_CCA0379 | DN 267 | *Symphyomyrtus* | *Glandulosae* | 1187 | |

**Table S2**. Concordance factors of all taxonomic groups within the CBA clade on the species tree based on 1,187 BUSCO loci. Groups with only one described species are excluded. Entries in bold refer to clades of interest. Clade: taxonomic groups included; gCF: gene concordance factor; gN: number of decisive gene trees used in gCF calculation; sCF: site concordance factor; sN: number of decisive parsimony informative sites used in sCF calculation (rounded down); qCF: quartet concordance factor; qN: number of decisive quartet trees used in qCF calculation; bb: UFBoot support based on 1,000 replicates; pp: ASTRAL posterior probability. All CFs are rounded down for easier comparison.

| clade | gCF | gN | sCF | sN | qCF | qN | bb | pp |
|---|---|---|---|---|---|---|---|---|
| **CBA + *Eucalyptus s.s.*** | **99** | **1160** | **97** | **70640** | **99** | **1160** | **100** | **1.00** |
| subg. *Angophora* + subg. *Corymbia** | 28 | 1153 | 37 | 10258 | 38 | 1153 | 100 | 1.00 |
| **subg. *Angophora*** | **89** | **1151** | **85** | **12216** | **95** | **1151** | **100** | **1.00** |
| **subg. *Corymbia**** | **45** | **1151** | **63** | **9842** | **72** | **1151** | **100** | **1.00** |
| sect. *Notiales*** + sect. '*Calophyllae*' | 15 | 1164 | 37 | 7023 | 43 | 1164 | 100 | 1.00 |
| sect. *Notiales*** | 7 | 1144 | 37 | 5153 | 40 | 1144 | 100 | 1.00 |
| sect. '*Calophyllae*' | 13 | 1166 | 48 | 6548 | 49 | 1166 | 100 | 1.00 |
| **subg. *Blakella*** | **78** | **1158** | **75** | **12721** | **90** | **1158** | **100** | **1.00** |
| sect. *Abbreviatae* | 56 | 1156 | 61 | 7787 | 68 | 1156 | 100 | 1.00 |
| sect. '*Maculatae*' + sect. '*Naviculares*' | 16 | 1161 | 37 | 6972 | 36 | 1161 | 99 | 0.90 |
| sect. '*Maculatae*' | 55 | 1162 | 73 | 7669 | 79 | 1162 | 100 | 1.00 |
| sect. '*Naviculares*' | 26 | 1159 | 54 | 6760 | 56 | 1159 | 100 | 1.00 |

* including *E. trachyphloia* (CCA1440)

** excluding *E. trachyphloia* (CCA1440)

**Table S3**. Concordance factors of all taxonomic groups within the subgenus *Eudesmia* on the species tree based on 1,187 BUSCO loci. Groups with only one described species are excluded. Entries in bold refer to clades of interest. Clade: taxonomic groups included; gCF: gene concordance factor; gN: number of decisive gene trees used in gCF calculation; sCF: site concordance factor; sN: number of decisive parsimony informative sites used in sCF calculation (rounded down); qCF: quartet concordance factor; qN: number of decisive quartet trees used in qCF calculation; bb: UFBoot support based on 1,000 replicates; pp: ASTRAL posterior probability. All CFs are rounded down for easier comparison.

| clade | gCF | gN | sCF | sN | qCF | qN | bb | pp |
|---|---|---|---|---|---|---|---|---|
| **subg. *Eudesmia*** | **64** | **1124** | **61** | **12221** | **84** | **1124** | **100** | **1.00** |

| | | | | | | | | |
|---|---|---|---|---|---|---|---|---|
| sect. *Limbatae* + sect. *Reticulatae* | 14 | 1124 | 35 | 6679 | 36 | 1124 | 100 | 0.96 |
| sect. *Limbatae* | 20 | 1182 | 47 | 6481 | 56 | 1182 | 100 | 1.00 |
| sect. *Reticulatae* | 44 | 1118 | 56 | 6826 | 59 | 1118 | 100 | 1.00 |

**Table S4**. Concordance factors of all taxonomic groups within the subgenus *Eucalyptus* on the species tree based on 1,187 BUSCO loci. Groups with only one described species are excluded. Entries in bold refer to clades of interest. Clade: taxonomic groups included; gCF: gene concordance factor; gN: number of decisive gene trees used in gCF calculation; sCF: site concordance factor; sN: number of decisive parsimony informative sites used in sCF calculation (rounded down); qCF: quartet concordance factor; qN: number of decisive quartet trees used in qCF calculation; bb: UFBoot support based on 1,000 replicates; pp: ASTRAL posterior probability. All CFs are rounded down for easier comparison.

| clade | gCF | gN | sCF | sN | qCF | qN | bb | pp |
|---|---|---|---|---|---|---|---|---|
| **subg. *Idiogenes* + subg. *Eucalyptus*** | **50** | **1175** | **71** | **7949** | **82** | **1175** | **100** | **1.00** |
| subg. *Eucalyptus* | 12 | 1179 | 40 | 6710 | 41 | 1179 | 100 | 1.00 |
| sect. *Longistylus* | 2 | 1187 | 35 | 6323 | 35 | 1187 | 100 | 0.78 |
| (sect. *Eucalyptus* + sect. *Amentum*) + sect. *Frutices* | 4 | 1187 | 43 | 6471 | 45 | 1187 | 100 | 1.00 |
| sect. *Frutices* | 4 | 1187 | 45 | 6258 | 46 | 1187 | 100 | 1.00 |
| sect. *Eucalyptus* + sect. *Amentum* | 5 | 1187 | 43 | 5815 | 44 | 1187 | 100 | 1.00 |
| sect. *Amentum* | 22 | 1135 | 46 | 5420 | 43 | 1135 | 100 | 1.00 |
| sect. *Eucalyptus* | 9 | 1187 | 55 | 5550 | 55 | 1187 | 100 | 1.00 |

**Table S5**. Concordance factors of all taxonomic groups within the subgenus *Symphyomyrtus* on the species tree based on 1,187 BUSCO loci. Groups with only one described species are excluded. Entries in bold refer to clades of interest. Clade: taxonomic groups included; gCF: gene concordance factor; gN: number of decisive gene trees used in gCF calculation; sCF: site concordance factor; sN: number of decisive parsimony informative sites used in sCF calculation (rounded down); qCF: quartet concordance factor; qN: number of decisive quartet trees used in qCF calculation; bb: UFBoot support based on 1,000 replicates; pp: ASTRAL posterior probability. All CFs are rounded down for easier comparison.

| clade | gCF | gN | sCF | sN | qCF | qN | bb | pp |
|---|---|---|---|---|---|---|---|---|
| **subg. *Symphyomyrtus*** | **21** | **1187** | **53** | **8121** | **62** | **1187** | **100** | **1.00** |
| **MEL clade** | **33** | **1187** | **73** | **10150** | **83** | **1187** | **100** | **1.00** |

| | | | | | | | | |
|---|---|---|---|---|---|---|---|---|
| sect. *Pumilio* | 34 | 1187 | 67 | 6677 | 68 | 1187 | 100 | 1.00 |
| (sect. *Maidenaria* + sect. *Latoangulatae* + sect. *Racemus*) + (sect. *Exsertaria* + sect. *Incognitae*) | 3 | 1187 | 34 | 5601 | 35 | 1187 | 100 | 0.83 |
| sect. *Maidenaria* + (sect. *Latoangulatae* + sect. *Racemus*) | 1 | 1187 | 42 | 5405 | 43 | 1187 | 100 | 1.00 |
| sect. *Maidenaria* | 7 | 1187 | 53 | 5283 | 56 | 1187 | 100 | 1.00 |
| sect. *Latoangulatae* + sect. *Racemus* | 4 | 1051 | 42 | 4143 | 39 | 1051 | 100 | 1.00 |
| sect. *Latoangulatae* | 12 | 1043 | 45 | 3795 | 51 | 1043 | 100 | 1.00 |
| sect. *Exsertaria* + sect. *Incognitae* | 0 | 1187 | 39 | 5312 | 40 | 1187 | 100 | 1.00 |
| sect. *Exsertaria* | 0 | 1185 | 35 | 4537 | 33 | 1185 | 100 | 0.06 |
| sect. *Incognitae* | 13 | 1146 | 45 | 4951 | 45 | 1146 | 100 | 1.00 |
| sect. *Bisectae* + (sect. *Domesticae* + sect. *Sejunctae* + AGD clade) | 3 | 1187 | 38 | 8508 | 40 | 1187 | 100 | 1.00 |
| **sect. *Bisectae*** | **4** | **1187** | **45** | **8194** | **49** | **1187** | **100** | **1.00** |
| sect. *Domesticae* + (sect. *Sejunctae* + AGD clade) | 3 | 1185 | 40 | 8690 | 43 | 1185 | 100 | 1.00 |
| sect. *Sejunctae* + AGD clade | 5 | 1185 | 46 | 8509 | 48 | 1185 | 100 | 1.00 |
| **AGD clade** | **1** | **1187** | **34** | **7481** | **34** | **1187** | **100** | **0.40** |
| sect. *Adnataria* | 1 | 1187 | 41 | 7335 | 42 | 1187 | 100 | 1.00 |
| sect. *Platysperma* + (sect. *Glandulosae* + sect. *Bolites* + sect. *Dumaria*) | 0 | 1187 | 35 | 7635 | 36 | 1187 | 100 | 0.78 |
| sect. *Platysperma* | 21 | 1187 | 69 | 9476 | 66 | 1187 | 100 | 1.00 |
| (sect. *Glandulosae* + sect. *Bolites*) + sect. *Dumaria* | 0 | 1187 | 42 | 7211 | 42 | 1187 | 100 | 1.00 |
| sect. *Glandulosae* + sect. *Bolites* | 1 | 1187 | 41 | 7011 | 42 | 1187 | 100 | 1.00 |
| sect. *Dumaria* | 1 | 1187 | 49 | 7277 | 49 | 1187 | 100 | 1.00 |

### SUPPLEMENTARY FIGURES

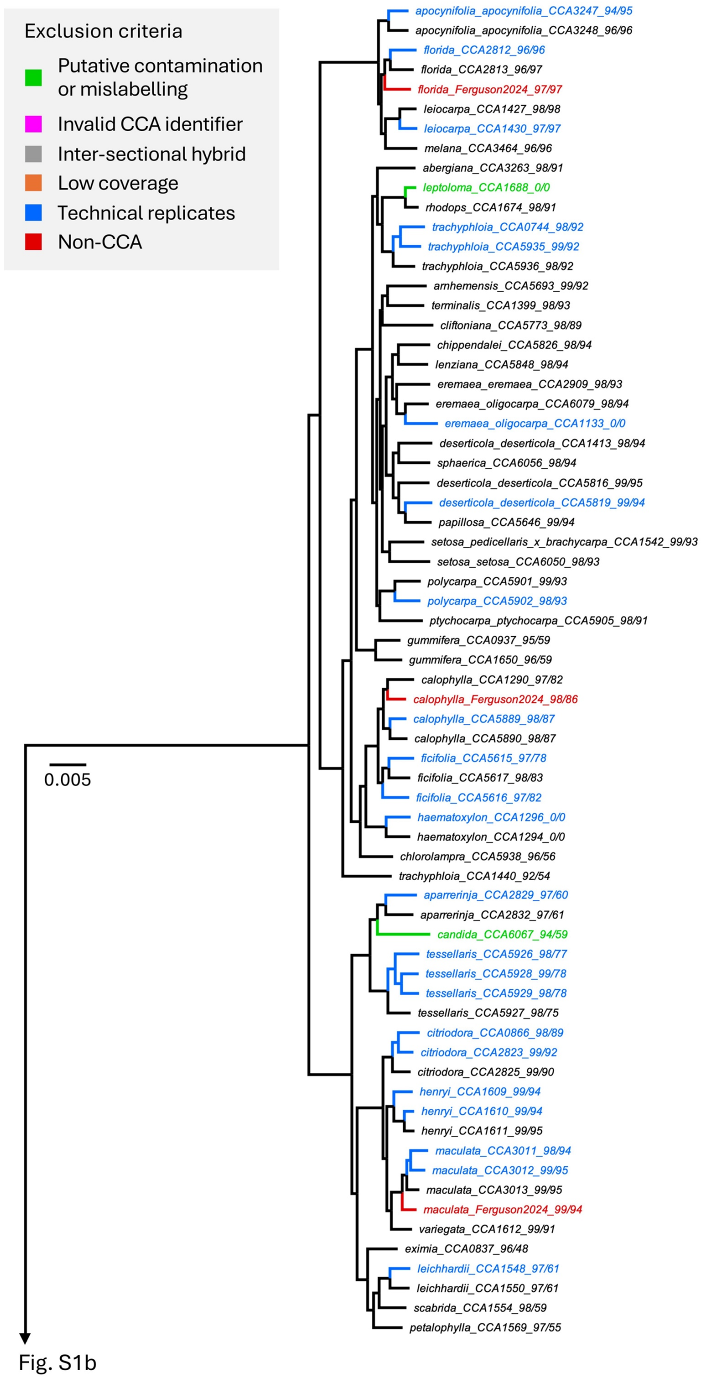


**Figure S1a**. Initial eucalypt concatenated tree inferred using IQ-TREE2 (part 1). The two numerical suffixes represent the subgenus/section consistency scores. Branch lengths are in the unit of substitution per site. Colouring denotes different filtering criteria (Table S1). Note that the taxonomic consistency scores were calculated prior to any label fixing.

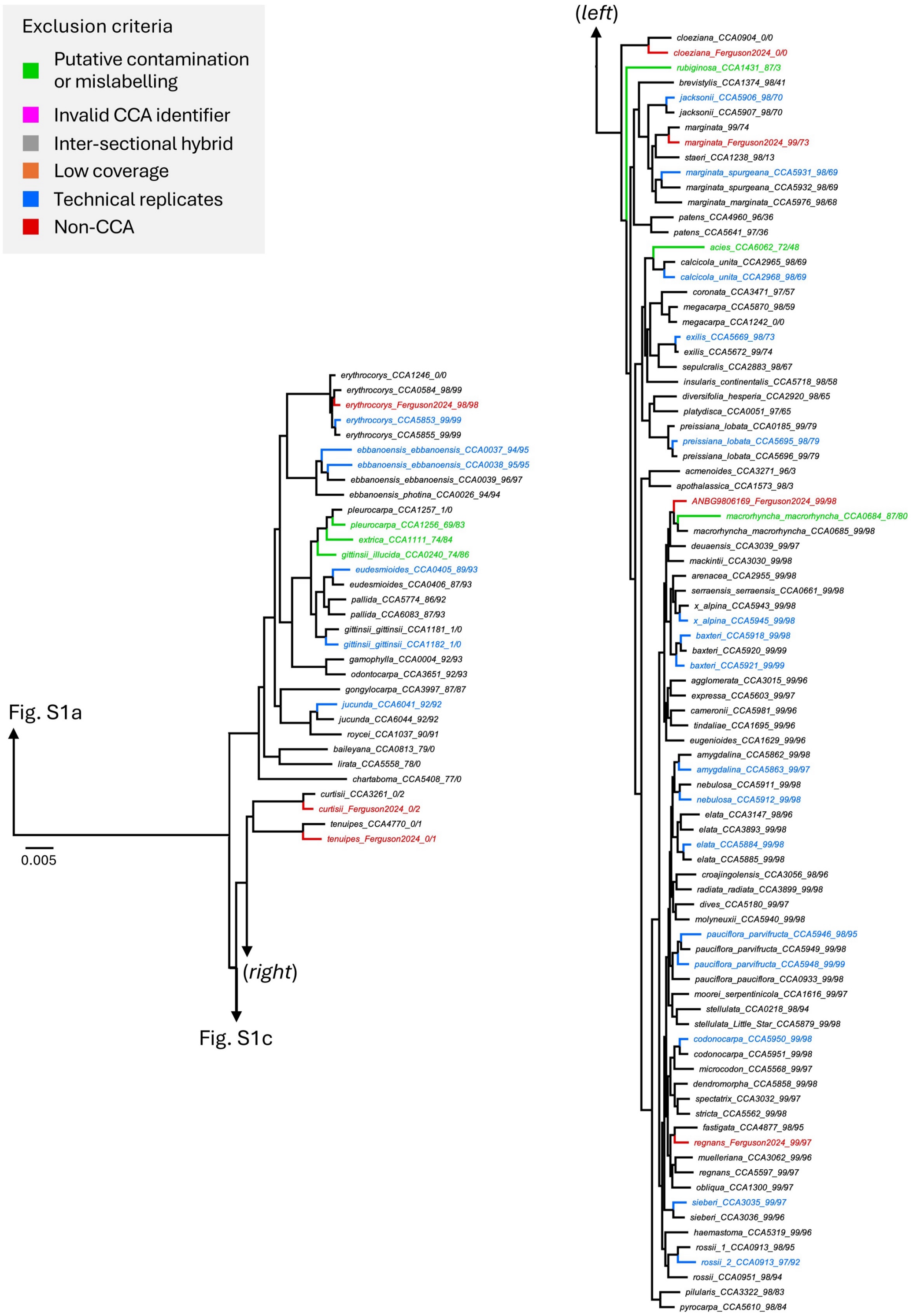


**Figure S1b**. Initial eucalypt concatenated tree inferred using IQ-TREE2 (part 2). The two numerical suffixes represent the subgenus/section consistency scores. Branch lengths are in the unit of substitution per site. Colouring denotes different filtering criteria (Table S1). Note that the taxonomic consistency scores were calculated prior to any label fixing.

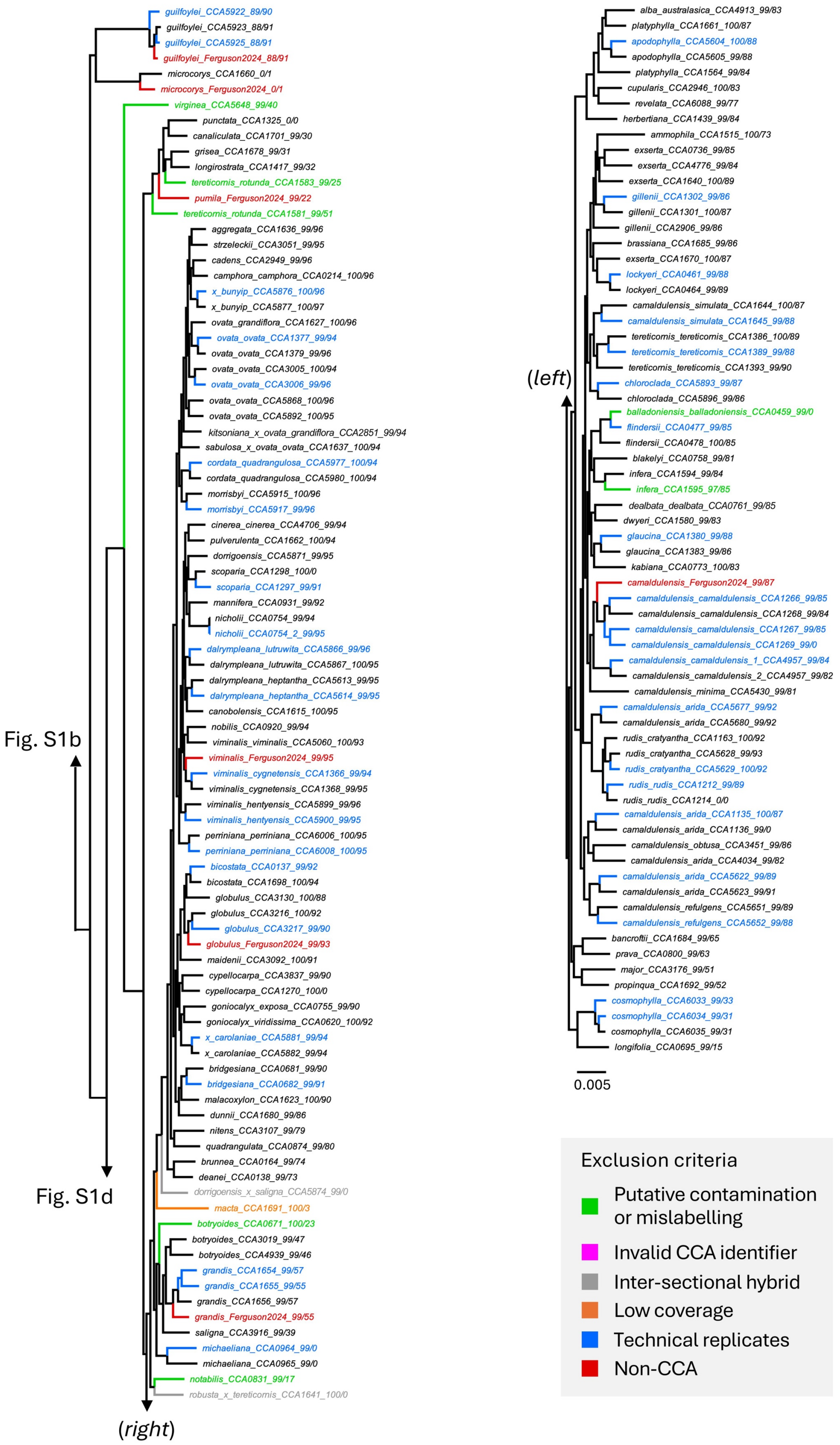


**Figure S1c**. Initial eucalypt concatenated tree inferred using IQ-TREE2 (part 3). The two numerical suffixes represent the subgenus/section consistency scores. Branch lengths are in the unit of substitution per site. Colouring denotes different filtering criteria (Table S1). Note that the taxonomic consistency scores were calculated prior to any label fixing.

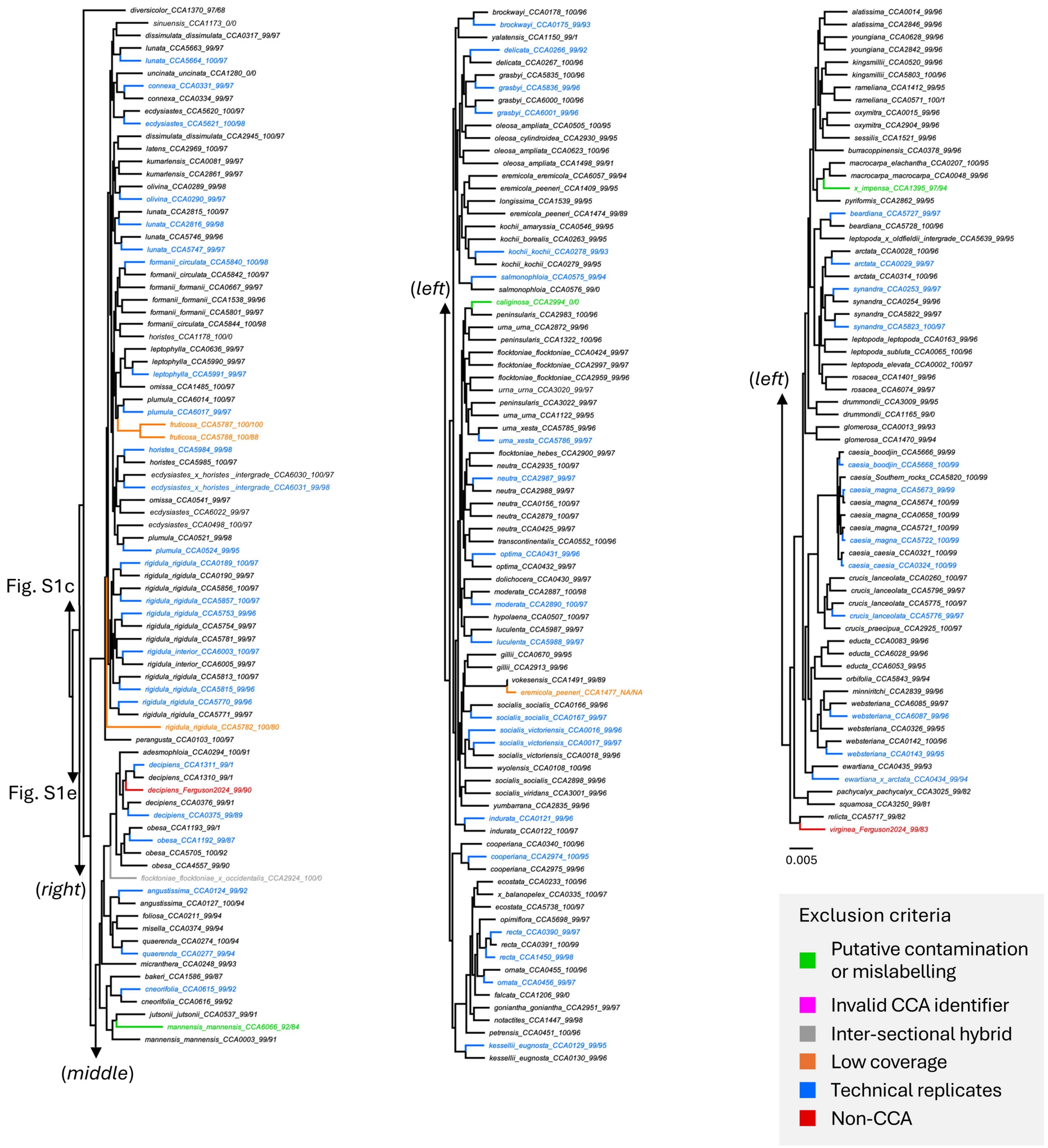


**Figure S1d**. Initial eucalypt concatenated tree inferred using IQ-TREE2 (part 4). The two numerical suffixes represent the subgenus/section consistency scores. Branch lengths are in the unit of substitution per site. Colouring denotes different filtering criteria (Table S1). Note that the taxonomic consistency scores were calculated prior to any label fixing.

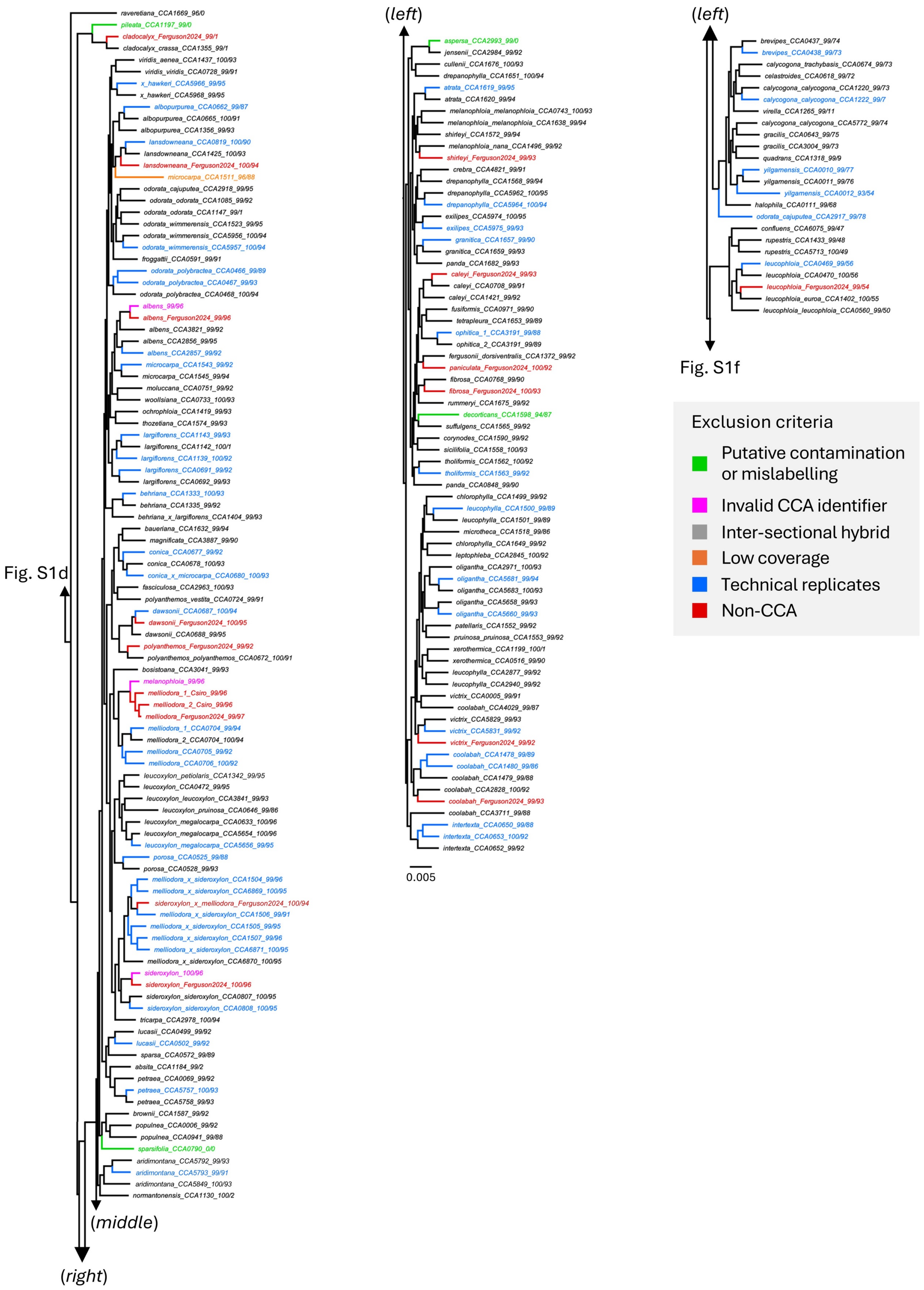


**Figure S1e**. Initial eucalypt concatenated tree inferred using IQ-TREE2 (part 5). The two numerical suffixes represent the subgenus/section consistency scores. Branch lengths are in the unit of substitution per site. Colouring denotes different filtering criteria (Table S1). Note that the taxonomic consistency scores were calculated prior to any label fixing.

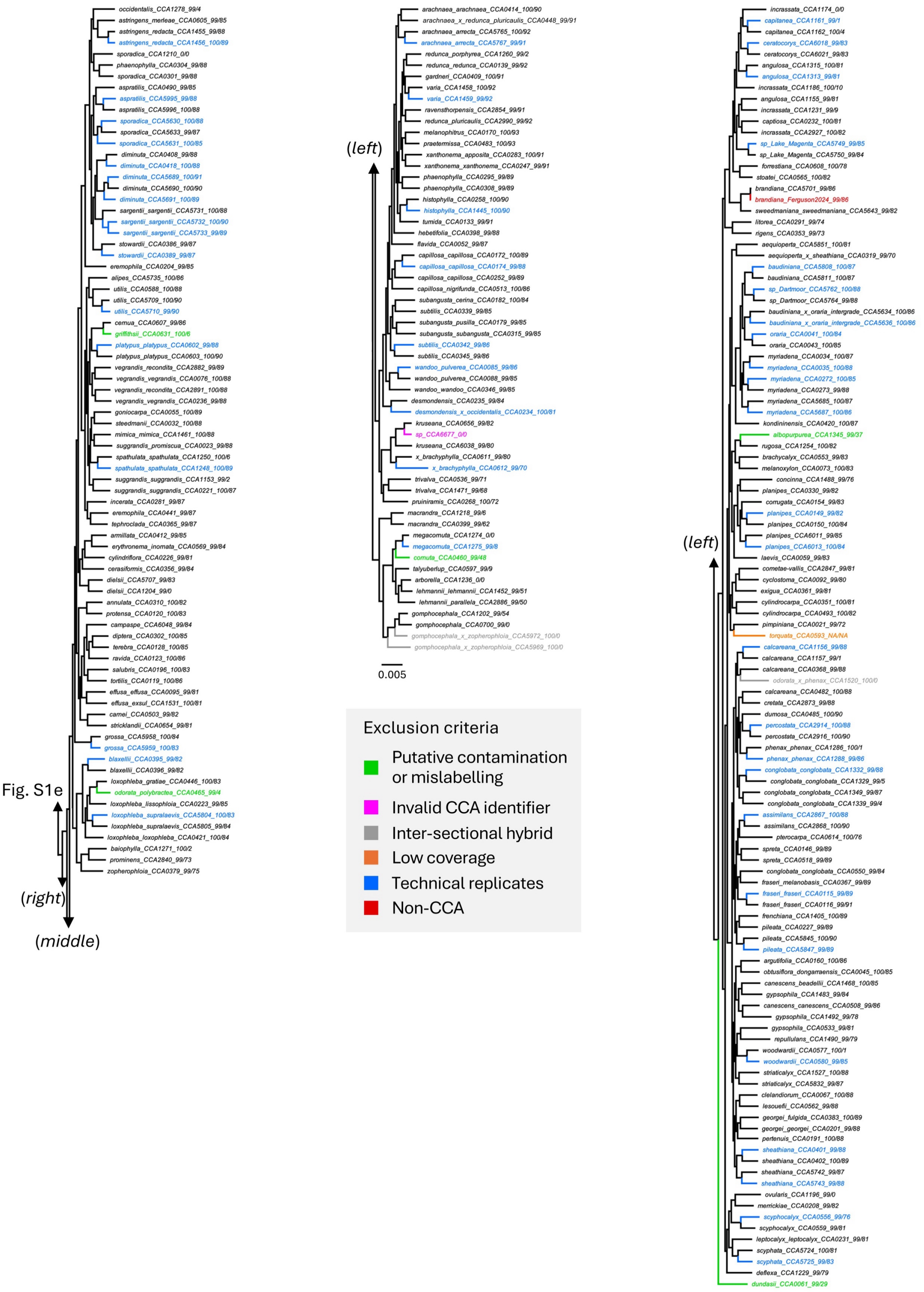


**Figure S1f**. Initial eucalypt concatenated tree inferred using IQ-TREE2 (part 6). The two numerical suffixes represent the subgenus/section consistency scores. Branch lengths are in the unit of substitution per site. Colouring denotes different filtering criteria (Table S1). Note that the taxonomic consistency scores were calculated prior to any label fixing.

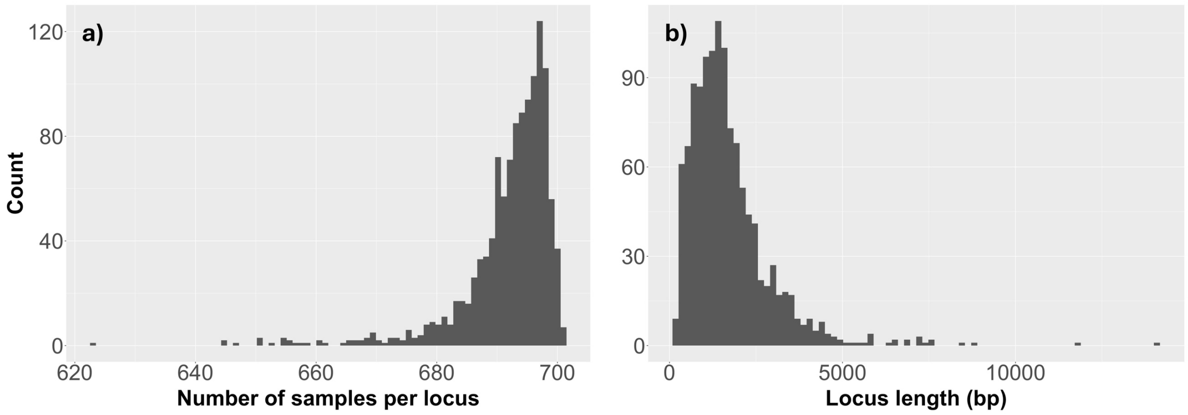


**Figure S2**. Distribution of (a) number of samples, and (b) locus length across 1,187 BUSCO loci comprising 701 samples. The x-axis is binned into 80 bins.

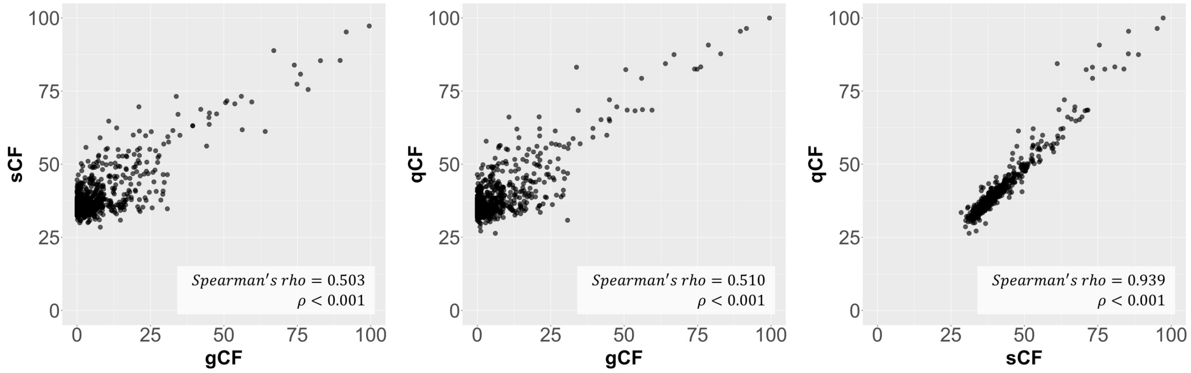


**Figure S3**. Scatter plots between gCF, sCF, and qCF on the species tree of 701 samples (Figs. 4-8). ρ reflects the approximate ρ-value of the Spearman's rank correlation analysis.

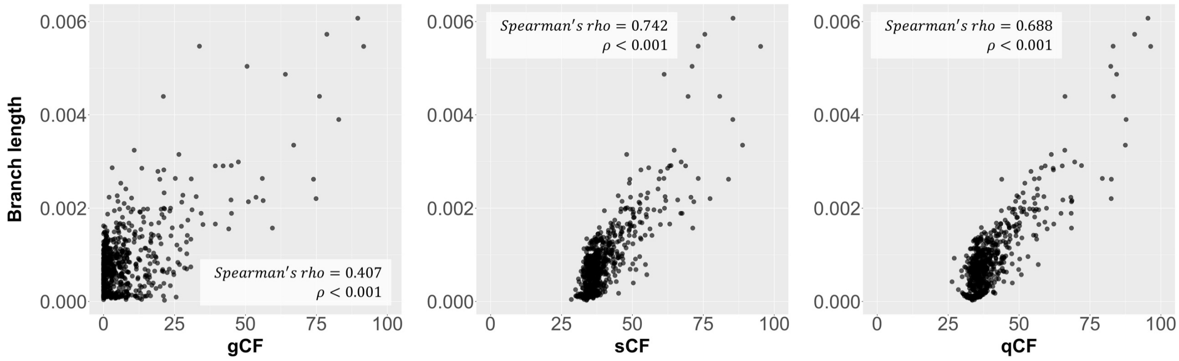


**Figure S4**. Scatter plots between concordance factors and branch lengths on the species tree of 701 samples (Figs. 4-8). The longest branch length that separates the CBA clade from the rest of eucalypts is excluded from the analysis. ρ reflects the approximate ρ-value of the Spearman's rank correlation analysis.

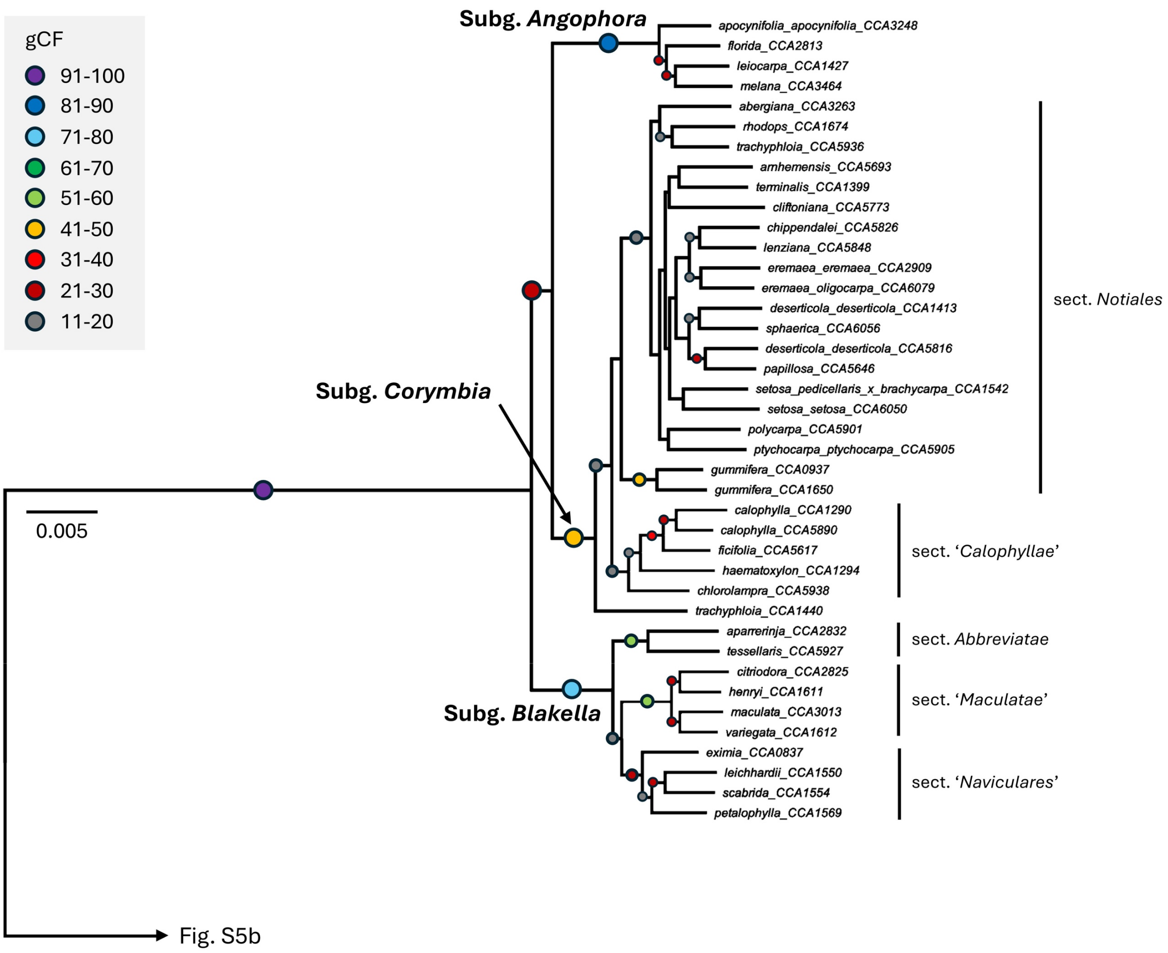


**Figure S5a.** Subtree of eucalypt species tree inferred using IQ-TREE2 (Fig. 3): subgenera *Angophora, Blakella,* and *Corymbia*. Branch lengths are in the unit of substitution per site. Dots represent gCF, with gCF < 11% not shown.

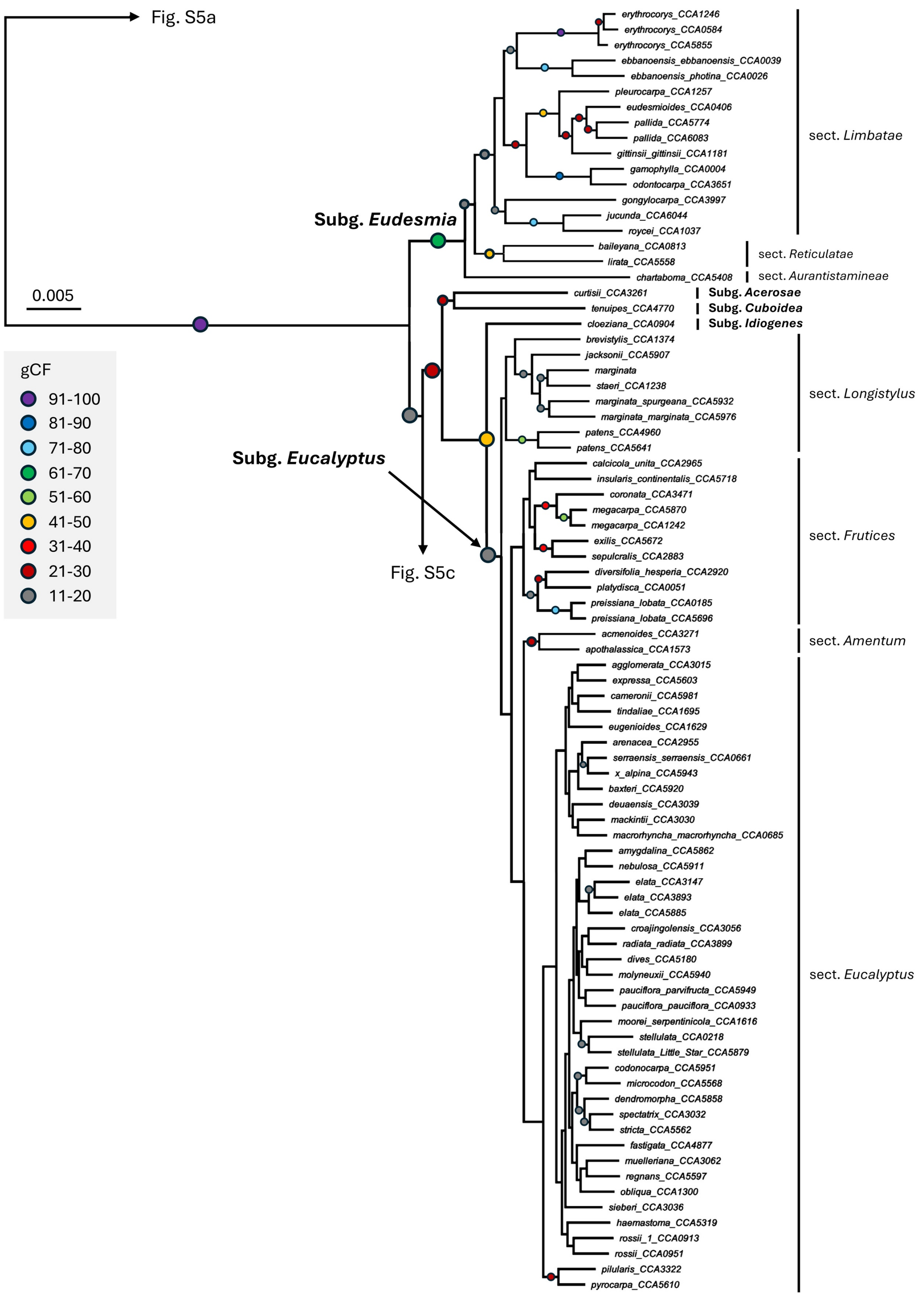


**Figure S5b.** Subtree of eucalypt species tree inferred using IQ-TREE2 (Fig. 3): subgenera *Eudesmia* and *Eucalyptus*. Branch lengths are in the unit of substitution per site. Dots represent gCF, with gCF < 11% not shown.

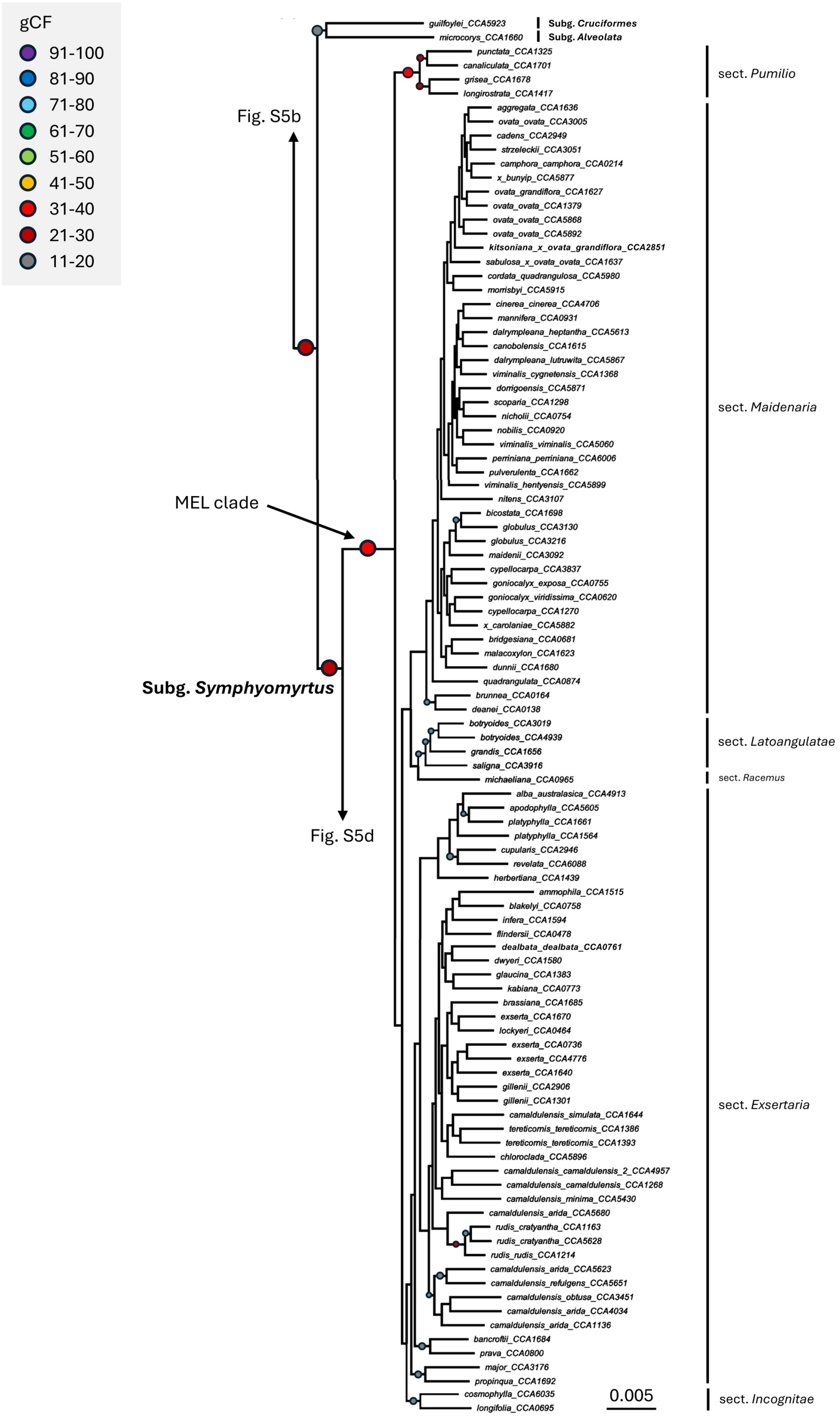


**Figure S5c.** Subtree of eucalypt species tree inferred using IQ-TREE2 (Fig. 3): subgenus *Symphyomyrtus* (MEL clade). Branch lengths are in the unit of substitution per site. Dots represent gCF, with gCF < 11% not shown.

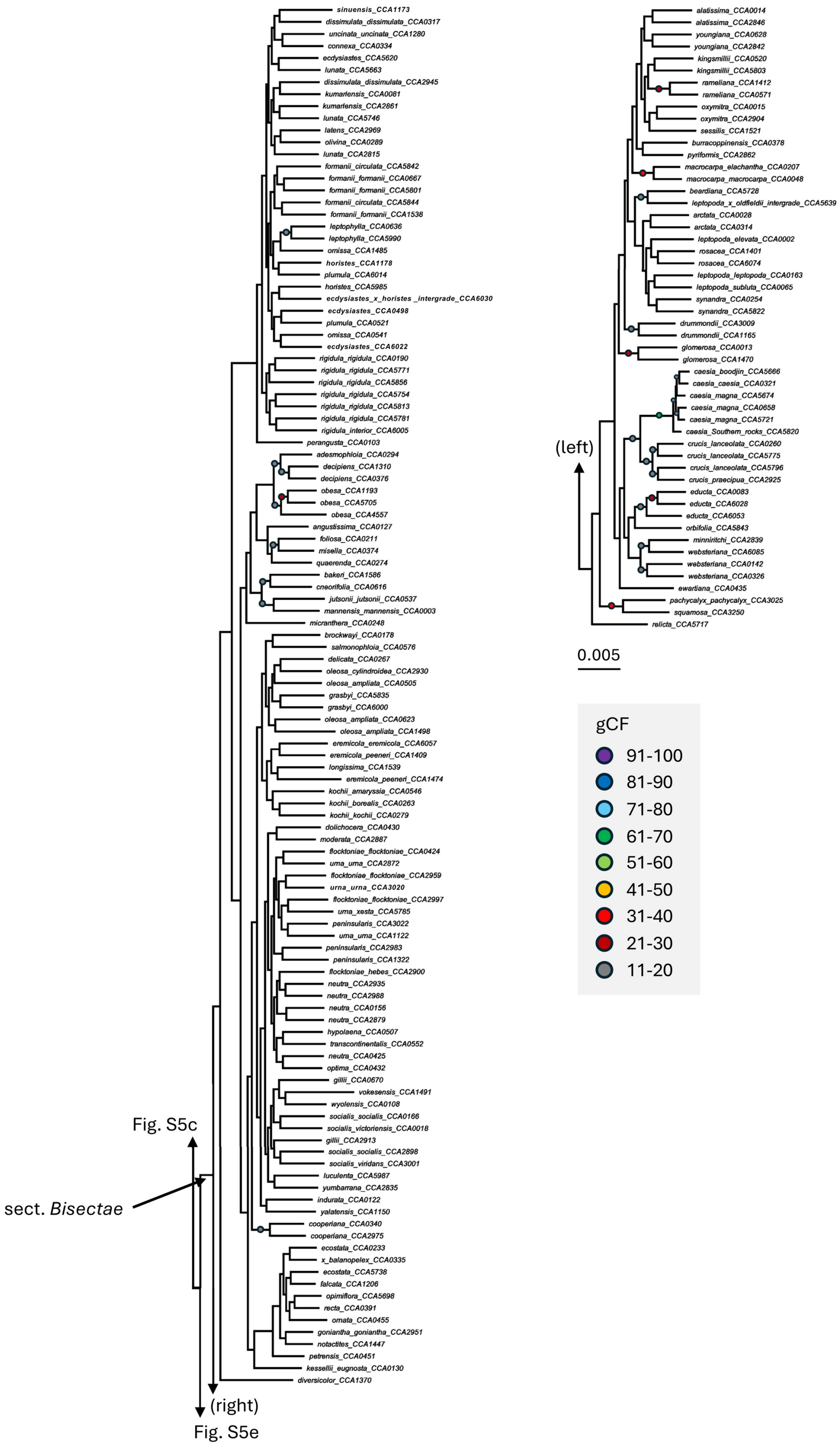


**Figure S5d.** Subtree of eucalypt species tree inferred using IQ-TREE2 (Fig. 3): subgenus *Symphyomyrtus* (*Bisectae* clade). Branch lengths are in the unit of substitution per site. Dots represent gCF, with gCF < 11% not shown.

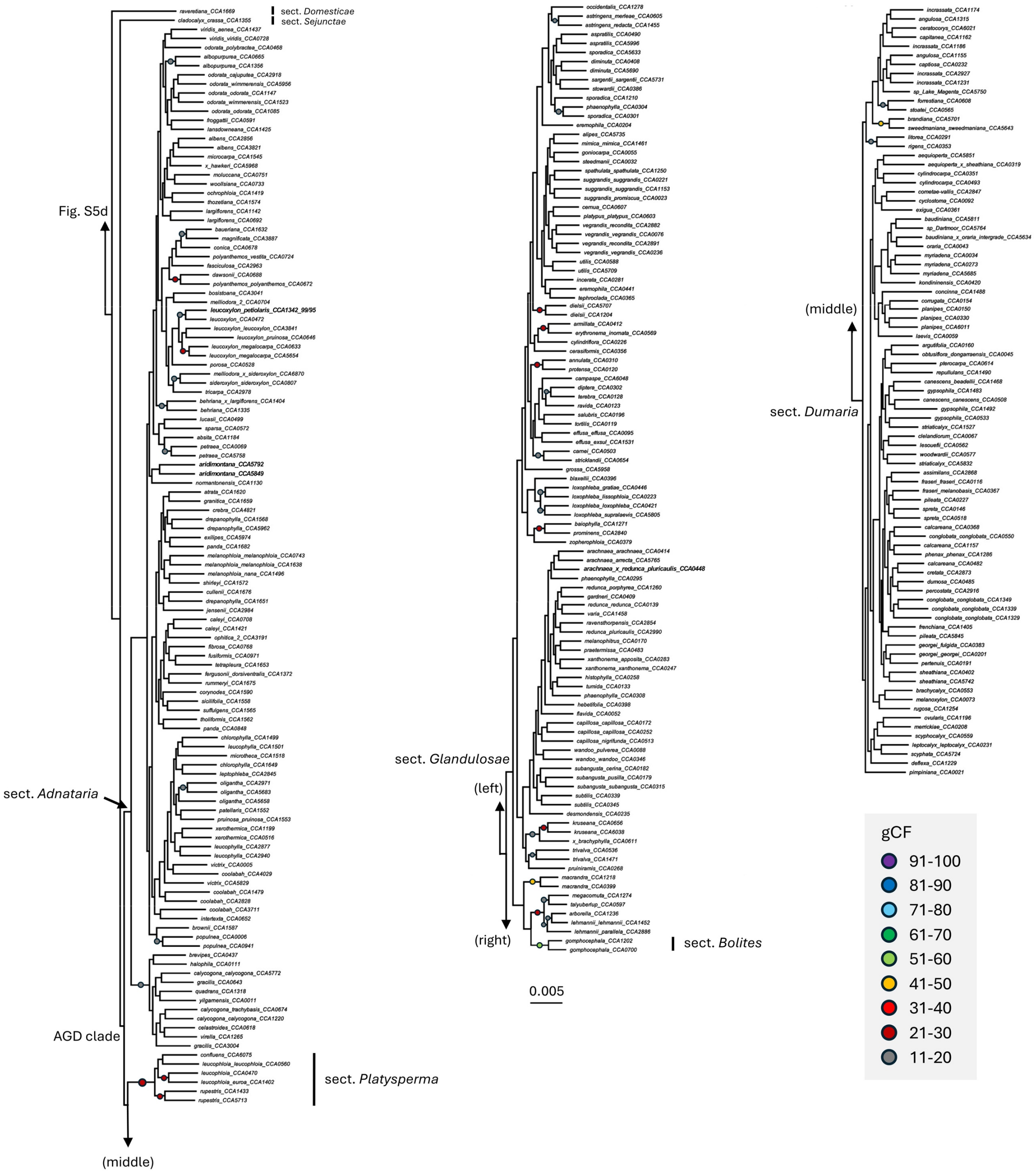


**Figure S5e.** Subtree of eucalypt species tree inferred using IQ-TREE2 (Fig. 3): subgenus *Symphyomyrtus* (AGD clade). Branch lengths are in the unit of substitution per site. Dots represent gCF, with gCF < 11% not shown.

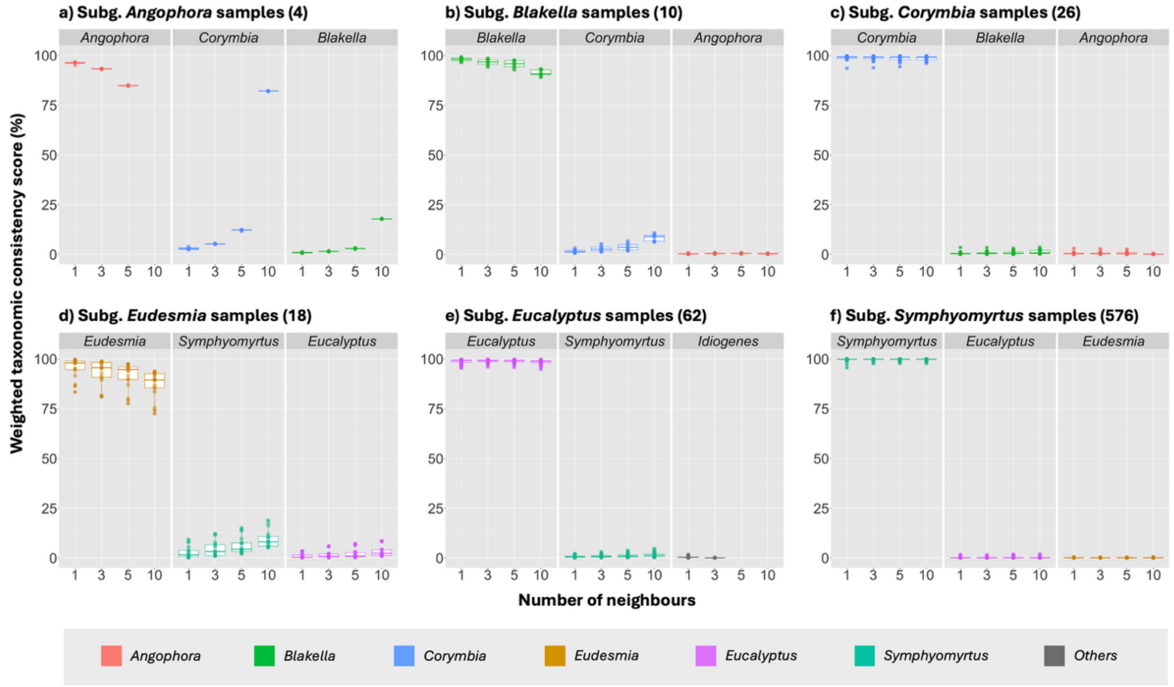


**Figure S6.** Subgenus consistency scores from 701 eucalypt samples across 1,187 gene trees based on one, three, five, and ten nearest neighbours on each gene tree. The Figure excludes subgenera with <3 representative samples. Each subtitle refers to the expected taxonomy of the samples, with numbers in parentheses representing the total number of samples. Each dot represents individual samples. Each panel shows three groups with the highest average count across different numbers of neighbours. Each colour denotes different subgenera.

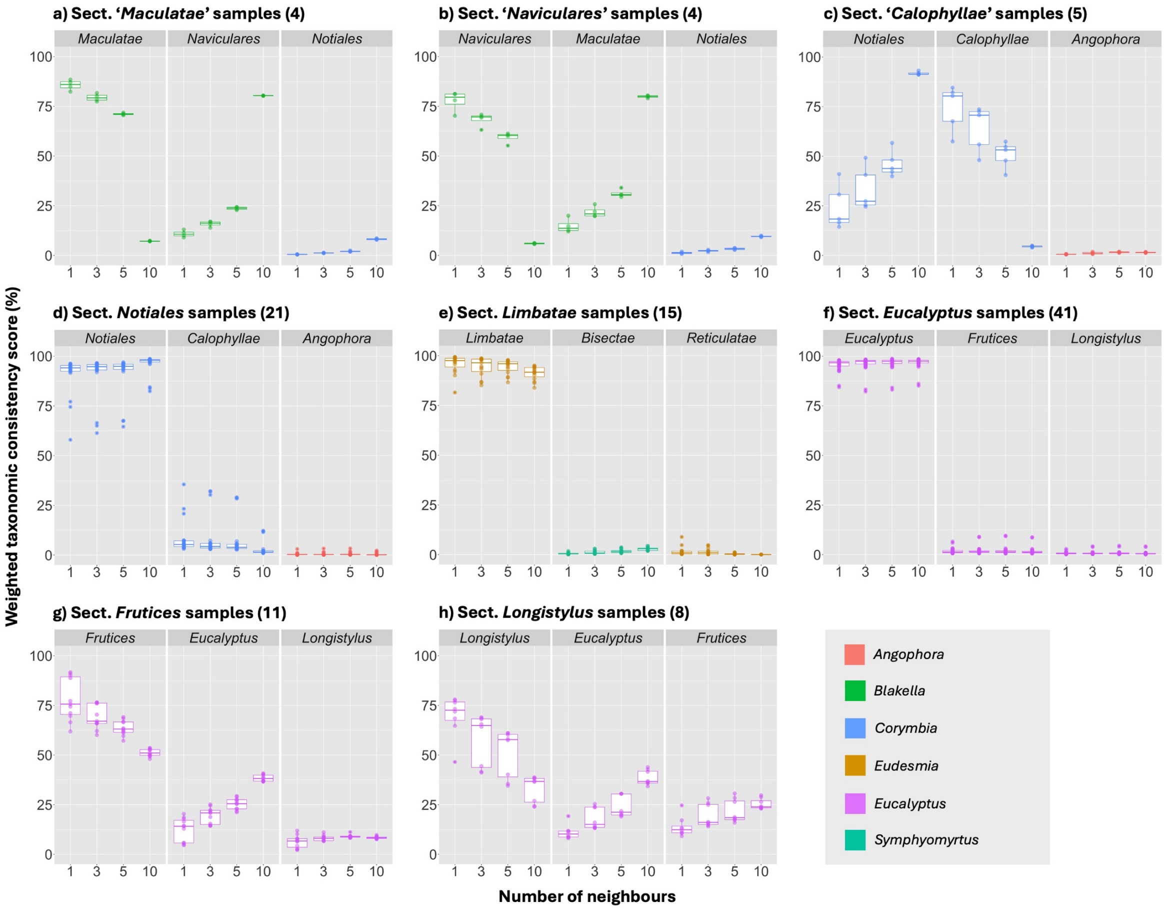


**Figure S7.** Section consistency scores (except for subg. *Symphyomyrtus*) from 701 eucalypt samples across 1,187 gene trees based on one, three, five, and ten nearest neighbours on each gene tree. The Figure excludes sections with <3 representative samples. Each subtitle refers to the expected taxonomy of the samples, with numbers in parentheses representing the total number of samples. Each dot represents individual samples. Each panel shows three groups with the highest average count across different numbers of neighbours. Each colour denotes different subgenera.

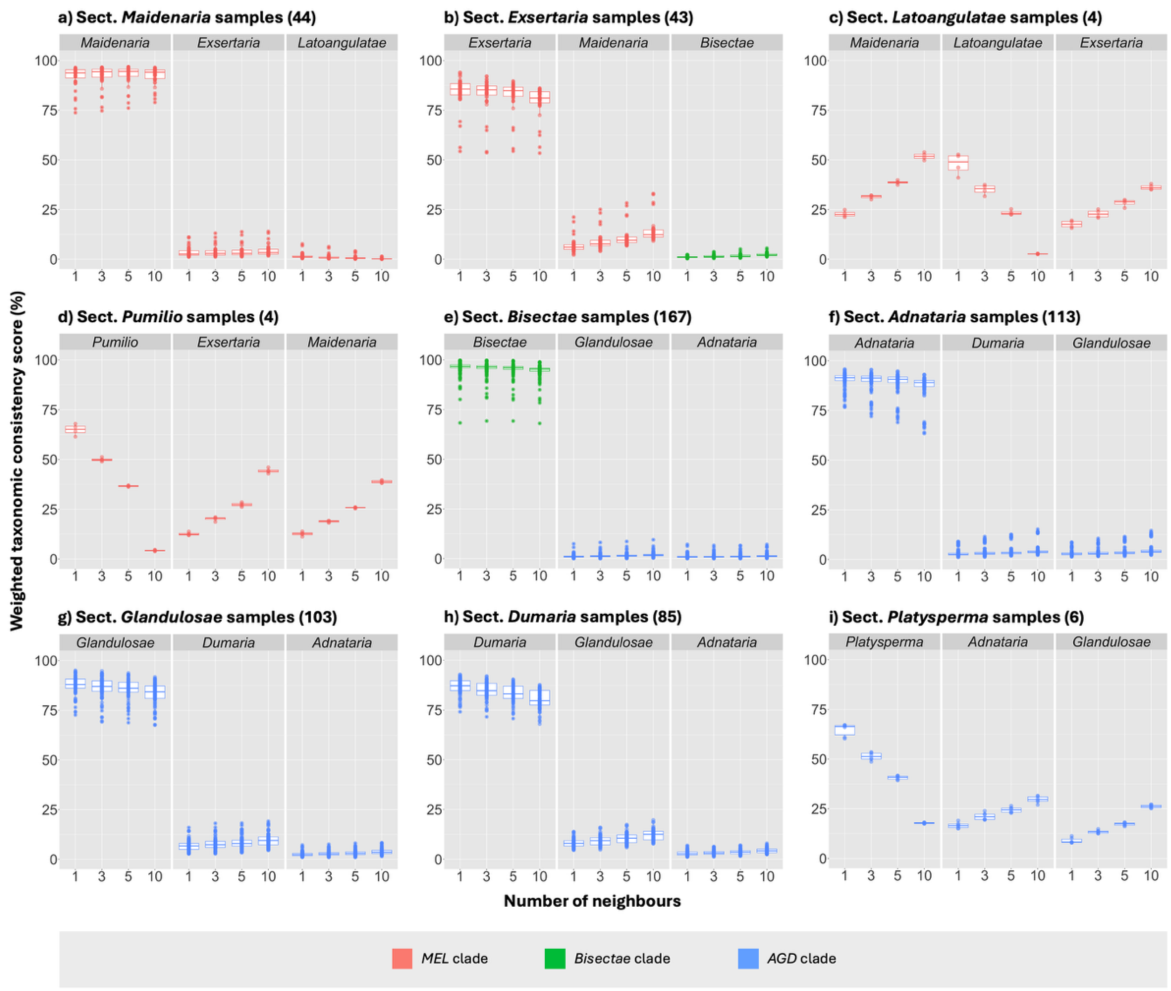


**Figure S8.** Section consistency scores of subg. *Symphyomyrtus* from 701 eucalypt samples across 1,187 gene trees based on one, three, five, and ten nearest neighbours on each gene tree. The Figure excludes sections with <3 representative samples. Each subtitle refers to the expected taxonomy of the samples, with numbers in parentheses representing the total number of samples. Each dot represents individual samples. Each panel shows three groups with the highest count across different numbers of neighbours. Each colour denotes different clades.
